# Early-onset β-amyloidosis in human brains with hematological malignances and cardiovascular diseases: Revisiting injury/stress induced axonal pathology

**DOI:** 10.1101/2025.08.16.670658

**Authors:** Yan Wang, Qi-Lei Zhang, Peng Zhou, Tian Tu, Zhong-Ping Sun, Xiao-Jie Zhang, Ewen Tu, Hui-Ping Chen, Hai-Ying Cheng, Aihua Pan, Jian Wang, Xiao-Xin Yan

**Affiliations:** Department of Anatomy and Neurobiology, Xiangya School of Basic Medical Sciences, Central South University, Changsha, Hunan 410013, China; Department of Pathology, Second Xiangya Hospital, Central South University, Changsha, Hunan 410031, China; Department of Neurology, Xiangya Hospital, Central South University, Changsha, Hunan 410008, China; Department of Psychiatry, Brain Bank for Psychiatric Disorders, The Second Xiangya Hospital of Central South University, Changsha Hunan 410011, China; Department of Neurology, The Second People’s Hospital of Hunan Province, Changsha, Hunan 410007, China; Department of Pathology, Hunan Guangxiu Hospital, Changsha, Hunan 410017, China; Reproductive and Stem Cell Engineering Institute, Xiangya School of Basic Medical Sciences, Central South University, Changsha, Hunan 410013, China

**Keywords:** Age-related dementia, Blood brain barrier, Neuritic pathology, Intraneuronal β-amyloid, White matter hyperintensity

## Abstract

β-Amyloid (Aβ) and tau pathologies are hallmarks of Alzheimer’s disease (AD) and they develop in human brain following differential spatiotemporal trajectories. As such, young/adult-onset tau-independent β-amyloidosis is rare. We encountered four such cases among 397 banked brains, with the donors died of hematological malignances (blood cancers) or cardiovascular diseases. To explore the pathological implications, we examined 17 brains (10-87 year-old, y) from blood cancer patients and three (52-82 y) with cardiovascular diseases, focusing on vascular injury, axonal pathology and Aβ formation. Aβ plaques occurred in two adult brains (31 y, 63 y) with blood cancers and two (52 y, 65 y) with cardiovascular diseases in the absence of tau. In the blood cancer brains, 17/17 had vascular injuries seen in hematoxylin-eosin stained sections, 13/17 had iron leakage, and 13/17 had axonal pathology. Malignant cell infiltration was found in 5/14 brains with myeloid, lymphocytic and lymphoma malignances, with light chain infiltration in 3/3 brains with multiple myeloma. In the cardiovascular disease brains, Aβ deposition primarily as diffuse plaques occurred in the cerebral cortex, with vascular and axonal pathologies in the white matter, striatum and internal capsule. Using a multi-labeling approach, the injury/stress induced axonal pathology was found to concur with β-amyloid processor protein elevation and enhanced β-secretase 1 processing but not intraneuronal Aβ accumulation. The current findings suggest that hematological malignances and cardiovascular diseases are risk conditions for early-onset cerebral β-amyloidosis, potentially attributable to vascular injury.

## Introduction

Extracellular β-amyloid peptide (Aβ) deposition and intraneuronal phosphorylated-tau (pTau) accumulation are hallmark neuropathologies of Alzheimer’s disease (AD), the most common cause of dementia among the elderly (Montine et al. 2012). Population-based prevalence studies suggest that cardiovascular diseases, diabetes, obesity and traumatic brain injury, which are often associated with cerebrovascular deficits, increase the risk of developing AD (Taudorf et al. 2019, GBD 2021 Nervous System Disorders Collaborators, 2024). Blood-brain barrier damages by periphery factors, such as microbial pathogens, environmental toxins and endogenous harmful metabolites, have been also proposed to play a role in AD pathogenesis (de la Torre 2010; Seaks and Wilcock, 2020; Chandra et al. 2024, Nihart et al. 2025).

Aβ and tau pathologies develop in the human brain in a spatiotemporally paradoxical manner, with Aβ plaque formation occurring later than pTau accumulation relative to age, initiated in the cerebral neocortex versus the limbic regions, respectively (Braak et al. 1991, Thal et al. 2002, Serrano-Pozo et al. 2011). Thus, primary age-related tauopathy (PART) refers to the occurrence of tau in the absence of Aβ lesion, which is very common among the elderly and can been seen in mid-age even younger individuals (Crary et al. 2014; Braak et al. 2025). In their seminal work staging Aβ and neurofibrillary tangle (NFT) pathogenesis in human brain, Braak and colleagues mentioned that one (Down syndrome, 17 year-old, y) out of 2332 brains exhibited Aβ plaques without tau pathology (Braak et al. 2006, Braak et al. 2011). In an early commentary article (Hyman and Gomez-Isla, 1997), it was noted that only 6 of 409 patients (1.5%) had cerebral Aβ without NFT at brain autopsy, although 115 (28.1%) cases had NFT but no Aβ. Further, in an original study including 105 autopsied brains from cognitively intact subjects (40–104 y), no case was found to have Aβ without NFT (Tsartsalis et al., 2018). Moreover, a recent review mentioned that only 12 (1.9%) in 628 brains collected in a hospital-banked cohort presented cerebral Aβ but lacking tau lesions (Thal et al. 2022).

Supported by a body-donation program for medical education and research, the Xiangya Human Brain bank has collected 397 whole postmortem brains during the past ten years from donors died of various diseases with death ages ranging from infancy to 103 y (Yan et al. 2015, Li et al. 2023, Wang et al. 2025). We encountered four brains with cerebral Aβ plaques in the absence of tauopathies from adult donors died of either hematological malignances (blood cancers, 31 y and 65 y) or cardiovascular diseases (52 y and 65 y). Pathological evaluation on such brain samples could help understand the risk factors and mechanism governing cerebral Aβ formation. In addition, neurological complications in patients with blood cancers have been increasingly emphasized clinically (Jurczyszyn et al., 2016; Paul et al. 2022, Mallio et al. 2023). Although central nervous system (CNS) involvement of blood cancers has been reported in early autopsy studies (Wolk et al. 1974; Jellinger et al. 1976; Barcos et al. 1987), few data are available for the presence of AD-type or related neuropathologies in this group of common malignancy. As blood cancers and cardiovascular diseases are inherently related to the circulation system, we suspected that vascular injury might be the key factor promoting the amyloidogenic cascade in the brain. Along this line of thinking, the current study examined 17 available brains obtained from children to old-age individuals died of blood cancers, and three additional brains from donors died of cardiovascular disorders that exhibited microscopically prominent Aβ deposition but no or only very early stage tau pathology, using a battery of histological and immunohistochemical labeling methods.

## Materials and Methods

### Human brain samples and tissue preparation

The present study was carried out in compliance with the Code of Ethics of the World Medical Association (Declaration of Helsinki) and the approval (2020KT-37, 4/10/2020; #2023-KT084, 6/21/2023) by the Ethics Committee of Central South University Xiangya School of Medicine. Brain banking was established via a willed body donation program (Yan et al. 2015). Basic histological assessment of Alzheimer’s and Parkinson disease neuropathologies were performed according to the Standard Brain Banking Protocol recommended by the China Brain Bank Consortium (Qiu et al. 2019). The brains used in the current study were selected based on the last hospitalization record. A total of 17 brains from donors died at 10-87 y with blood cancers were pathologically evaluated for the presence of Aβ deposition, tau and tangle lesions, amyloidogenic axonal pathology, vascular injury and brain infiltration of cancer markers. Three brains from donors died of acute cardiovascular conditions (52-82 y) were included in the present study following initial finding of microscopically overt cerebral Aβ deposition with no or mild neuronal tauopathy (Table 1). Briefly, each brain was bisected following removal from the skull. The right half-brain was sliced into 1 cm thickness and preserved fresh-frozen at –70 °C. The left-half brain was immersed in formalin for 2–3 weeks, which was then cut coronally into 1 cm thick slices. Tissue blocks covering major neuroanatomical structures were dissected out, including cerebral neocortical areas, the hippocampal formation, amygdala, striatum, thalamus and hypothalamus, midbrain, pons and medulla oblongata, and the cerebellar cortex and dentate nuclei. The brain blocks were prepared into cryostat (35 μm-thick) and paraffin (3 μm-thick) sections followed by histological and immunohistochemical preparations.

**Table 1.** Demographic information of brain donors and summary of histopathological findings.

| Brain coding | Clinical diagnosis | Age (y) | Sex | Survival time (m) | PMD (hr) | Braak NFT staging | Thal Aβ staging | Axonal pathology | Vascular injury | Iron leakage | Cancer marker infiltration |
| --- | --- | --- | --- | --- | --- | --- | --- | --- | --- | --- | --- |
| 1 | AML | 10 | M | 6 | 12 | 0 | 0 |  | (+) |  |  |
| 2 | AML | 17 | M | 20 | 5 | 0 | 0 |  | (+) |  |  |
| 3 | AML | 37 | F | 8 | 12 | 0 | 0 | (+) | (+) |  |  |
| 4 | AML | 14 | M | 4 | 9 | 0 | 0 |  | (+) | (+) |  |
| 5 | AML | 16 | M | 3 | 11 | 0 | 0 | (+) | (+) | (+) |  |
| 6 | AML | 50 | F | 6 | 8 | 0 | 0 | (+) | (+) |  |  |
| 7 | AML | 21 | M | 12 | 5 | 0 | 0 | (+) | (+) | (+) | CD68/MPO/Lys+ cells |
| 8 | AML | 16 | F | 4 | 6 | 0 | 0 | (+) | (+) | (+) | CD68/MPO/Lys+ cells |
| 9 | AML | 31 | F | 4 | 11 | 0 | 1 | (+) | (+) | (+) | CD68/MPO/Lys+ cells |
| 10 | ALL | 32 | F | 15 | 11 | 0 | 0 | (+) | (+) | (+) |  |
| 11 | ALL | 24 | M | 36 | 8 | 0 | 0 |  | (+) | (+) |  |
| 12 | MDS | 48 | M | 12 | 13 | 0 | 0 | (+) | (+) | (+) | CD68/Lys+ astrocytes |
| 13 | Lymphoma | 51 | M | 42 | 9 | 0 | 0 | (+) | (+) | (+) | CD20+ cells |
| 14 | Lymphoma | 87 | M | 24 | 6 | IV | 1 | (+) | (+) | (+) |  |
| 15 | MM | 62 | F | 16 | 4 | II | 0 | (+) | (+) | (+) | Light chain $\kappa$ |
| 16 | MM | 63 | F | 72 | 7 | 0 | 5 | (+) | (+) | (+) | Light chain $\lambda$ |
| 17 | MM (demented) | 74 | F | 36 | 5 | IV | 4 | (+) | (+) | (+) | Light chain $\kappa$ |
| 18 | AMI | 52 | M | N/A | 11 | 0 | 1 | (+) | (+) | (+) |  |
| 19 | CCI | 65 | M | 60 | 11 | 0 | 2 | (+) | (+) | (+) |  |
| 20 | Hypertension, CAD, SAH | 82 | F | 72 | 6 | 1a | 2 | (+) | (+) | (+) |  |

AML: acute myeloid leukemia; ALL: acute lymphocytic leukemia; MDS: Myelodysplastic syndrome; MM: multiple myeloma; AMI: acute myocardial infarction; CCI: chronic cerebral ischemia; CAD: coronary artery disease; SAH: subarachnoid hemorrhage. Survival time (months): duration between the first clinical diagnosis and death. "N/A": information not available as the patient died following the first hospitalization; PMD: postmortem delay in hours (hr); Braak staging of neurofibrillary tangle (NFT) pathology was based on pTau immunolabeling with the AT8 antibody and silver stains, Thal staging on Aβ immunolabeling with the 6E10 antibody; Axonal pathology was assessed with the 6E10, APP and BACE1 antibodies; Vascular injury was appraised with hematoxylin-eosin stain, and iron leakage with Prussian blue stain.

### Histological stains

Hematoxylin-eosin (HE) stain (HE Stain Kit, 60502ES76, Yeasen Biotechnology Co., Ltd., China) and Prussian Blue Iron Stain (Prussian Blue Stain Kit #60533ES20, Yeasen, China) were carried out using a Leica ST5010 Autostainer (Leica Biosystems, USA) according to manufacturer’s instructions. Bielschowsky and Gallyas silver stains were carried out according to physical development protocols (Shi et al. 2020; Zhang et al. 2024). Two fluorescent tracers were used in the present study to observe amyloid binding along with immunofluorescence (detailed below). The Amylo-Glo is a bright-blue fluorescent dye containing styrylbenzene similar to Thioflavin S/T, and can stains amyloid deposits as well as mature and ghost tangles (Yang et al. 2024). A ready-to-use kit of this reagent was obtained commercially and used according to manufacturer’s instruction (Cat#TR-300-AG, Biosensis Pty Ltd. SA5031, Australia). DANIR 8c is a highly potent and selective red fluorescent probe developed for Aβ imaging (Fu et al. 2016). Briefly, following double or single immunofluorescent antibody labeling, the paraffin sections were counterstained with Amylo-Glo or DANIR-8c and differentiated in 50% ethanol under microscopic monitoring (Zhang et al., 2024).

### Immunohistochemistry and immunofluorescence

Frozen and paraffin brain sections were stained immunohistochemically free-floatingly and on-slide, respectively, with the avidin-biotin complex (ABC) method, using one of the antibodies listed in Table 2. The immunolabeling protocol was detailed in our recent studies (Tu et al. 2020; Jiang et al. 2022; Yang et al. 2024). Briefly, the paraffin sections were dewaxed in xylene and rehydrated through descending ethanol solutions, followed by antigen retrieval in ethylenediaminetetraacetic acid (EDTA, 1 mM, pH 8) at 98 °C for 10 minutes. For Aβ immunolabeling, the sections were additionally treated with 90% formic acid at room temperature for 10 minutes. Cryostat and the above-treated paraffin sections were incubated in with 0.01M phosphate-buffered saline (PBS, pH7.3) containing a primary antibody, next with the pan-specific biotinylated secondary antibody (horse anti-mouse, rabbit and goat IgG, 1:200, BA-1300, Vector Laboratories, USA), and further with the avidin-biotin complex (ABC) (1:200, PK-6100, Vector Laboratories). Hematoxylin counterstain was applied on some immunolabeled sections for neuroanatomical orientation. The microslides were allowed for air-dry, dehydrated, cleared and mounted with glass coverslips.

**Table 2.** Primary antibodies used in the present study.

| Antibody name | Host species | Manufacturer | Product code | Dilution |
| --- | --- | --- | --- | --- |
| β-Amyloid (Aβ1-16) 6E10 | mouse | ThermoFisher | MA5-51794 | 1 to 2000 |
| β-Amyloid (Aβ17-24) 4G8 | mouse | Millipore | MFCD03457546 | 1 to 2000 |
| β-Amyloid (Aβ near NT) D54D2 | rabbit | Cell Signaling | 38424SF | 1 to 1000 |
| β-Amyloid (Aβ38-42) Ter42 | rabbit | Dr. Mori | Cai et al, 2012 | 1 to 1000 |
| β-Amyloid precursor protein (APP 750-CT) Y188 | rabbit | Abcam | Y188 | 1 to 2000 |
| β-Amyloid precursor protein (APP NT66-81) 22C11 | mouse | Merck | MAB348 | 1 to 2000 |
| β-Secretase 1 (BACE1) | rabbit | Abcam | EPR19523 | 1 to 2000 |
| Phospho-Tau (Ser202, Thr205, pTau) AT8 | mouse | ThermoFisher | MN1020 | 1 to 4000 |
| CD68 | rabbit | Sigma-Aldrich | HPA048982 | 1 to 500 |
| CD138 | rabbit | Sigma-Aldrich | HPA006185 | 1 to 500 |
| CD20 | mouse | Thermo Fisher | 14-0202-82 | 1 to 500 |
| Myeloperoxidase (MPO) | rabbit | Invitrogen | PA5-16672 | 1 to 500 |
| Lysozyme (Lys) | rabbit | Thermo Fisher | PA1-29680 | 1 to 500 |
| Multiple myeloma oncogene 1 (MUM1) | rabbit | Abcam | ab247079 | 1 to 500 |
| Immunoglobulin light chain $\kappa$ | mouse | Abcam | ab124727 | 1 to 500 |
| Immunoglobulin light chain $\lambda$ | mouse | Abcam | ab218430 | 1 to 500 |

Fluorescent labeling experiments were carried out in paraffin sections from selected brains for mouse anti-Aβ 6E10 and rabbit anti-BACE1 immunolabeling with 4’,6-diamidino-2-phenylindole (DAPI) counterstain, mouse anti-APP (22C11) and rabbit anti-BACE1 with Amylo-Glo counterstain, and BACE1 immunolabeling with DANIR-8c and DAPI counterstain.

The immunolabeling was visualized using Alexa Fluor® 488 and Alexa Fluor®594 conjugated donkey anti-mouse and anti-rabbit secondary antibodies (1:100, Invitrogen, Carlsbad, CA, United States). Following the fluorescent staining, the sections were briefly incubated in 0.1% Sudan black to block autofluorescence, then coverslipped with the VECTASHIELD Vibrance® antifade mounting medium.

### Image capture, pathological evaluation and figure preparation

Bright-field images were captured in a digital slide scanner (KF-PRO-120, Konfoong bioinformation tech co., Ltd., China) using the 20X objective, examined and extracted with the *K*-Viewer 1.0.5 software (KFBIO Digital Slide Viewer). Fluorescently labeled sections were scanned with a Keyence KV-8000 imaging system (Keyence Corporation, Osaka, Japan) using a Z-stack setting of 1 µm. The images were examined with the BZ-X800 Wide Image Viewer (Keyence Corporation, Osaka, Japan), with the whole section image or selected regions exported. The resulting image files were further edited with the BZ-X800 Analyzer (Keyence Corporation, Osaka, Japan), including the preparation of overlapped and channel-separate micrographs, and insertion of scale bar or captured local areas. Figures were assembled with Photoshop 2024, with brightness and contract of the whole figure adjusted as needed.

## Results

### Brief case information and macroscopical brain examination

Clinical diagnosis and demographic information of the donors and major histopathological findings are summarized in Table 1 accordingly. The hematological malignance group consisted of cases died of acute myeloid leukemia (AML, n=9, 10-50 y), acute lymphocytic leukemia (ALL, n=2, 24 y and 31 y), myelodysplastic syndrome (MDS, n=1, 48 y), lymphoma (n=2, 51 y and 87 y) and multiple myeloma (MM, n=3, 62-74 y). The 74 y MM patient (#17) had memory loss including confusion of time, space and family members, mood swings and communication difficulties during the past two year before death. Other cancer patients were cognitively intact before the terminal illness. In the cardiovascular disease group, the 65 y patient (case #19) with chronic cerebral ischemia were bedridden and had difficulty in communication for about two years before death, while the other two cases were cognitively intact before the last hospitalization. On examination of the whole brain or the fresh-prepared and formalin-fixed slices, no macroscopic infarction, hemorrhage, spongiform lesion and particular atrophy of the cerebral lobes or the hippocampus were observed among the samples except for case #20, who had a clinical record of hypertension and coronary artery disease and died one day with an acute cerebral stroke diagnosed as subarachnoid hemorrhage (SAH). A large amount of blood was seen in the posterior cranial fossa and cerebral ventricles at brain collection, while the cerebral parenchymal structures were largely intact. Images of the fixed brain slices from representative cases are provided as supplemental material (Supplemental Figures 1-6). Braak stage and Thal phase indicated in the table were scored based on the regional involvement of the lesions by examining paraffin sections of all neuroanatomical regions stained with AT8, Gallyas silver and 6E10 (Braak and Braak, 1991; Braak et al., 2006; Thal et al., 2002) (Supplemental Figure 7).

### Microscopical labeling pattern in the brains without neuropathological findings

Microscopical images of brain sections from the 14 y donor with AML (Case #4 in Table 1) are shown as examples to illustrate the histological and antibody labeling patterns in the absence of neuropathological changes (Figure 1). Signs of mild vascular injury were incidentally observed in sections with HE stain, which appeared as a widening of the perivascular space at some intracerebral blood vessels (Figure 1A, A1). The relative amounts of red blood cells and white blood cells (nucleated, generally <5%) inside the blood vessels appeared to be in a normal proportion (Figure 1A1). In Prussian blue stain with eosin counterstain, a small number of aged red blood cells (stained brown) and nucleated blood cells were seen outside the walls of some vessels by closer examination (Figure 1B, B1, insert). A few lightly stained iron deposits could be identified, primarily in the white matter (WM) (Figure 1B2, insert). The 6E10 antibody did not visualize any specific labeling in the sections of this brain, as seen in the temporal lobe section covering the neocortex, entorhinal cortex and hippocampal formation (Figure 1C, C1, C2). There were also no neuronal or neuritic profiles labeled by the AT8 antibody across the regions (Figure 1D, D1, D2). The APP (Y188) antibody displayed the normally arranged axonal fiber systems in the WM as well as the neural tracts in the hippocampal formation including the perforant path (PP) and alveus (Alv) (Figure 1E, E1, E2). The BACE1 antibody exhibited a neuropil labeling pattern across the cortical grey matter and hippocampal cellular layers, with the mossy fiber (mf) terminals in the dentate gyrus and CA3 distinctly labeled (Figure 1F, F1, F2), consistent with previously reported BACE1 labeling pattern in mammalian brains (Zhang et al. 2009, Cai et al. 2010).

**Figure 1.**
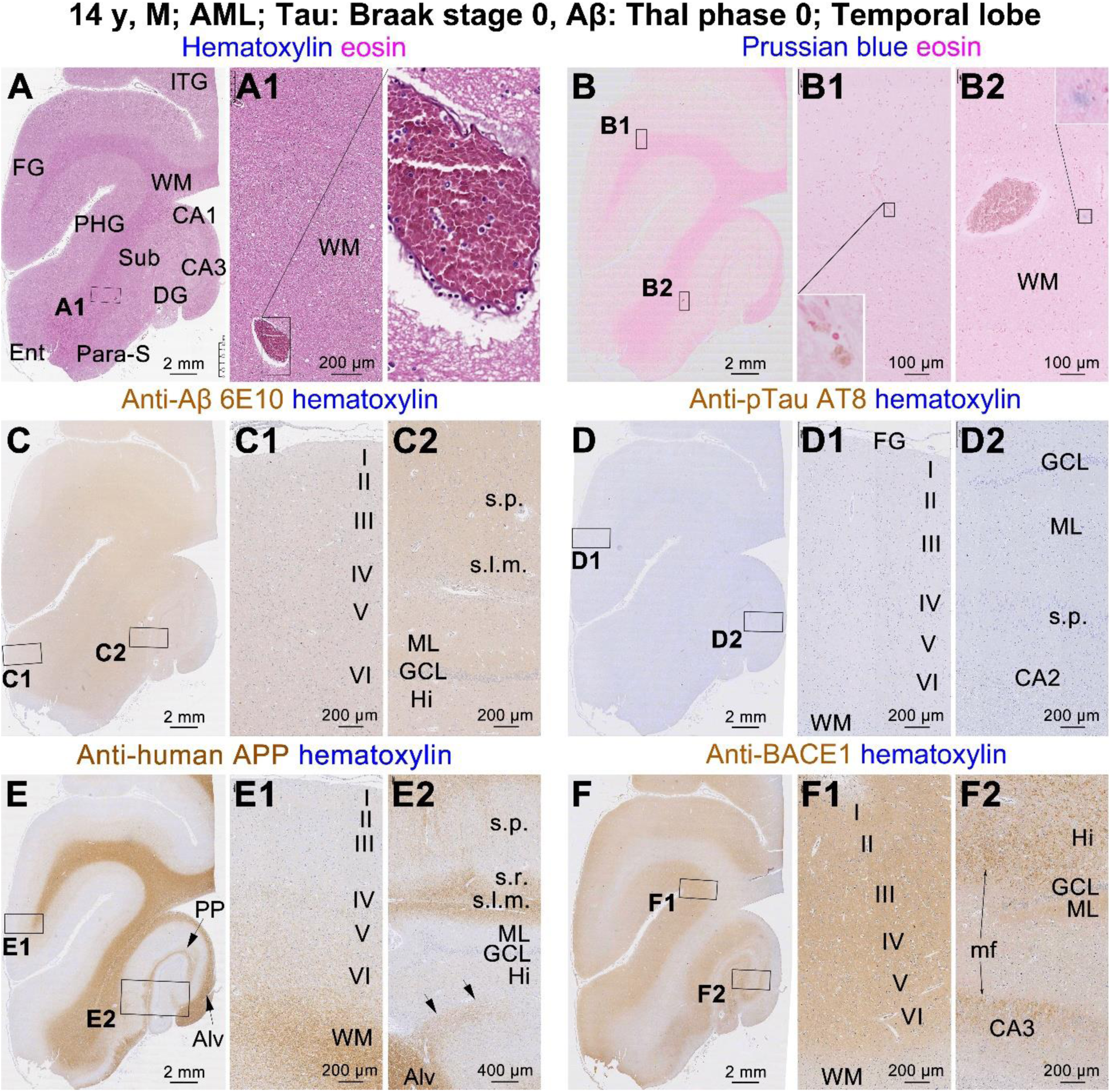
Histological and immunohistochemical labeling patterns in the temporal lobe sections from a youth case (#4 in Table 1) of AML without microscopically detectable neuropathologies. (**A, A1, A2**) Hematoxylin and eosin (HE) stain shows increased perivascular space and a normal-looking proportion of red and white blood cells in a large white matter (WM) blood vessel. (**B, B1, B2**) Prussian blue stain shows perivascular microbleed (brown-stained and nucleated cells) and mild iron deposition in the WM. (**C, C1, C2**) The 6E10 antibody displays light background reactivity across the temporal lobe structures. (**D, D1, D2**) The AT8 antibody exhibits a negative immunolabeling in the section. (**E, E1, E2**) The immunoreactivity of Y188 APP antibody is localized to the WM and axonal tracts of the hippocampal formation including the perforant path (PP) and alveus (Alv), with a part of axonal fibers (arrows) of the latter entering the hilus (Hi) of the dentate gyrus (DG). (**F, F1, F2**) The BACE1 antibody marks a diffuse neuropil reactivity in the cortical grey matter and hippocampal cellular layers, with little labeling in the WM and axonal bundles. However, the mossy fiber (mf) in the hilus and CA3 are distinctly labeled. Additional abbreviations: ITG: inferior temporal gyrus; FG: fusiform gyrus; PHG: parahippocampal gyrus; Ent: entorhinal cortex; Sub: subiculum; Para-S: parasubiculum; CA1, CA2 and CA3: Ammons horns subregions; I-VI: cortical layers; s.p.: stratum pyramidale; s.r.: stratum reticulata; s.l.m.: stratum lacunosum-moleculare; ML: molecular layer; GCL: granule cell layer.

### Tau-independent extracellular Aβ deposition in the four adult brains

In the brain from the 31 y donor with AML (case #9 in Table 1), the 6E10 antibody labeled extracellular Aβ plaques in the cerebrum with an overall low burden, including in the frontal, occipital and temporal neocortex (Figure 2A, A1, A2, B, B1, B2; Supplemental Figure 8A, A1-A5), but not in the hippocampal formation, insula, striatum and other subcortical regions (Supplemental 8A6; Supplemental Figure 9A A1-A3). The plaques appeared to be diffuse-like, with some large-sized ones strongly labeled (Figure 1A2, B2). The 6E10 labeling also occurred locally in the WM, with the labeled profiles consisting of swollen axonal processes mostly distributed around blood vessels (Figure 2A, A1, B, B1). Swollen axonal processes and sphericles were present in the striatum and internal capsule, packed into several large neuritic clusters in the globus pallidum (GP) and internal capsule, with others sparsely distributed in these regions (Supplemental Figure 9A, A1-A7). Extracellular Aβ deposition was present around the densely packed clusters. No pTau immunolabeled neuronal and neuritic profiles were found in this brain, including in the hippocampal and entorhinal subregions (Supplemental Figure 8B, B1-B4; Supplemental Figure 9B, B1-B8).

**Figure 2.**
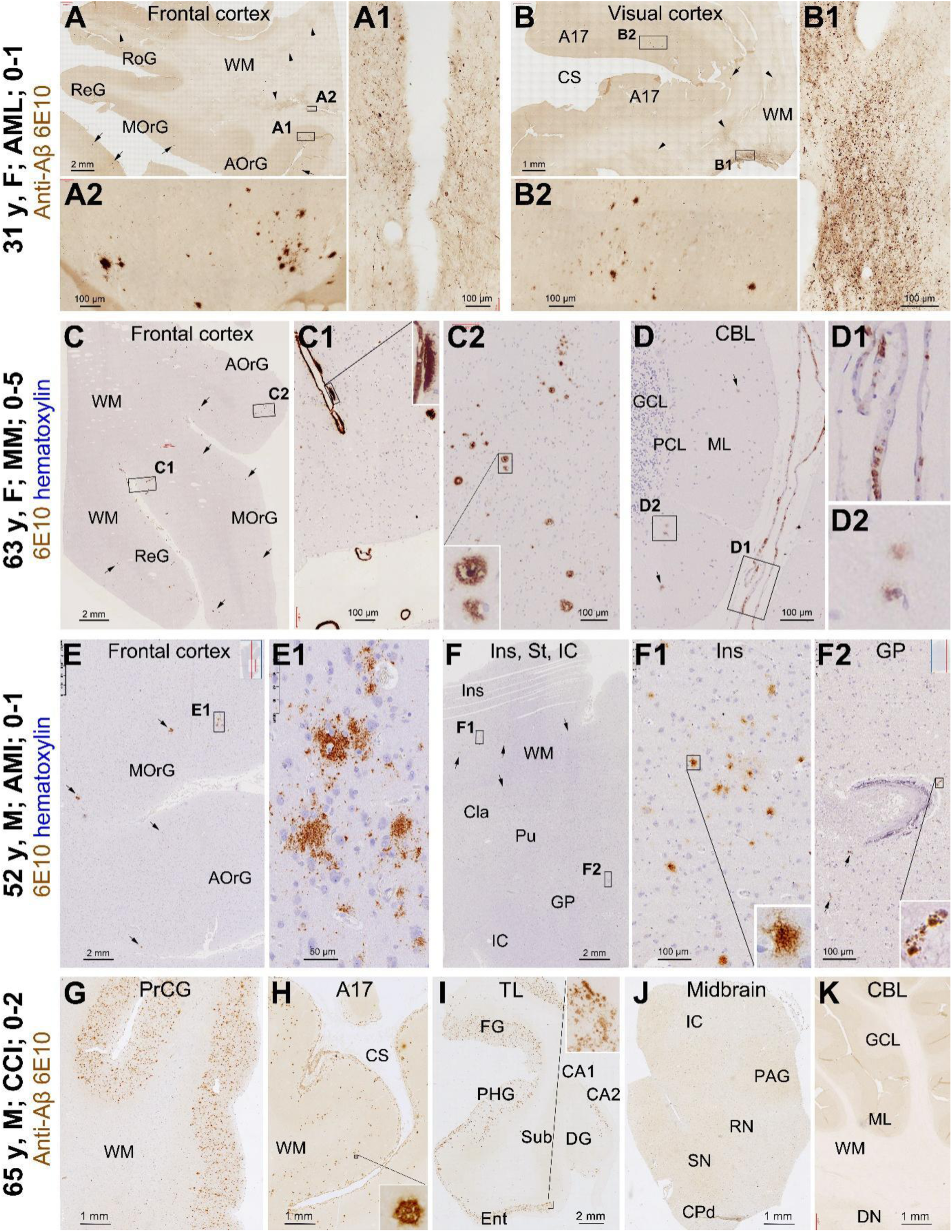
Immunolabeling revealed by the 6E10 antibody in representative sections from four adult brains. The case information indicated on the left includes age in years (y), sex (M/F), clinical diagnosis of death-causing disease and Braak (0) and Thal (1, 2, 5) scores of tau and Aβ pathologies assessed in the brain. (**A, A1, A2, B, B1, B2**) Frontal and visual cortex of the 31 y case with acute myeloid leukemia (AML), 6E10 immunolabeled Aβ plaques (arrows) are present in the grey matter (**A2, B2**), with swollen axonal processes in local white matter areas (arrowheads), especially dense around blood vessels (**A1, B2**). (**C, C1, C2, D, D1**) Frontal and cerebellar cortices of the 63 y case with multiple myeloma (MM). Compact and diffuse plaques (arrows) as well as cerebral amyloid angiopathy (CAA) are present in the neocortical grey matter, some compact plaques are cored (**C2**). CAA is found in leptomeningeal and intracerebral vessels; some associated with perivascular plaques (**C1**). In the cerebellum, small-sized plaques are present in the molecular layer (**D, D2**), with lightly stained cellular profiles in the epithelial layer of pial vessels (**D1**). (**E, E1, F, F1, F2**): Frontal cortical and insular level sections from the 52 y case died of acute myocardial infraction (AMI). Diffuse Aβ plaques are present in the frontal and insular cortices (Ins) but not seen in the subcortical structures as shown in the insular level section. Vascular injury is manifested as increased perivascular space in the cortex. (**F1**) shows a blood vessel in the globulus pallidum (GP) with calcification in the wall and perivascular axon terminal-like 6E10 labeling. (**G, H, I, J, K**): Sections from multiple brain regions as indicated from the 65 y case of chronic cerebral ischemia (CCI). Note the heavy load of Aβ plaques in the precentral gyrus (PrCG) (**G**), temporal neocortical and entorhinal cortical areas (**I**), relatively low amounts of plaques in the primary visual cortex or area 17 (A17) (**H**), and a lack of plaques in the subicular and CA1 subregions (**I)**. The cortical plaques are predominantly the diffuse type, while a small number of cored compact plaques are also found (**H**, insert). No plaques are present in subcortical structures including the midbrain (**J**) and cerebellum (CBL) (**K**). Additional abbreviations: AOrG: anterior orbit gyrus; MOrG: meddle orbit gyrus; ReG: rectal gyrus; RoG: Rostral cingulate gyrus; CS: calcarine sulcus; GCL: granule cell layer; PCL: Purkinje cell layer; ML: molecular layer. WM: white matter; St: striatum; Cla: claustrum; Pu: putamen; IC: internal capsule; IC: inferior colliculus; PAG: periaqueduct grey; RN: red nucleus; SN: substantia nigra; Cpd: cerebral peduncle; LC: RF: reticular formation; DR: dorsal raphe; CBT/CST: corticobulbar and corticospinal tracts; DN: dentate nucleus. Scale bars are as indicated.

In the brain of the 63 y donor with MM (case #16), the 6E10 antibody labeled multiple forms of Aβ pathology throughout the cerebral cortex and in some subcortical structures with a low burden. Thus, compact and diffuse plaques as well as cerebral amyloid angiopathy (CAA) were found in all cerebral lobes, the claustrum, striatum, thalamus and cerebellum (Figure 2C, D; Supplemental Figure 10). As the lesions occurred in the cerebellum, we scored the Aβ pathology as Thal phase 5, although such a wide distribution was atypical since this score would typically be seen in AD patients (Thal et al., 2002). No pTau or tangle profiles were found in the brain, including in the hippocampus and locus coeruleus (Supplemental Figure 11).

In the brain of the 52 y donor (case #18) died following an acute myocardial infarction, the 6E10 antibody labeled small amounts of diffuse plaques in the frontal cortex and insular cortex (Figure 2E, E1, F, F1) and occasionally in other neocortical areas (not shown). No plaques were found in the hippocampal formation (Supplemental Figure 12A), striatum and internal capsule (Figure 2F, F2). The AT8 antibody did not label neuronal or neuritic profiles in any brain region, as shown in the temporal lobe section (Supplemental Figure 12B, B1) and in the section covering the insula, striatum and internal capsule (Supplemental Figure 12D, D1-D4).

In the brain of the 65 y donor (case #18) with chronic cerebral ischemia (CCI), Aβ plaques were densely packed in most neocortical areas, with a relatively lower burden in the visual cortex. A few plaques were found in the CA2 area and the molecular layer of the dentate gyrus, but not in CA1 and subicular areas in the hippocampal formation, nor in the subcortical structures (Figure 2G-K; Supplemental Figures 13, 14). The Aβ plaques appeared to be predominantly the diffuse type, although some compact plaques including those with a core could be found by close examination. CAA and subpial deposition were also present in the sections (Figure 2G, H). No neuronal or neuritic profiles were observed in the brain either with AT8 immunolabeling or Gallyas silver stain (Supplemental Figures 13, 14).

### Vascular and axonal pathologies in the adult brains with tau-independent Aβ deposition

Vascular injury indicated by iron leakage and axonal pathology labeled by the 6E10, APP and/or BACE1 antibodies were observed and sometimes anatomically associated to each other in the above four adult brains. Thus, in the 31 y AML case, the APP (Y188 antibody ) immunolabeling was heavy and uneven in the WM of the same cerebral regions with 6E10 labeled Aβ plaques, with the labeled axons had local swellings (Supplemental Figure 15A, B). 6E10 and APP labeled axonal pathology was also evident in the insular WM, striatum and internal capsule (Supplemental Figure 15C, C1). BACE1 immunolabeling appeared to be increased but uneven in the cortical WM (Supplemental Figure 16A, A1, A2; B, B1), with swollen axons and clusters present in the internal capsule (Supplemental Figure 16C, C1-C3). Thus, in Prussian blue counterstained sections, iron deposition could colocalize with 6E10, APP and BACE1 labeled neuritic clusters, although it was not consistently present in the clusters with extracellular Aβ deposition (Figure 3A-G; Supplemental Figure 17A-C and enlarged panels).

**Figure 3.**
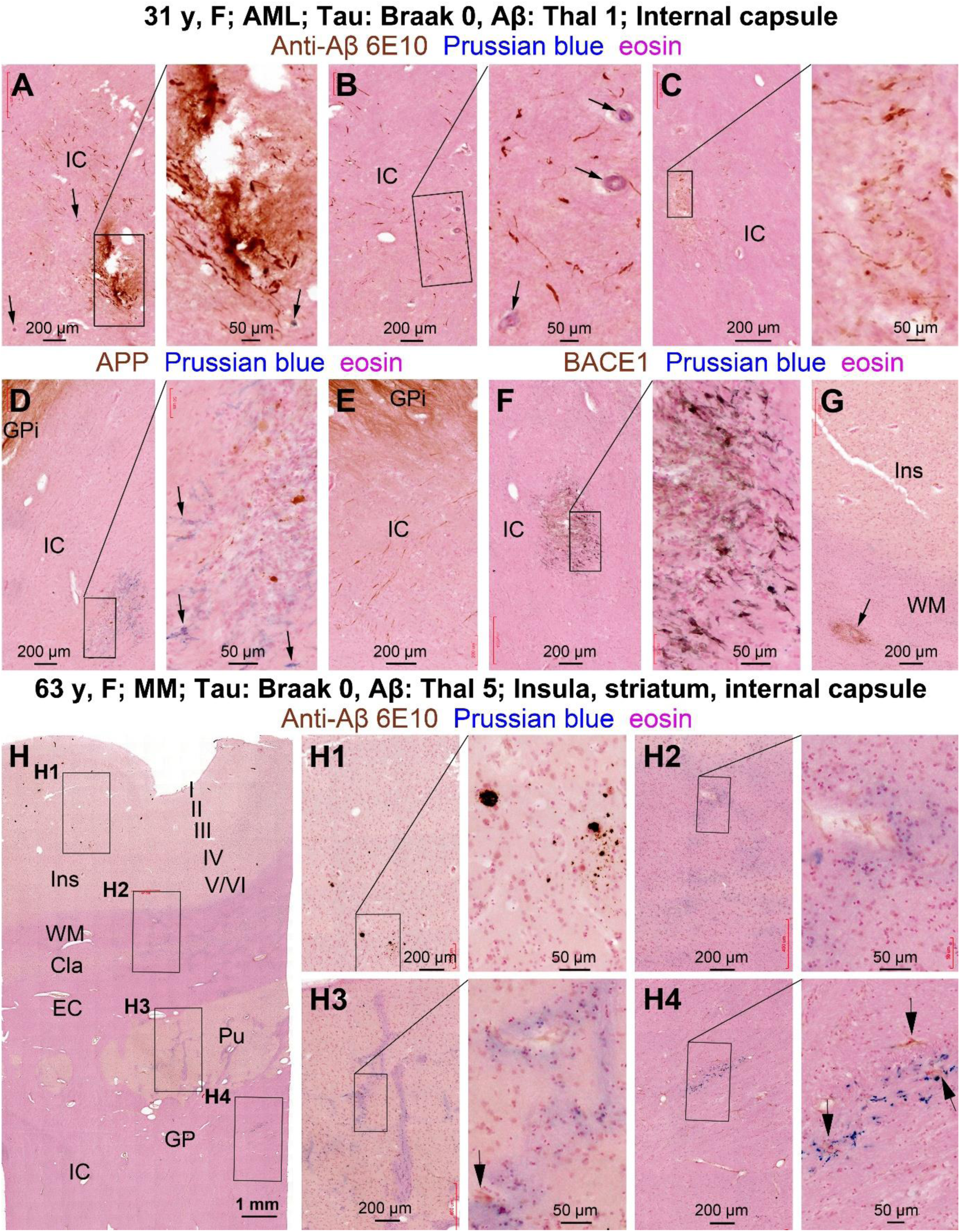
6E10 labeled Aβ deposition and axonal pathology relative to iron leakage in representative sections from two adult human brains with hematological malignancies. Case information, histological preparation and scale bars are as indicated. Panels (**A-G**) and enlarged fields are high magnification images from the internal capsule region in the brain of the 31 y AML case, referring to Supplemental Figures 15-17 for whole section views. Swollen axonal processes are labeled by the 6E10, APP and BACE1 antibodies and are densely packed in areas counterstained by Prussian blue or with extracellular Aβ deposition. Some small blood vessels show hyalinization in the wall (arrows). Panels (**H, H1-H4**) and enlarged areas show 6E10 immunolabeling with Prussian counterstain in an insular level section from the 63 y case with MM. Aβ plaques are present in the insular cortex (Ins) (**H, H1**). Iron stain is present around some blood vessels in cortical white matter and the striatum, and also in some striatal fiber bundles. A few 6E10 immunolabeled swollen axons (arrows) are seen around the blood vessels with perivascular iron deposition (**H2-H4**). Abbreviations: I-VI: cortical layers; WM: white matter; Cla: claustrum; EC: external capsule: Pu: putamen: GPi: internal globulus pallidum: IC: internal capsule.

In the 63 y MM case, rosette-like neuritic clusters were present in the neocortex in BACE1 immunolabeling (Supplemental Figure 18A, B). Double immunofluorescence confirmed a colocalization of these presynaptic sphericles with Aβ deposition, collectively representing neuritic compact plaques (Supplemental Figure C). In the cerebellum, BACE1 labeled neuritic clusters were small in size and present in the molecular and granule cell layers (Supplemental Figure 18D, D1). Light BACE1 labeling was seen in endothelial layer in blood vessels with mild CAA lesion (Supplemental Figure 18E). In 6E10 labeled sections with Prussian blue counterstain, a diffuse iron deposition was seen in the WM or neuronal fiber bundle areas, with distinct deposition around blood vessels (Figure 3H, H1-H4; Supplemental Figure 19).

In the 52 y AMI case, vascular injuries were found in brain sections with HE and Prussian stains, including perivascular and WM edema, microbleed, vascular hyalinization and calcification, which were particularly evident in the striatum and internal capsule (Supplemental Figure 12C, C1, E, E1-E4). BACE1 and APP antibodies labeled thickened axonal processes around some damaged blood vessels (Supplemental Figure 12F, F1, F2, G, G1). In the frontal and insular cortex, Aβ plaques were present around blood vessels with increased perivascular space, while neuritic-like 6E10 labeling could be seen around the injured vessels in the internal capsule associated with microbleed and/or wall calcification (Figure 2E1, F1, F2).

In the 65 y case of chronic cerebral ischemia, CAA and vascular injury were present in the neocortex besides Aβ plaques (Figure 2G, H), as well as in the stratum lacunosum-moleculare (s.l.m.) in CA2 and around the pia of the DG (Supplemental Figure 13A, A1, A2). Increase of perivascular space, perivascular edema, thinning and breakdown of vascular wall were observed in HE stain in all neocortical regions, hippocampal formation, striatum and internal capsule (Supplemental Figures 13D, 14C, C1, F, F1). The axonal APP labeling appeared to be enhanced or denser around blood vessels especially those in the cortical WM and striatum with perivascular iron leakage, although there were no apparently swollen sphericles or structural disruption along the axonal processes (Supplemental Figures 13E, E1, E2; 14J, J1, J2). BACE1 immunolabeling appeared patch-like with locally enhanced reactivity in the cortex with a large amount of the diffuse Aβ plaques (Supplemental Figures 13F, F1, F2).

### Vascular and axonal pathologies in other youth/adult cases of hematological malignances

Vascular injuries and axonal pathologies were found in the brains of youth and adult cases of hematological malignances in which no extracellular Aβ plaques were evident, with one brain (case# 16) had tauopathy at Braak stage II. Overall, vascular injuries appeared as increased perivascular space, vessel wall breakdown, microbleed and/or iron deposition. Axonal pathology was found in a subset of brains, appearing as swollen axonal processes or sphericles in 6E10, APP or BACE1 labeling. The major pathological findings are described below (see Table 1).

In the brain of case #1 (10y, AML), microbleed and minor iron leakage were observed in HE and Prussian stained sections, as shown in the striatum level sections (Supplemental Figure 20). The immunolabeling did not reveal neuropathological changes (not shown). In the brain of case #2 (17 y, AML), microbleed without apparent iron leakage and immunolabeled neuropathologies were observed in the brain sections as shown in the temporal lobe sections (Supplemental Figure 21). In case #3 (37 y, AML), microbleed was found in the temporal lobe and striatum level sections, with localized axonal pathology also seen in the latter region (Supplemental Figure 22). In case #5 (16 y, AML), apparent intracerebral vascular disruption, microbleed and iron leakage were present in the striatum and internal capsule, while no apparent alteration in APP and BACE1 immunolabeling were observed (Supplemental Figure 23). In case #6 (50 y, AML), intracerebral and meningeal microbleed were evident in the temporal lobe and striatum, with axonal pathology in the latter by 6E10, APP and BACE1 immunolabeling (Supplemental Figure 24). In case #7 (21 y, AML), pathological findings included vascular injury associated with cancer cell infiltration (will be addressed further), minor iron leakage and localized axonal pathology around injured vasculature (Supplemental Figures 25, 26). In case #8 (16 y, AML), vascular injury, cancer cell infiltration and diffuse axonal pathology were found the brain sections (Supplemental Figures 27).

Case #10 was a 32 y with acute lymphocytic leukemia (ALL), vascular damage, microbleed and iron leakage were seen in the cerebral and striatum level sections without overt axonal pathology (Supplemental Figure 28). In the brain of case #11 (24 y, ALL), iron leakage was evident around vasculature and in axonal bundles in the striatum and internal capsule (Supplemental Figure 29). Case #12 was a 48 y with myelodysplastic syndrome, tumorous cell infiltration was evident at meningeal and intracerebral vasculature, which were associated with microbleed, iron leakage and axonal pathology (Supplemental Figures 30, 31). Case #13 was a 52 y with mantle cell lymphoma. An increased proportion of nucleated blood cells was seen in the meningeal blood vessels, with vascular disruption, microbleeds and minor iron leakage around these and some intracerebral blood vessels (Supplemental Figure 32).

Case #15 was a 62 y patient died of MM. The AT8 antibody visualized pTau positive neurons and neurites restricted to the entorhinal cortex, with a few labeled neuronal somata and processes in hippocampal formation by closer examination, thus exhibiting a distribution pattern consistent with NFT as Braak stage II. There were no extracellular Aβ deposition found in this brain. However, the 6E10, APP and BACE1 antibodies consistently labeled axonal pathology in local WM regions in the cerebrum, the striatum and internal capsule, all of which were associated with perivascular microbleeds, hyalinosis/calcification in the vascular wall, or localized iron leakage (Supplemental Figures 33, 34).

### Vascular and axonal pathologies in the elderly brains with **Aβ** deposition and tauopathy

Two elderly brains from blood cancer patients and one with cardiovascular disease had Aβ deposition and tauopathy along with vascular injury and axonal pathology. Brain #14 was from an 87 y patient with lymphoma, a few extracellular Aβ plaques (Thal phase 1) was found in the frontal and parietal cortical regions, but not in other cortical regions, the hippocampal formation and subcortical structures (Figure 4A; Supplemental Figures 35, 36A). The AT8 antibody labeled neuronal and neuritic profiles in the hippocampal formation and basal neocortical areas, with a few in the insular cortex, consistent with a Braak stage IV score (Figure 4B; Supplemental Figure 36B). Despite the very mild extracellular Aβ deposition in the neocortex , 6E10, APP and BACE1 labeled swollen axons were present in the cortical WM and internal capsule, especially in the areas with vascular lesions and iron leakage (Figure 4A, C, D; Supplemental Figures 35B, B1-B3; 37). In fact, these antibodies labeled axonal elements inside the wall of the damaged striatal arteries (Figure 4A1, C1, D1). Brain # 17 was from a 74 y donor clinically diagnosed as MM with Alzheimer-type dementia. In HE and Prussian blue stains, perivascular edema and microbleed were prominent in the insular level sections, with iron leakage around blood vessels and in the axonal fiber bundles (Figure 5A, A1, A2, B, B1, B2). Extensive diffuse and compact Aβ plaques as well as CAA existed in the cerebral and subcortical regions, consistent with the pattern of Thal phase 4 (Figure 5C, C1, C2; Supplemental Figure 38A, A1-A4). AT8 labeled neuronal tauopathy was scored as Braak stage IV in this brain, as illustrated in the temporal lobe and insular level sections (Figure 5D, D1, D2; Supplemental Figure 38B, B1-B4). The BACE1 and APP antibodies visualized dystrophic neuritic clusters in the grey matter consistent with the presence of compact or neuritic plaques. These antibodies also labeled swollen axonal processes in the cortical WM, striatum and internal capsule (Figure 5E, E1, E; F, F1, F2; Supplemental Figure 38C, D and enlarged panels).

**Figure 4.**
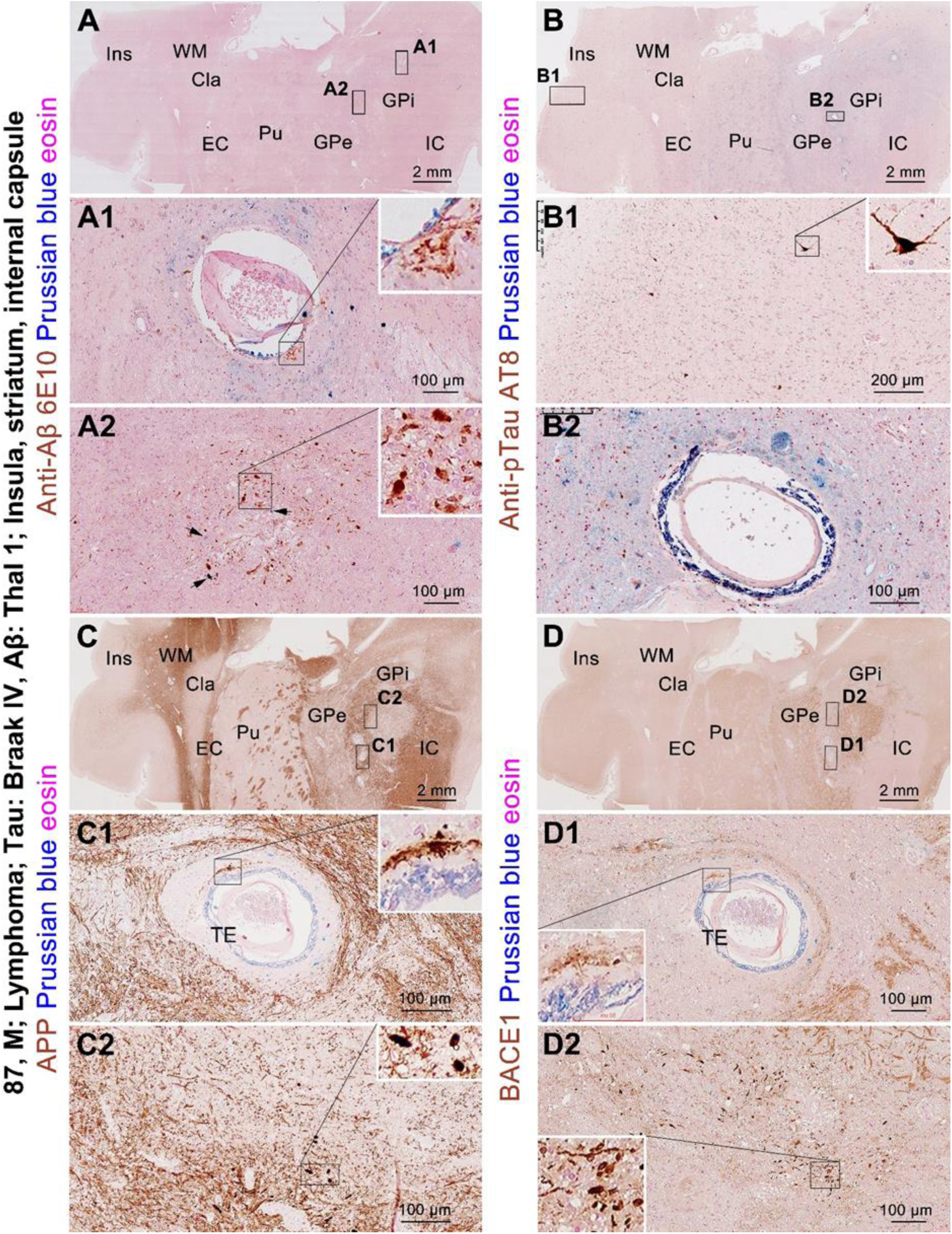
Vascular injury and amyloidogenic axonal pathology in the insular level sections from the 87 y case of lymphoma. (**A, A1, A2**) No Aβ plaques are labeled by the 6E10 antibody across the anatomical areas including in the insular cortex (**A**). However, swollen axonal processes are labeled by this antibody around injured blood vessels and inside the vascular wall with local iron deposition in the striatum and internal capsule (**A1, A2**, arrows). (**B, B1, B2**) A few pTau immunoreactive neurons are seen in the insular cortex (**B1**), but no pTau labeled axonal structures are present around injured blood vessels (**B2**). (**C, C1, C2**) The APP labeled axonal processes are heavily packed around injured blood vessels in striatum, with some forming sphericles (insert in **C2**). Iron deposition and sprouting axons also occur inside the wall of the blood vessels (**C1**, insert). (**D, D1, D2**) In a consecutive section, BACE1 labeled axonal pathology exhibit the same pattern as with the 6E10 and APP labeled counterparts, including inside the wall of large striatal arteries with iron infiltration (**A1, C1, D1**). Ins: insular cortex; WM: white matter; Cla: claustrum; EC: external capsule; Pu: putamen; GPe: globulus pallidum external part; GPi: globulus pallidum internal part. IC: internal capsule. Scale bars are as indicated.

**Figure 5.**
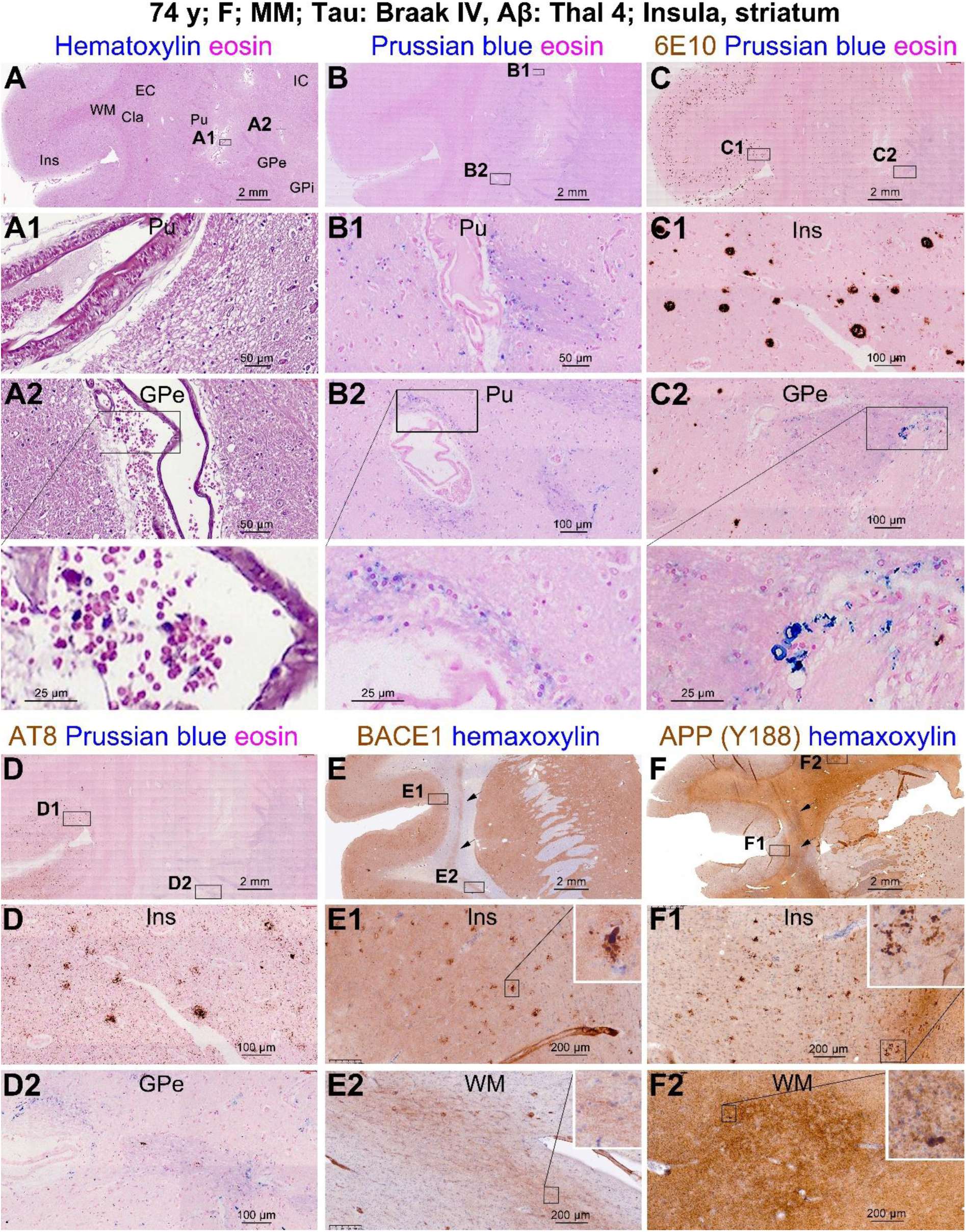
Comparative assessment of vascular injury, Aβ deposition, tauopathy and axonal pathology in the striatum level sections from the 74 y donor with multiple myeloma. In HE (**A, A1, A2**) and Prussian (**B, B1, B2**) preparations, microbleed (arrows) and iron deposition are evident around blood vessels in the striatum and internal capsule, with iron infiltration also seen in the axonal fiber bundles. (**C, C1, C2**) Aβ plaques are labeled in the insular cortex, claustrum and putamen. The blood vessels show increased perivascular space, some associated with iron deposition and localized axonal labeling (**C2,** enlarged area). (**D, D1, D2**)Tau positive neuronal, neuritic clusters and neuropil threads are present in the insular cortex, with labeled neurites in the striatum. BACE1 (**E, E1, E2**) and APP (**F, F1, F2**) labeled dystrophic presynaptic terminals are present in the insular cortex and arranged as clusters (**E, E1, F, F1**). Fine axonal processes are also labeled by these antibodies in the insular white matter, with the staining locally enhanced (**E2, F2**). The anatomical areas are indicated in the HE stained section. Ins: insular cortex; WM: white matter; Cla: claustrum; EC: external capsule; Pu: putamen; GPe: globulus pallidum external part; GPi: globulus pallidum internal part. IC: internal capsule. Scale bars are as indicated.

Brain #20 was from an 82 y patient with hypertension and coronary artery disease died following acute SAH. In the temporal lobe, the 6E10 antibody labeled extensive diffuse plaques throughout the neocortex and entorhinal cortex, with only a small amount of Aβ deposition in the dentate gyrus near the pia and none in other hippocampal areas (Figure 6A, A1-3). Further, few plaques were found in subcortical structures including the striatum (Supplemental Figure 39A, A1-A5). However, 6E10 labeled swollen axonal processes were packed in local WM areas (Figure 6A, A1-A3). The AT8 antibody detected a few neuronal somata with sparsely distributed neurites in the transentorhinal area, consistent with a Braak’s staging of tauopathy as 1a (Figure 6B, B1-3) (Braak and Del Tredici, 2011). The APP Y188 antibody visualized the WM and hippocampal fiber pathways, with a clearly enhanced labeling in local WM areas indicating axonal pathology, with axonal swelling and disruption seen at high magnification (Figure 6C, C1, C2). BACE1 immunolabeling was also enhanced in the same WM areas with 6E10/APP labeled axonal pathology, with swollen axonal profiles visible by closer examination (Figure 6D, D1). The hippocampal mossy fibers were labeled by the BACE1 antibody and appeared to be normal-looking (Figure 6D, D2).

### Infiltration of malignant cellular and soluble markers in the brains with blood cancers

An increased proportion of nucleated relative to non-nucleated cells inside blood vessels in HE stained brain sections suggests potential pathological infiltration of cancerous white blood cells in the region. We carried out immunolabeling using molecular marker for blood cancers in sections from five brains with infiltration of nucleated blood cells and also from the three brains with MM. In brain sections from the 31 y donor with AML (case #9), myelogenous cell markers CD68, MPO and lysozyme immunoreactive cells accumulated inside blood vessels (Figure 7A-D). In sections from the 48 y donor (case #12) with MDS, some labeled cells occurred outside blood vessels, with the CD68 labeled cells exhibiting a glial morphology, whereas MPO and lysozyme labeled cells appeared to be bloodborne myelogenous cells (no visible processes) that leaked into the perivascular space (Figure 7E-H). CD68, MPO and lysozyme immunoreactive cells were found inside blood vessels in sections from the 21 y (case #7) (Supplemental Figures 25, 26) and the 16 y (case #8) (Supplemental Figure 27) cases with AML. In the 51 y donor (case #13) diagnosed with mantle cell non-Hodgkin lymphoma, nucleated cells were accumulated in meningeal and intracerebral blood vessels, which were immunolabeled by the CD20 (Figure 7I), a cytokine marker of this type of tumor cells (Shadman et al. 2019).

**Figure 6.**
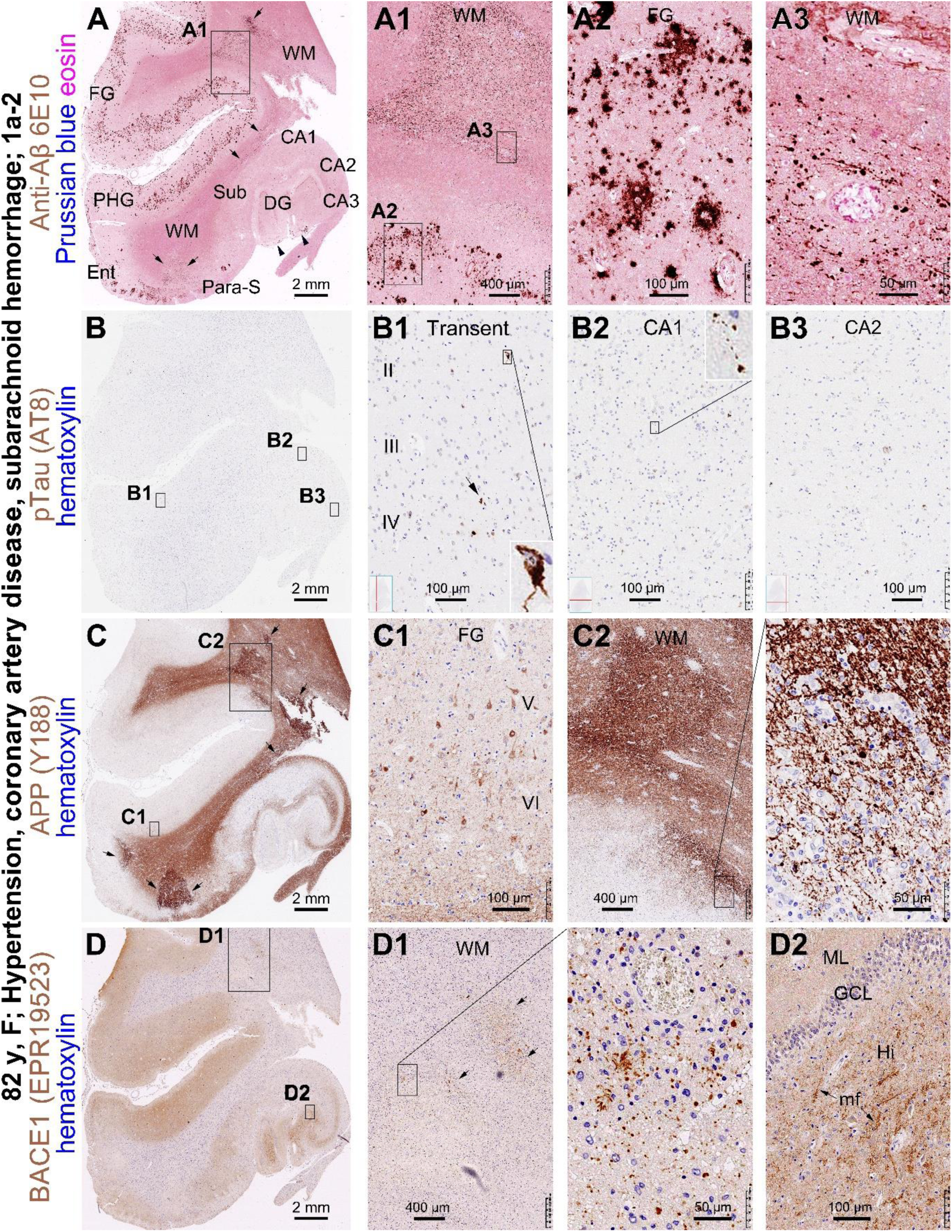
Comparative assessment of Aβ deposition, tauopathy, axonal pathology and vascular injury in temporal lobe sections of the 82 y case with cardiovascular disease. (**A**) In the 6E10 antibody labeled section with Prussian blue counterstain, diffuse Aβ plaques are loaded in the temporal neocortex and entorhinal cortex (**A1, A2**). However, only a few labeling occurs in the hippocampal formation, which is located in the dentate gyrus near the pia matter (arrowheads). 6E10 labeling is enhanced in local white matter areas (**A**, arrows), which consists of densely packed swollen axonal processes that appear to be denser around small blood vessels (**A1, A3**). (**B**) In the AT8 stained section, only a few labeled neurons are observed across the entire section; they are present only in the transentorhinal cortex (**B1**). A few labeled neurites can be found in CA1 by close examination (**B2**), while no labeled elements are seen in CA3 and DG (**B, B3**). (**C**) In the APP (Y188) labeled section, the localized white matter lesions (arrows) are highlighted. Few dystrophic neuritic clusters are seen in the cortex (**C1**). Some swollen axons are seen among the strongly labeled axonal processes in the damaged white matter areas (**C2**). (**D**) In BACE1 labeling, swollen axons are also present in damaged white matter areas and they appear denser around blood vessels (**D1**). Normal-looking mossy fiber (mf) terminals are labeled by the BACE1 antibody in the dentate gyrus (DG) (**D2**). CA1-CA3: subregions of the Ammon’s horn; Sub: subiculum; Para-S: parasubiculum; Ent: entorhinal cortex; PHG: parahippocampal gyrus; FG: fusiform gyrus; II-IV: cortical layers; ML: molecular layer; GCL: granule cell layer; Hi: hilus. Scale bars are as indicated.

**Figure 7.**
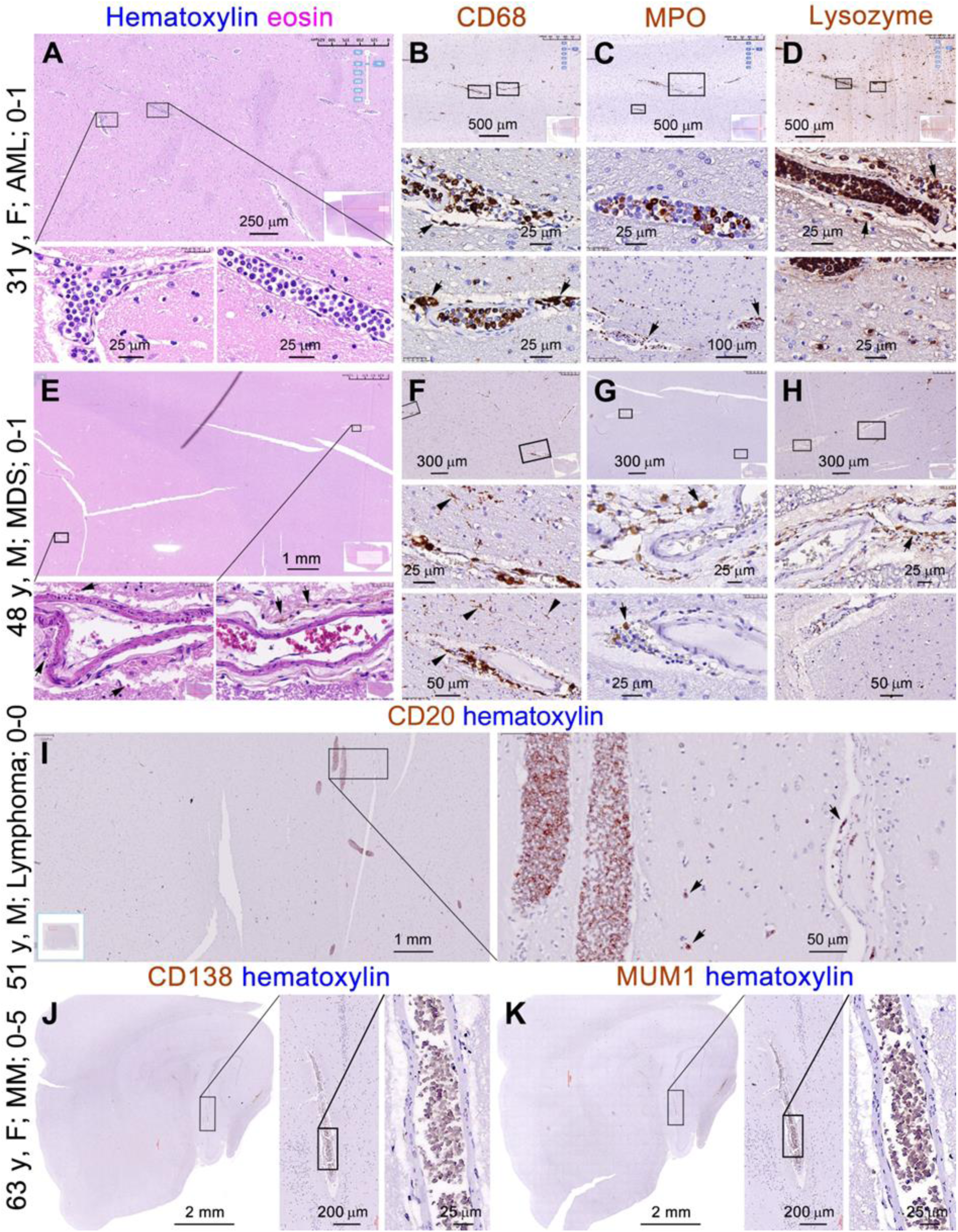
Characterization of brain infiltration of tumorous cells in four cases of hematological malignancies. The images show hematoxylin and eosin stain and immunolabeling of tumorigenic cellular markers in representative sections from four brains in the blood cancer group. The cases information include Braak (0) and Thal scores (1 and 5) of the brains. In the striatum level sections from the 31 y donor with acute myeloid leukemia (AML), nucleated blood cells are accumulated inside the blood vessels (**A**), which are partially labeled by CD68 **(B**) and MPO (**C**) antibodies and appear to be mostly labeled by the lysozyme antibody (**D**). In the striatum level sections from the case of myelodysplastic syndrome (MDS), microbleed is seen around blood vessels in the white matter and striatum (**E**, arrows). CD68 (**F**), MPO (**G**) and lysozyme (**H**) labeled cells are present in the perivascular space, which lack cellular protruding. However, additional CD68 labeled cells in small somal size and with short processes are present around and away from the blood vessels, which are morphological resembling microglia or oligodendrocytes (**F**). In the striatum level section from the 51 y case of lymphoma, CD20 immunoreactive cells accumulated in some blood vessels (I). In the temporal lobe sections from the 63 y MM case, there are no positive immunolabeling of the two plasma cell markers CD138 (**I**) and multiple myeloma oncogene 1 (MUM1) (**J**) inside or outside the brain vasculature. Scale bars are included in the image panels.

In the brains of MM cases, no blood cells were labeled by the plasma cell markers CD138 or MUM1 (Parks et al. 2019) (Figure 7J, K), but there existed aberrant immunolabeling of the immunoglobulin light chains (Figure 8). Thus, sections from the 62 y and the 74 y cases were infused by the light chain κ in the absence of λ (Figure 8A, B, C, D and enlarged panels). An opposite labeling pattern for the two light chains was found in sections from the 63 y patient (Figure 8G, H and enlarged panels; Supplemental Figure 39A-D). The infiltration was lighter in the brain of the 62 y relative to the 63 y and 74 y. Thus, the immunolabeling was located around the pia and in WM in the youngest case (Figure 8B, B1), but in the white and grey matters in the other two cases (Figure 8D, G). The light chain infiltration occurred throughout the brain including in the cerebellum and brainstem (images not shown). Some neuronal somata were also labeled, which were stained lighter in intensity and fewer in number in the 62 y brain. These labeled neuronal somata appeared shrinking and surrounded by an unlabeled perisomal cleft, with the severely shrunken somata crushed into a small remain a large unlabeled cellular zone (Figure 8D1, G2, arrows). The nuclei of the light chain labeled neurons appeared to be enlarged and stained fuzzing, or appeared to be dissolved (Figure 8D1, G2, arrowheads). For staining assay control, brain sections from the donors with AML and chronic cerebral ischemia were batch-processed along with the sections from the MM cases, with the former showed little light chain immunolabeling (Supplemental Figure 39E-H). Also, as an indication of specificity of the light chain antibodies, a strong but amorphous immunolabeling was present inside the blood vessels in the brain sections, representing the immunoglobin reactivity in the plasma of blood (Figure 8B2, D1, G1).

**Figure 8.**
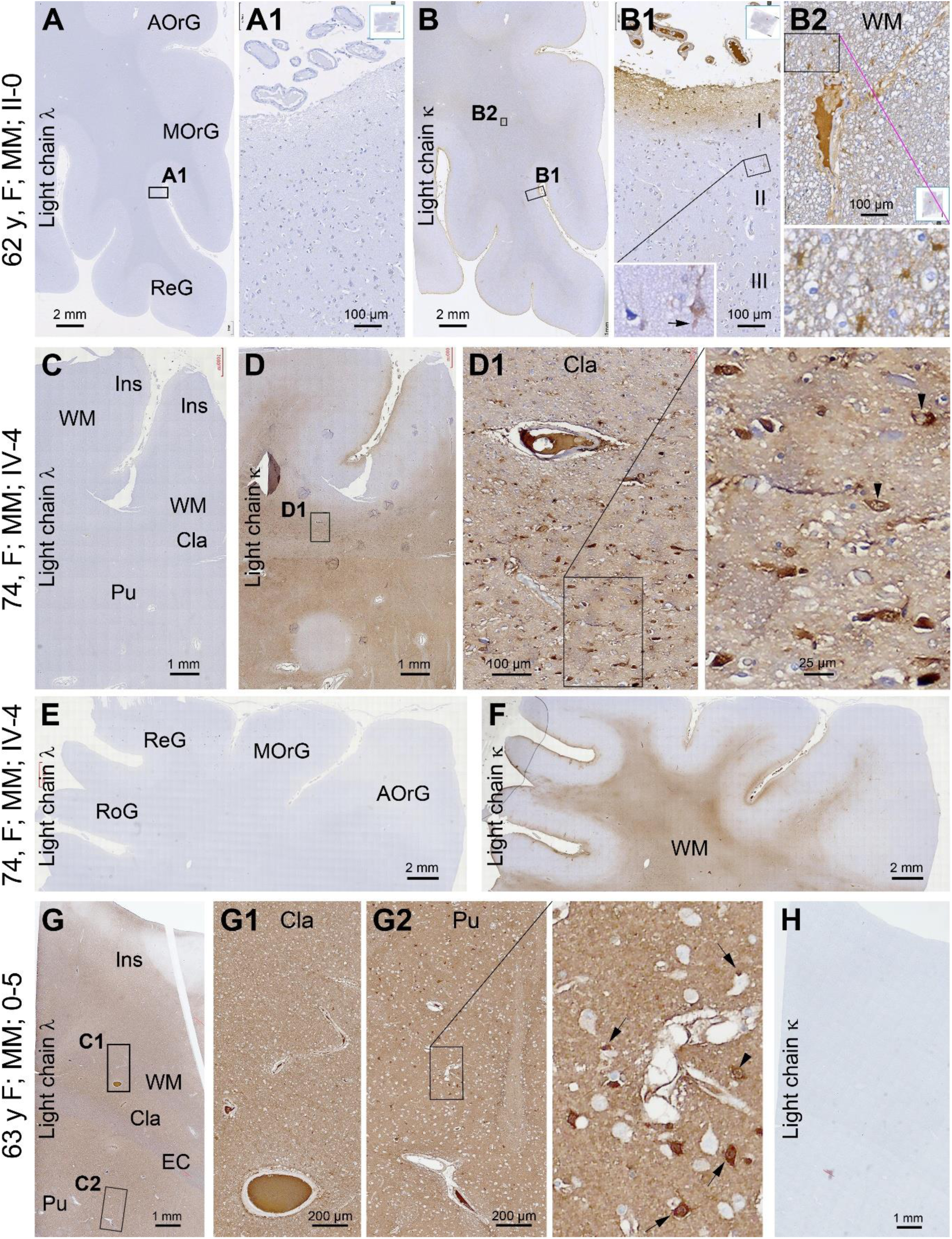
Infiltration of immunoglobin light chains into the brain in the three cases of multiple myeloma. Case information, neuroanatomical structures and light chain types are as indicated, with Braak stages and Thal phases indicated with Roman symbols and Arabic numbers, respectively. In the 62 y (**A, A1, B, B1, B2**) and 74 y (**C, D, D1, E, F**) cases, no immunoreactivity of light chain λ is present in the brain sections, whereas labeling of the light chain κ is infiltrated into the grey and white matter of the cortex, striatum and internal capsule. Note that in the 62 y case, the infusion of light chain κ is relatively light. Opposite to the above two cases, light chain λ (**G, G1, G2**) instead of κ (**H**) infiltration is present in the brain of the 63 y case. The light chain labeling also occurs in neuronal somata and proximal dendritic processes. A subpopulation of these labeled neuronal profiles appear shrunken, with some crushed into a small area in an empty cell outline (**D1 G2** and enlarged panels, pointed by arrows). The nuclei of the labeled neuronal somata appeared to be enlarged with a loss of chromatin material (Enlarged zones of **D1, G2**; pointed by arrowheads). Intravascular plasm shows the corresponding light chain immunoreactivity (**B2, D1, G1**). AOrG: anterior orbit gyrus; MOrG: medial orbit gyrus; ReG: rectal gyrus; Ins: insular cortex; Cla: claustrum; EC: external capsule; Pu: putamen; WM: what matter. Scale bars are included in the image panels.

### Characterization of the amyloidogenic antibody labeling of axonal pathology

The axonal labeling in the WM and striatum by the 6E10, APP and BACE1 antibodies in both the blood cancer and cardiovascular disease brains were impressive. To understand the nature of this pattern, we carried out further experimental characterization using the insular level sections from the brain of the 82 y case, which exhibited distinct Aβ plaques in the insular cortex and axonal pathology in the striatum that could serve an excellent internal control system for cross-validation (Supplemental Figure 40A-E and enlarged panels). Thus, a panel of additional Aβ and APP antibodies as well as fluorescent amyloid probes were applied for this purpose. As seen in consecutive paraffin sections, the APP (Y188) antibody labeled the normal-looking axonal bundles in the WM, putamen and GP with relatively light labeling intensity and fine anatomical arrangement, while it also heavily labeled the pathological axonal processes located in the putamen and packed around the border of the putamen and GP (Figure 9A, A2; Supplemental Figure 40B, B2-B5). In comparison, the 6E10, APP (22C11) and BACE1 antibodies selectively labeled the pathological axonal processes at the same above locations in the striatal subregions (Figure 9B, B1, B2, C, C1, C2; Supplemental Figure 40A, A2-A5, C, C3-C5, D2-D5), whereas these axonal profiles showed minimal AT8 labeling (Supplemental Figure 40E, E2-E5). The Aβ antibodies 6E10, 4G8, D54D2 and Ter42 all visualized the extracellular Aβ deposition predominantly as diffuse plaque profiles that were abundant in the insular cortex with a few in the striatum (Figure 9C, C1, D, D1, E, E1, F, F1). However, unlike 6E10, the 4G8, D54D2 and Ter42 antibodies only faintly visualized the above-mentioned axonal pathology in the putamen and GP (Figure 9C, C2, D, D2, E, E2, F, F2). In the triple fluorescent labeling preparation, BACE1 and APP (22C11) colocalization occurred clearly in the pathological axonal processes in the striatum, as well as in the dystrophic neuritic clusters sparsely seen in the insular cortex. The signal of the Amylo-Glo fluorescence was low across the insular cortex except for a bright labeling at the BACE1/APP colocalized neuritic clusters (Figure 9G, G1). Thus, Amylo-Glo (structurally similar to the Thioflavins) could not well display the diffuse plaques in the cortex, nor did it exhibit fluorescent labeling in the pathological axonal profiles (Figure 9G, G2). In comparison, the DANIR-8c probe labeled the diffuse as well as compact Aβ plaques in the insular cortex, with the latter associated with BACE1 immunolabeling in the presynaptic sphericles, collectively forming the compact neuritic plaques (Figure 9H, H1). However, even this potent Aβ tracer failed to exhibit binding signal inside the pathological axonal processes that were visualized by the BACE1 antibody in the striatal subregions (Figure 9H, H2).

**Figure 9.**
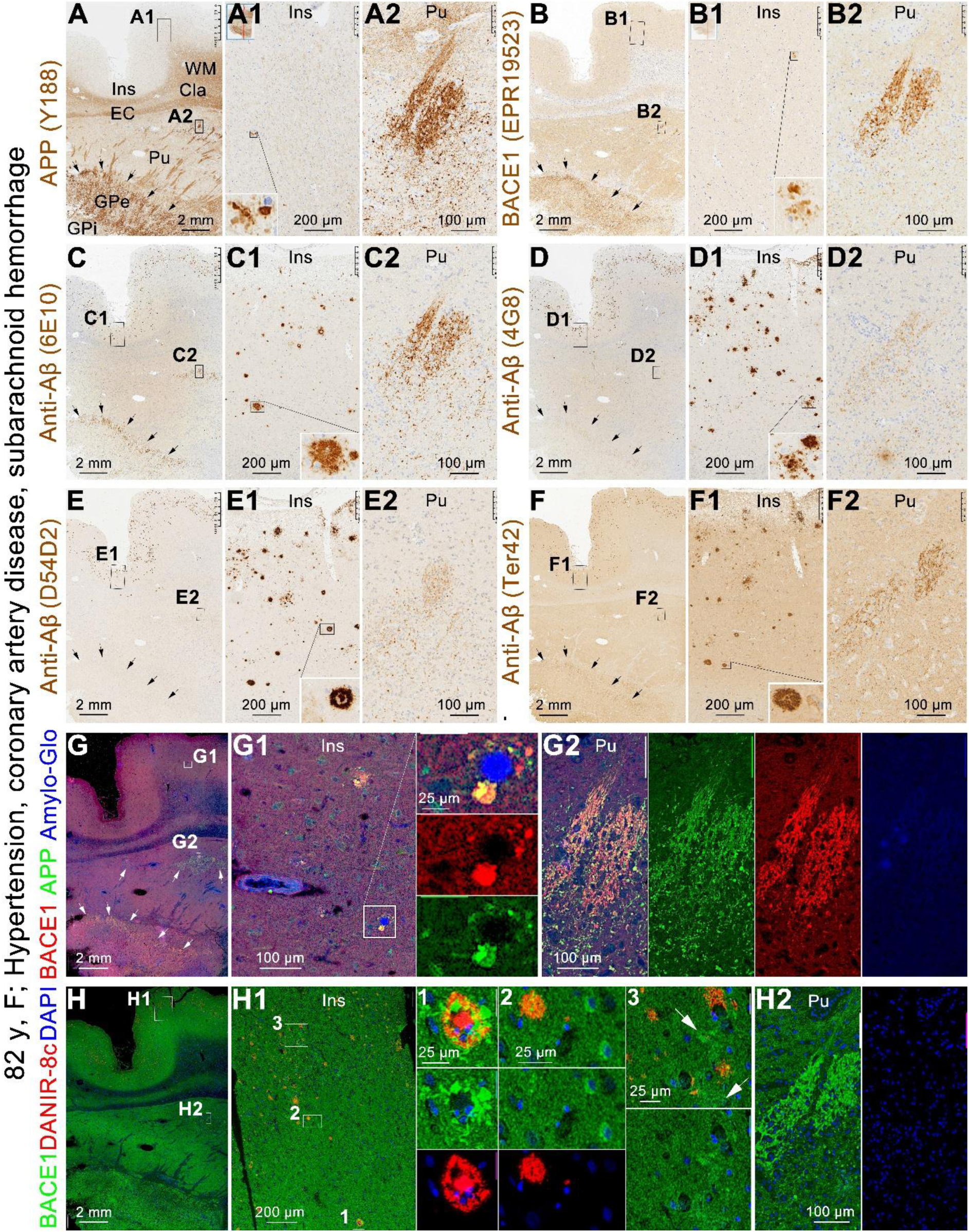
Comparative assessment of the axonal pathology with amyloidogenic protein antibodies along with Aβ antibodies and probes in consecutive insular level sections from the 82 y case with cardiovascular disease. (**A**) The APP (Y188) antibody visualizes normal axonal processes in the white matter and striatum, with an enhanced labeling of the dystrophic neuritic clusters in the cortex (**A1**), and the pathological (swollen and disorganized) axons occurring locally in the putamen (**A2**) and packed around the border between neo- and palostriata (**A**, arrows). (**B**) The BACE1 antibody also labels the aforementioned dystrophic neuritic clusters (**B1**) and pathological axonal processes (**B2**). The four Aβ antibodies all visualize the diffuse and compact (a few) plaques (inserts) in the insular cortex (Ins), and individually present in the striatum (**C, C1, D, D1, E, E1, F, F1**). However, only 6E10, but not the other three, Aβ antibodies exhibit a distinct labeling of the axonal pathology (**C2, D2, E2, F**). BACE1 and APP (22C11) labeling are colocalized in the dystrophic neuritic clusters (**G, G**1, inserts) as well as in the pathological axons (**G, G2**). Note that APP labeling by the 22C11 antibody also labels neuronal somata (**G1**). The Amylo-Glo fluorescent dye distinctly visualizes the core of the neuritic plaques but not the diffuse plaques, without labeling in the pathological axonal profiles (**G, G1, G2**). There exists Amylo-Glo labeling in the wall of some blood vessels, which may relate to autofluorescence (**G1**). (**H**) The DANIR-8c Aβ probe visualizes both diffuse and compact plaques in the insular cortex, with the latter colocalizing with BACE1 labeled neuritic clusters (**H1**, enlarged views, arrows). However, no binding signal is present in the areas with the axonal pathology (**H, H2**). Note that the BACE1 antibody also labels interneurons (**H1**, enlarged field#2) and small sized neuritic clusters away from the diffuse plaques (**H1**, enlarged field#3, pointed by arrows). Additional abbreviations: Cla: claustra; EC: external capsule; WM: white matter; Pu: putamen; GPe: external globulus pallidum; GPi: internal globulus pallidum. Scale bars are as indicated.

## Discussion

Among the 397 postmortem brains we have banked so far, the donor’s age of death ranged from infancy to 103 y (28 days to10 y, n=22; 11-25 y, n=26; 26-65 y, n=114; and 65-103 y, n= 235). In this community-based brain banking cohort, 97 samples were neuropathologically assessed as definitive PART with Braak stages from 1a to IV, with the donors died at 42 y to 96 y (mean ±SD=73±11.9 y, median=72). A total of 155 brains (54-103 y; 82±9.9 y, median=83) had AD-type Aβ plaque (Thal phase 1-5) and tau/tangle pathologies (Braak stage ≥ II). Thus, the cases of tau-independent cerebral β-amyloidosis in our brain repository are rare, consistent with existing literature (Hyman and Gomez-Isla, 1997; Braak et al. 2006; Braak et al. 2011; Tsartsalis et al., 2018; Thal et al. 2022). However, from another perspective, such cases would concur with strong factor(s) driving Aβ formation and allow a unique opportunity to explore the rationale.

Large sample size autopsy data involving brain histopathology are available from early studies. In a study of 170 cases of leukemia, the overall incidence of CNS involvement was 25%, highest in ALL and lower in AML (Wolk et al. 1974). In another study of 1206 brains with AML (n=585), ALL (308), chronic granulocytic leukemia (204) and chronic lymphocytic leukemia (109), the CNS involvement was <10% among the total cases, with leptomeningeal lesions ranging from single digits to 50% depending on cancer types (Barcos et al. 1987). A retrospective study of 145 autopsy cases of systemic malignant lymphomas reported CNS invasion of 26.2% of the total or 30.4% of the non-Hodgkin’s lymphomas (Jellinger et al. 1976). Brain autopsy data from patients with MM are limited and appear to have been largely related to leptomeninges (Slager et al. 1979).

Molecular markers are nowadays available for diagnosis of blood cancers. Using the myelogenous cell markers CD68, MPO and lysozyme, we confirmed cancer cell accumulation in meningeal and intracerebral vasculature in 3/9 cases of AML. These markers also detected myelogenous cells inside/around brain blood vessels in the MDS case. Intravascular infiltration of CD20 positive cells was observed in the brain with mantle cell lymphoma. In the three MM cases, we did not detect infiltration of CD138 or MUM1 positive cells in the brain, but observed light chain infusion in the extracellular space and their staining of neuronal somata. The light chain infusion was mild in the 62 y case and occurred largely near the meninge (e.g., layer I) and WM, and was prominent involving the entire parenchyma in the two brains (63 y, 74 y) with Aβ plaques. These findings indicate that blood cancers may cause neuropathological complications through intracranial malignant cell seeding or nesting and “perfusion” of cancer cell derived soluble factors. In fact, the light chain staining of neuronal somata points to a certain neurotoxic impact by the abnormal immunoglobins because the labeled somal profiles appeared to be shrinking, with some crushed into a small remain in an unstained cell space.

The HE and Prussian blue stains revealed frequent presence of vascular injury in the brains with hematological malignancies and cardiovascular diseases. Perivascular edema and microbleeds were found in 4/9 brains with AML, although iron leakage was not detected (Table 1). Vascular damage along with iron leakage was observed in the remaining brains in both disease groups. Overall, vascular injury and iron leakage occurred invariably at the sites of cancer cell infiltration. The cerebral WM, striatum and internal capsule showed the highest regional vulnerability to vascular injury. Importantly, iron leakage could spread into the projecting axonal bundles in addition to the perivascular sites, pointing to a possibility of broad WM damage. The more frequent vascular injury than cancer cell infiltration in the blood cancer group might relate to other factors. Blood cancer patients are treated with various chemotherapeutics, and these drugs not only can affect the amounts of malignant cells in their niche sites and in circulation, but also may cause chemical toxicity to vasculature including in the brain (Ikezoe, 2024). Additional antemortem factors may include the effect of age, and duration and severity of disease between individuals.

In the present study, Aβ deposition appeared primarily in the form of defuse plaques in the brains of the 31 y AML, 87 y lymphoma and the three cardiovascular disease cases. In the brains of 62 y and 74 y MM cases, compact and diffuse plaques coexisted (CAA as well). In comparison, 6E10, APP and BACE1 labeled axonal pathology was observed in the brains with as well as without extracellular Aβ plaques (Table 1). Thus, in most of the cases, the swollen/sprouting axonal processes occurred locally around injured blood vessels in the cerebral and subcortical white matter areas. However, widespread axonal pathology was found in the brains of the 31 y died of AML and 82 y died following SAH, likely representing an acute response.

BACE1 and active γ-secretase binding sites are enriched in neuropil and especially dense at some brain sites abundant of synapses, such as the olfactory glomeruli and hippocampal mossy fiber field (Yan et al., 2004, 2007; Zhang et al., 2009; Cai et al., 2010; Cao et al., 2012; Liu et al., 2013). It appears that synapses may be the major sites of Aβ genesis under physiological conditions. Evidence supports that synaptically released Aβ products deposit locally as extracellular plaques (Lazarov, 2002; Sheng et al., 2002). Aβ deposition can form compact and diffuse plaques, with the former associated with dystrophic neurites, whereas the latter are defined as being lack of these neurites (Armstrong, 1998). Several groups including us have shown that APP, BACE1 and APP β-C-terminal fragments (β-CTFs) accumulate in the dystrophic presynaptic axon terminals of compact plaques (Zhang et al. 2009, Cai et al. 2010; Kandalepas et al., 2013; Sadleir et al. 2016, Jordà-Siquier et al. 2022).

Diffuse Aβ plaques and APP axonal pathology are found in the brains of individuals suffered from cerebral hypoxia/ischemia (Guiroy et al., 1991; Jendroska et al., 1995; Priemer et al., 2022), infarction (Nakina et al., 1992) and traumatic injury (Roberts et al., 1991; Blumbergs et al., 1994; Ikonomovic et al., 2004; Johnson et al., 2010). We carried out a more inclusive appraisal on the stress/injury induced axonal pathology relative to amyloidogenesis in the current study. As assessed between consecutive insula level sections, the APP (Y188) antibody visualized both normal and the pathological axonal processes in the striatum, whereas the APP (22C11), BACE1 and 6E10 antibodies only labeled the latter. The pathological axons were faintly labeled by the 4G8, D54D2 and Ter42 Aβ antibodies, but not visualized by the Amylo-Glo and DANIR-8c Aβ probes. These findings suggest that elevations of APP, BACE1 and β-site cleavage products are a part of the axonal pathology. The interpretation is that the Y188 and 22C11 can detect APP C- and N-terminal fragments (respectively) besides the holo-APP, while the 6E10 can detect APP and β-CTFs besides Aβ products (Yang et al., 2022; Zhang et al., 2024). The faint labeling by other Aβ antibodies (4G8, D54D2 and Ter42) at these axons is most likely related to a cross-reactivity to non-Aβ APP fragments.

Based on the above multi-labeling characterization, we propose a potential link between stress/injury induced axonal pathology and diffuse plaque formation. Specifically, the elevated APP, BACE1and β-CTFs in the injured axonal processes can transport anterogradely to their terminals (Koo et al., 1990; Sheng et al., 2003), leading to an increased synaptic release of Aβ following ɤ-secretase processing. In this context, stress and injury related axonal pathology appear to commonly exist in aging human brains in multiple vulnerable regions (e.g., cortical WM, striatum and internal capsule) and may proceed by cycles of injure/recovery (Garnier-Crussard, 2023; Groh and Simons, 2025; Zheng et al., 2025). As such, white matter axonal pathology with APP, BACE1 and β-CTFs overload may create waves of Aβ release at distant terminal sites, especially in the synapse-rich cortical grey matter, presenting as early formation of diffuse plaques (Johnson et al., 2010).

## Conclusion

Based on initial finding of rare adult cases of tau-independent cerebral β-amyloidosis, we examined a cohort of brains with major types of blood cancers and cardiovascular diseases. Amyloidogenic axonal pathology was prevalent in the studied brains likely due to an inherent vulnerability to vascular injury in both disease conditions. Alzheimer’s disease related clinical and neuropathological research may be benefited from further studies involving large sample-sized elderly with blood cancers.

## ACKNOWLEDGMENTS

We thank Juan Jiang, Chen Yang, Dan-Dan Hu, Qian Li, Qian-Li Shen, Lily Wan for brain banking, and Xiao-Hua Tan for help with Motic light microscopic imaging.

## CONSENT STATEMENT

Written consent for whole body donation for medical education and research was obtained from the donors or next of kin of subjects in compliance with the body/organ donation laws and regulations set by Chinese government. Use of postmortem human brains was approved by the Ethics Committee for Research and Education at Xiangya School of Medicine, in compliance with the Code of Ethics of the World Medical Association (Declaration of Helsinki).

## DATA AVAILABILITY STATEMENT

The raw data supporting the conclusions of this article will be made available by the authors, without undue reservation. Supplementary Figures 1-40 are provided online.

## AUTHOR CONTRIBUTIONS

All authors have read and approved the final version of the manuscript. Conceptualization: XXY, JW; Methodology: YW, PZ, TT, ZPS, XJZ, HPC, HPC, HYC; Clinical data evaluation: ET, JW; Formal analysis and investigation: YW, TT, QLZ, JW; Writing - original draft preparation: YW, QLZ; Writing - review and editing: XXY; Funding acquisition: XXY, JW; Resources: PZ, ET, AP, HYZ, XXY, JW.

