## Supplemental Figures 1-40 for "Early-onset β-amyloidosis in human brains with hematological malignances and cardiovascular diseases: Revisiting injury/stress induced axonal pathology"

### Early-onset β-amyloidosis in human brains with hematological malignances and cardiovascular diseases: Revisiting injury/stress induced axonal pathology

Yan Wang[1], Qi-Lei Zhang[1*], Peng Zhou[2], Tian Tu[1,3], Zhong-Ping Sun[1], Xiao-Jie Zhang[4], Ewen Tu[5], Hui-Ping Chen[6], Hai-Ying Cheng[6], Aihua Pan[1], Jian Wang[6,7*] and Xiao-Xin Yan[1]

[1]Department of Anatomy and Neurobiology, Xiangya School of Basic Medical Sciences, Central South University, Changsha, Hunan 410013, China

[2]Department of Pathology, Second Xiangya Hospital, Central South University, Changsha, Hunan 410031, China

[3]Department of Neurology, Xiangya Hospital, Central South University, Changsha, Hunan 410008, China

[4]Department of Psychiatry, Brain Bank for Psychiatric Disorders, The Second Xiangya Hospital of Central South University, Changsha Hunan 410011, China

[5]Department of Neurology, The Second People's Hospital of Hunan Province, Changsha, Hunan 410007, China

[6]Department of Pathology, Hunan Guangxiu Hospital, Changsha, Hunan 410017, China

[7]Reproductive and Stem Cell Engineering Institute, Xiangya School of Basic Medical Sciences, Central South University, Changsha, Hunan 410013, China

a

### 31 year-old, female, acute myeloid leukemia

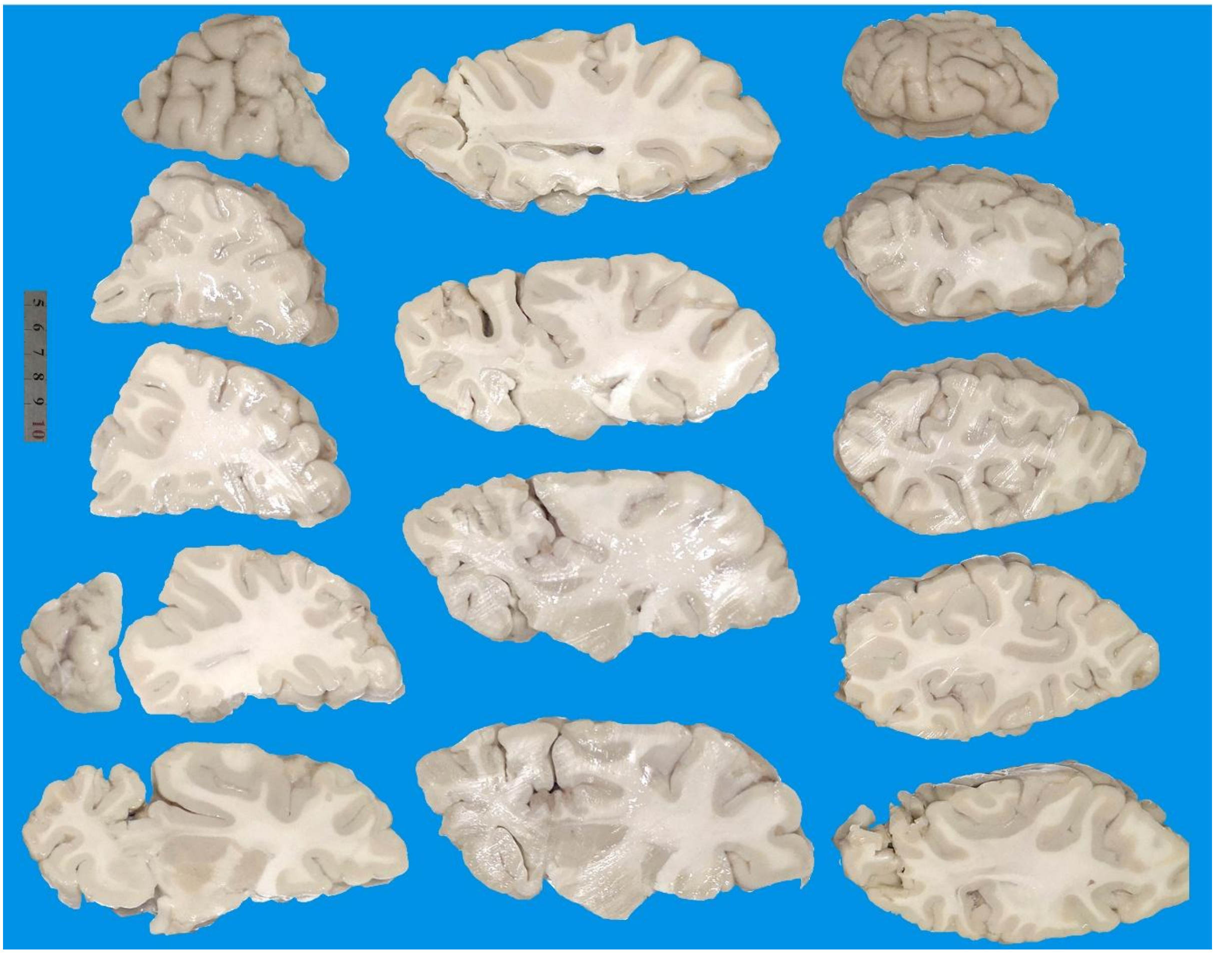

**Supplemental Figure 1. Images of the formalin-fixed brain slices of the 31 year-old case with acute myeloid leukemia.**

By naked eye examination of the slices, there are no macroscopic infarction, hemorrhage or spongiform lesion observed in this brain. The size and shape of the cerebral lobes, hippocampal formation, various subcortical structures and cerebral ventricles appear normal.

**63 years-old, female, multiple myeloma**

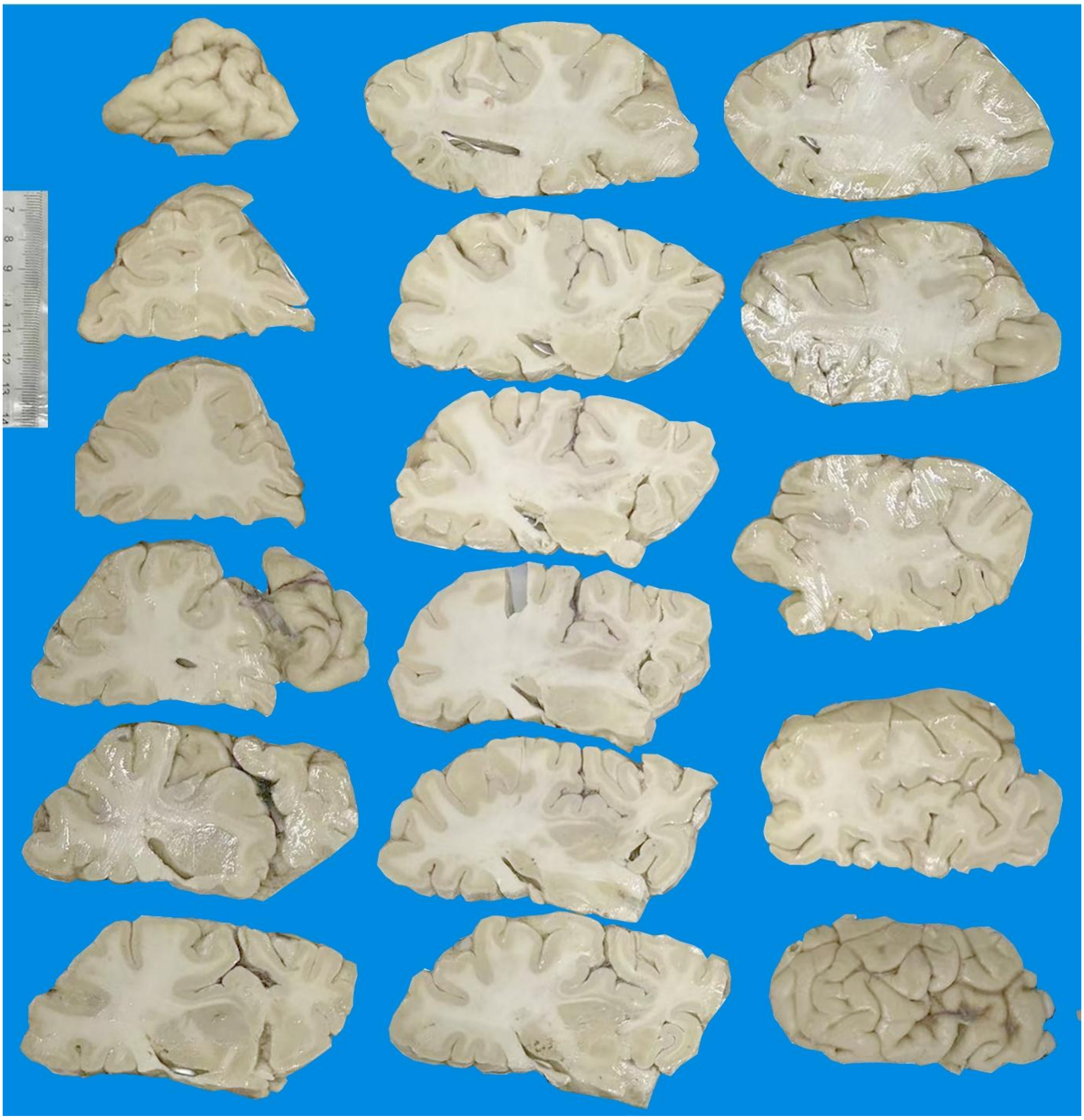

**Supplemental Figure 2. Images of the formalin-fixed brain slices of the 63 year-old case with multiple myeloma.**

No macroscopic infarction, hemorrhage or spongiform lesion are observed in this brain with visual examination of the formalin-fixed slices. The size and shape of the cerebral lobes, hippocampal formation, ventricular system and various subcortical structures appear normal.

### 52 year-old, male, acute myocardial infarction

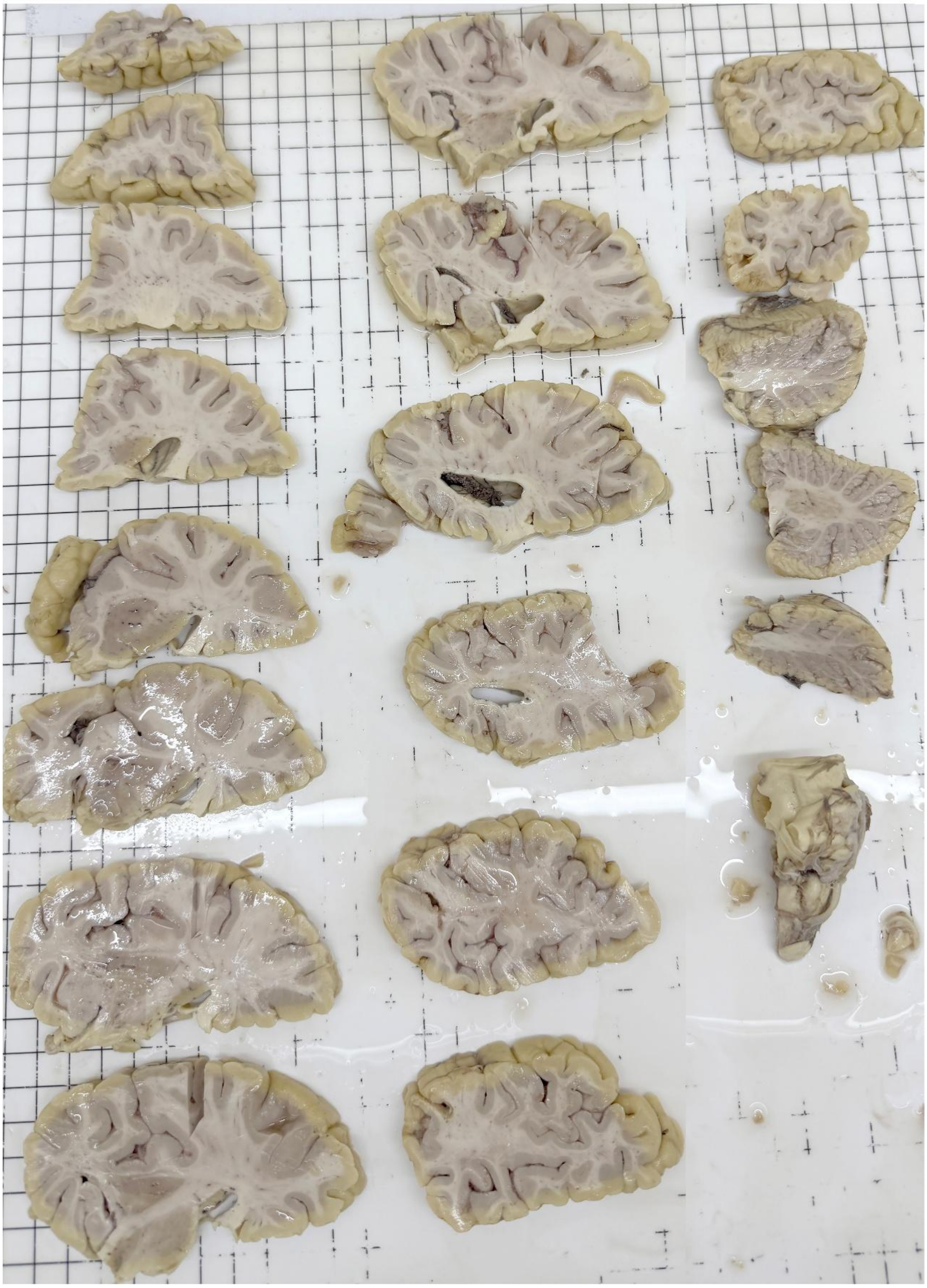

**Supplemental Figure 3. Images of the formalin-fixed brain slices of the 52 year-old case died of an acute myocardial infarction.** No macroscopic infarction, hemorrhage or spongiform lesion are observed in this brain by visual examination of the formalin-fixed slices. The size and shape of the cerebral lobes, hippocampal formation, ventricles and various subcortical structures appear normal. There are some darkened spots in the white matter and basal ganglia.

**65 years-old, male, chronic cerebral ischemia**

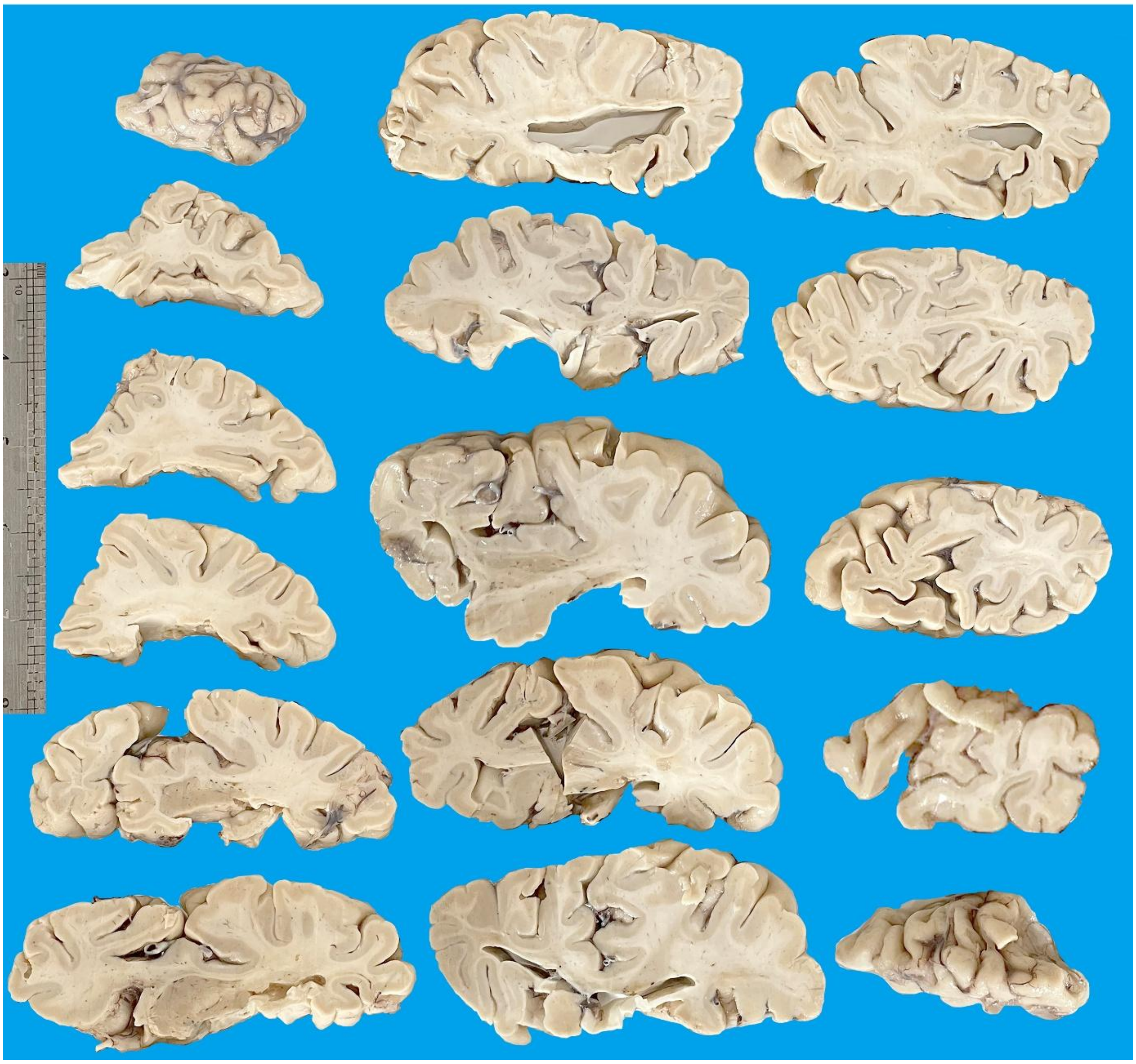

**Supplemental Figure 4. Images of the formalin-fixed brain slices of the 65 year-old donor suffered from chronic cerebral ischemia for approximately 5 years.** No macroscopic infarction, hemorrhage or spongiform lesion are observed in the formalin-fixed brain slices. The size and shape of the cerebral lobes, hippocampal formation, ventricles and various subcortical structures appear to be in normal range. There are some darkened spots resembling vascular profiles in the white matter and basal ganglia.

87 years-old, male, lymphoma

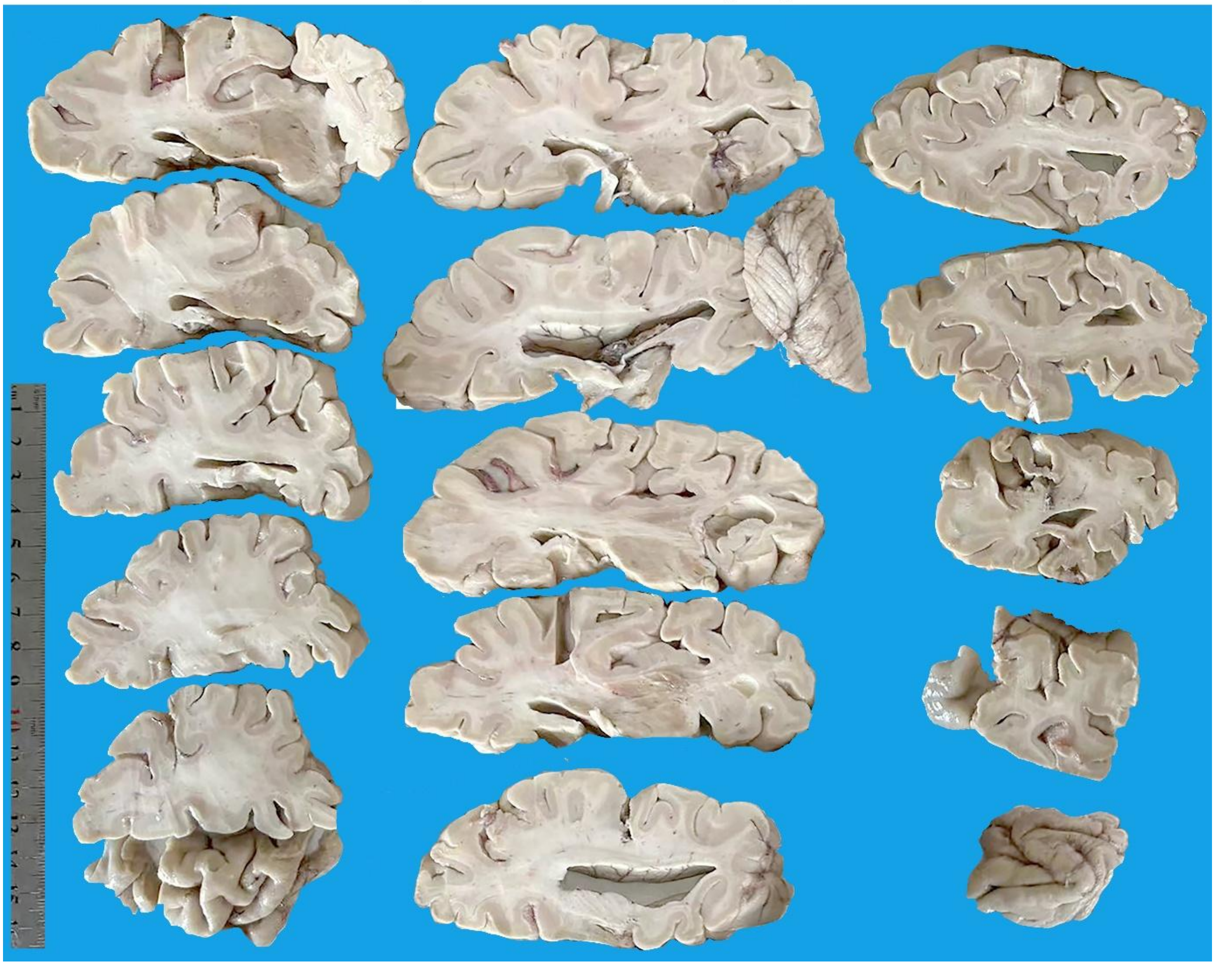

**Supplemental Figure 5. Images of the formalin-fixed brain slices of the 87 year-old donor who was clinically diagnosed with lymphoma.** No macroscopic infarction, hemorrhage or spongiform lesion are observed in the formalin-fixed brain slices. The size and shape of the cerebral lobes, hippocampal formation and various subcortical structures appear to be in normal range. The lateral ventricle appears to be somewhat enlarged. There are a few darkened spots resembling vascular profiles in the white matter and basal ganglia.

**82 y, female, hypertension, coronary heart disease, subarachnoid hemorrhage**

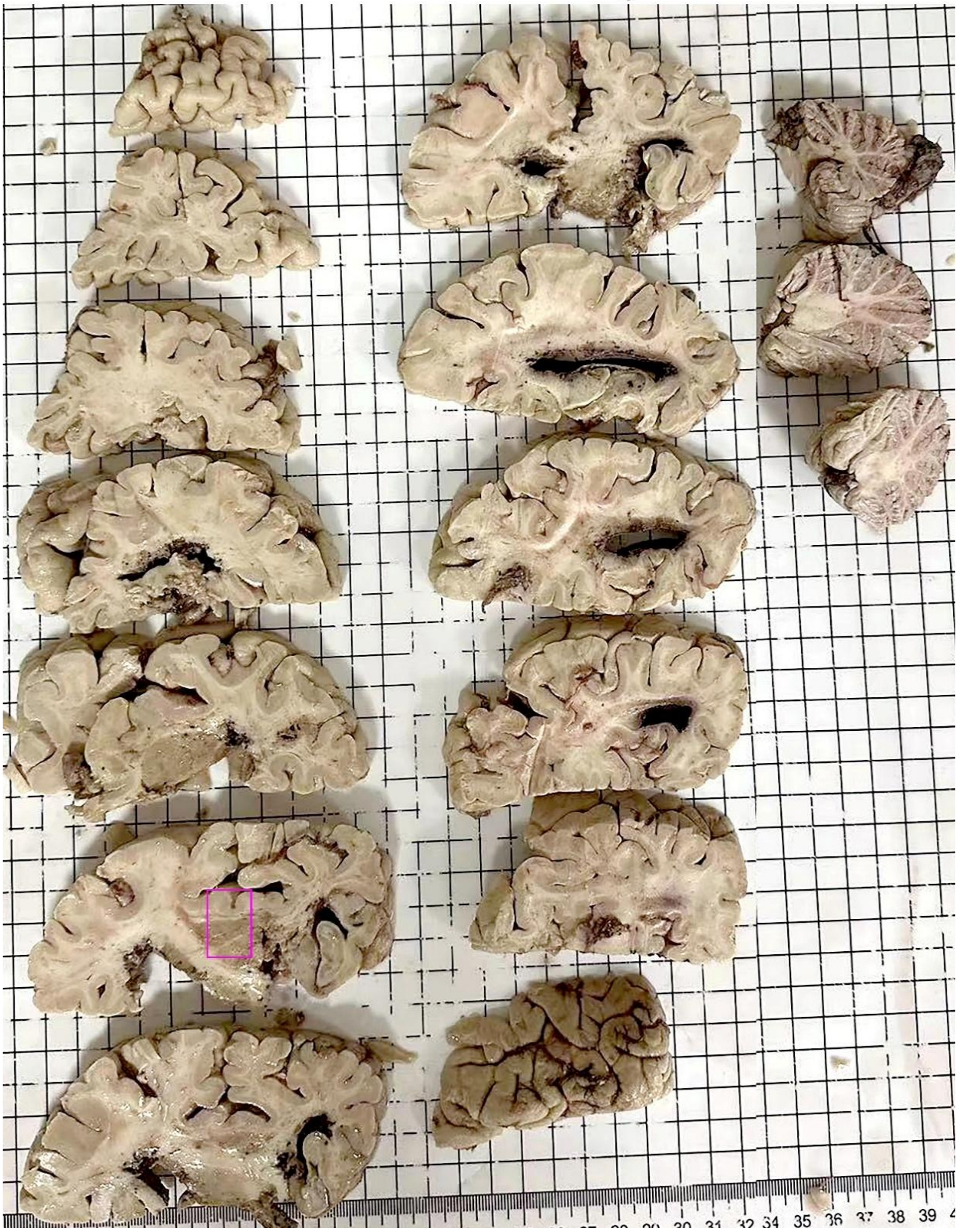

**Supplemental Figure 6. Images of the formalin-fixed brain slices of the 82 year-old donor who had a clinical record of hypertension and coronary heart diseases and died following an acute attack of subarachnoid hemorrhage.** Blood was filled in the ventricular system at brain collection. In the formalin-fixed slices, the cerebral lobes, hippocampal formation, the basal ganglia and other subcortical structures can be recognized. However, structures around the ventricles are damaged to various extents due to the blood accumulation in cerebral ventricles. Darkened areas are seen in the white matter and basal ganglia. The framed area is sampled for pathological study in the insular and striatal areas.

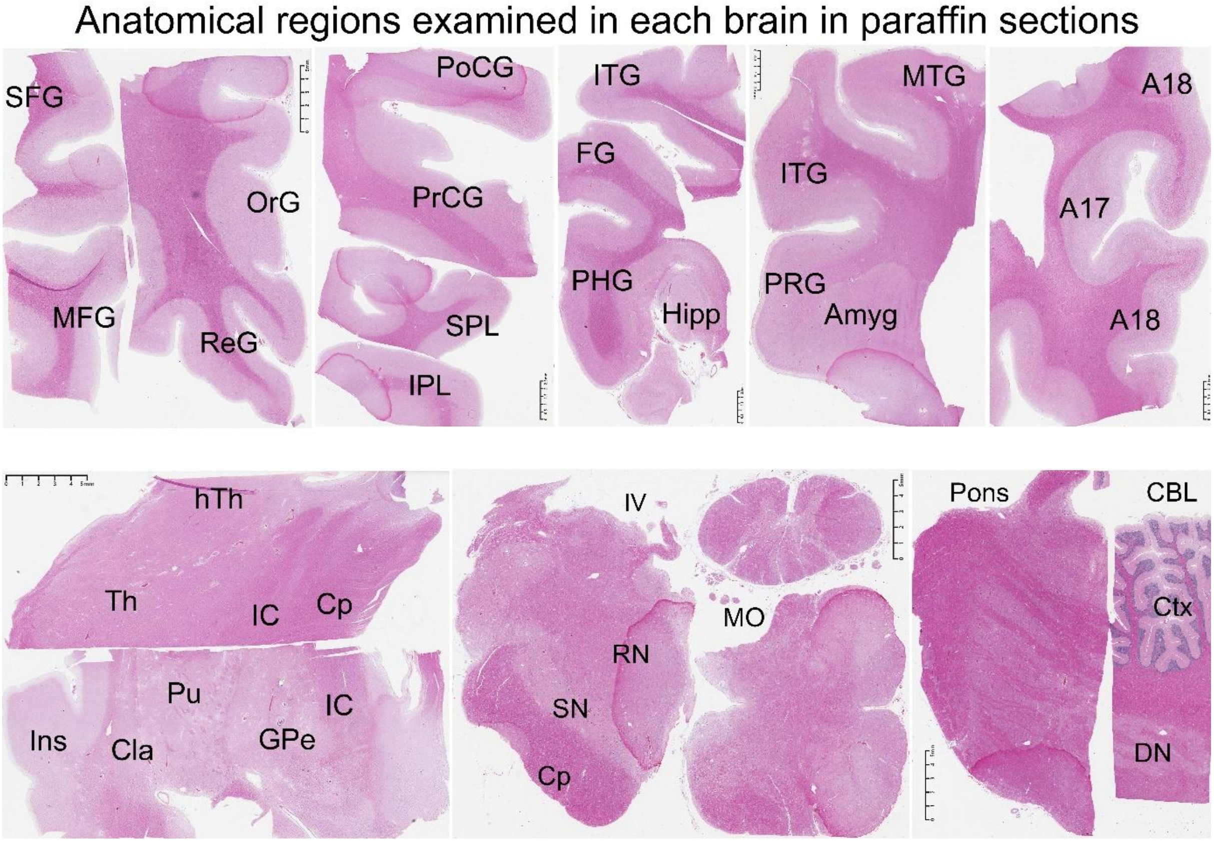


**Supplemental Figure 7. Hematoxylin and eosin (HE) stained sections showing the examined cerebral regions and subcortical structures for postmortem human brain study according to the Standard Operation Protocol set forth by the China Brain Banking Consortium.** The current study focuses on neuropathological characterization in the regions with microscopically evident vascular injury, iron leakage, amyloidogenic axonal pathology, along with β-amyloid deposition and tau pathology whenever present, in a given brain among all the cases in the study cohort.

Abbreviations for neuroanatomical structures labeled in this and other Supplemental Figures are listed below according to brain regions. (1) General: Ctx: cortex, I-VI: cortical layers, WM: white matter; LV, 3V and 4V: cerebral ventricles. (2) Frontal cortex: SFG: superior frontal gyrus, MFG: middle frontal gyrus, RoG: rostral cingulate gyrus; ReG: rectal gyrus, OrG: orbital gyrus, AOrG, MOrG, LorG: anterior, medial and lateral orbital gyri. (3) Primary motor and sensory cortex: PoCG: postcentral gyrus, PrCG: precentral gyrus. (4) Parietal cortex: SPL and IPL: superior and inferior parietal lobules. (5) Temporal lobe structures: MTG and ITG: middle and inferior temporal gyri, FG: fusiform gyrus, PHG: parahippocampal gyrus, Ent: entorhinal cortex, PRG: perirhinal gyrus, Para-S, Pre-S, Sub and Pro-S: subicular subregions, Hipp: hippocampus; Amyg: amygdalar complex; CA1-3: Ammon's horn subregions, s.o., s.p. and s.l.m.: strata oriens, pyramidale and lacunosum-moleculare, DG: dentate gyrus, ML: molecular layer, GCL: granule cell layer: GCL, Hi: hilus, PP: perforant path, Alv: alveus, MF/mf: mossy fiber, col-s: collateral sulcus. (6) Occipital cortex: A17: the primary visual cortex, A18: the secondary visual cortex, cs: calcarine sulcus. (7) Insular level structures: Ins: insular cortex, Cla: claustrum, EC: external capsule, Pu: putamen, GPe and GPi: the external and internal parts of globulus pallidum; Th: thalamus; hTh: hypothalamus; IC: internal capsule; Cp: cerebral peduncle; VA: ventral anterior nuclei, MD: mediodorsal nuclei, PHN: posterior hypothalamic nucleus, LHN: lateral hypothalamic nucleus. (8) Midbrain: RN: red nucleus; SN: substantia nigra; Brainstem: MO: medulla oblongata; (9) Cerebellum: CBL: cerebellum; Ctx: cortex; DN: dentate nucleus.

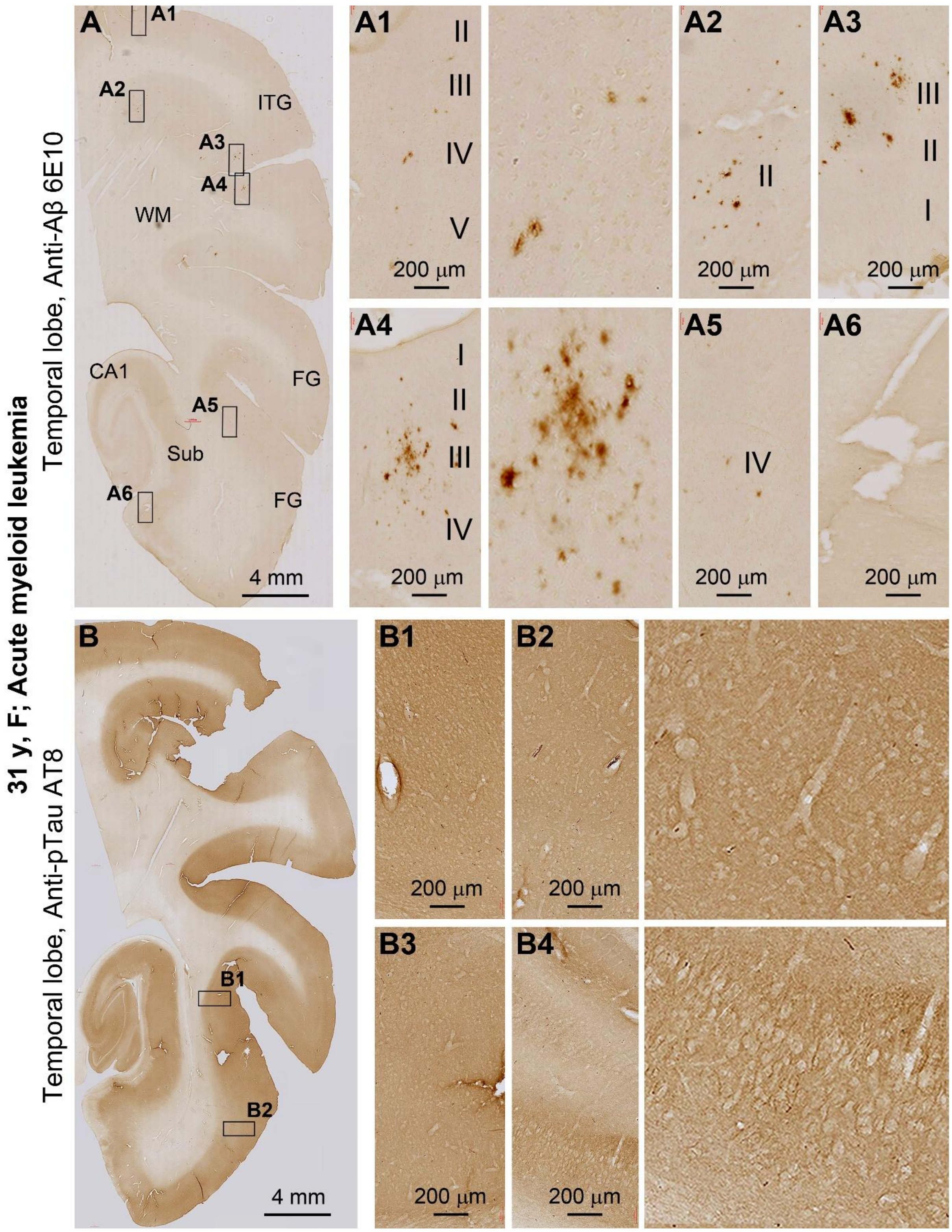


**Supplemental Figure 8. Characterization of β-amyloid and tau pathologies in the temporal lobe structures in the 31 year-old case with acute myeloid leukemia.** Panels (**A**) and enlarged areas show small amounts of extracellular Aβ deposition appearing as diffuse plaques labeled by the 6E10 antibody. These plaques are present in the temporal lobe neocortical regions including the inferior temporal gyrus (ITG) (**A1, A2, A3, A4**) and fusiform gyrus (FG) (**A5**). They are not detected in the hippocampal subregions (**A6**). (**B**) and enlarged panels show the lack of neuronal tauopathy with the AT8 antibody labeling.

31 y, F; Acute myeloid leukemia; insula, striatum, thalamus

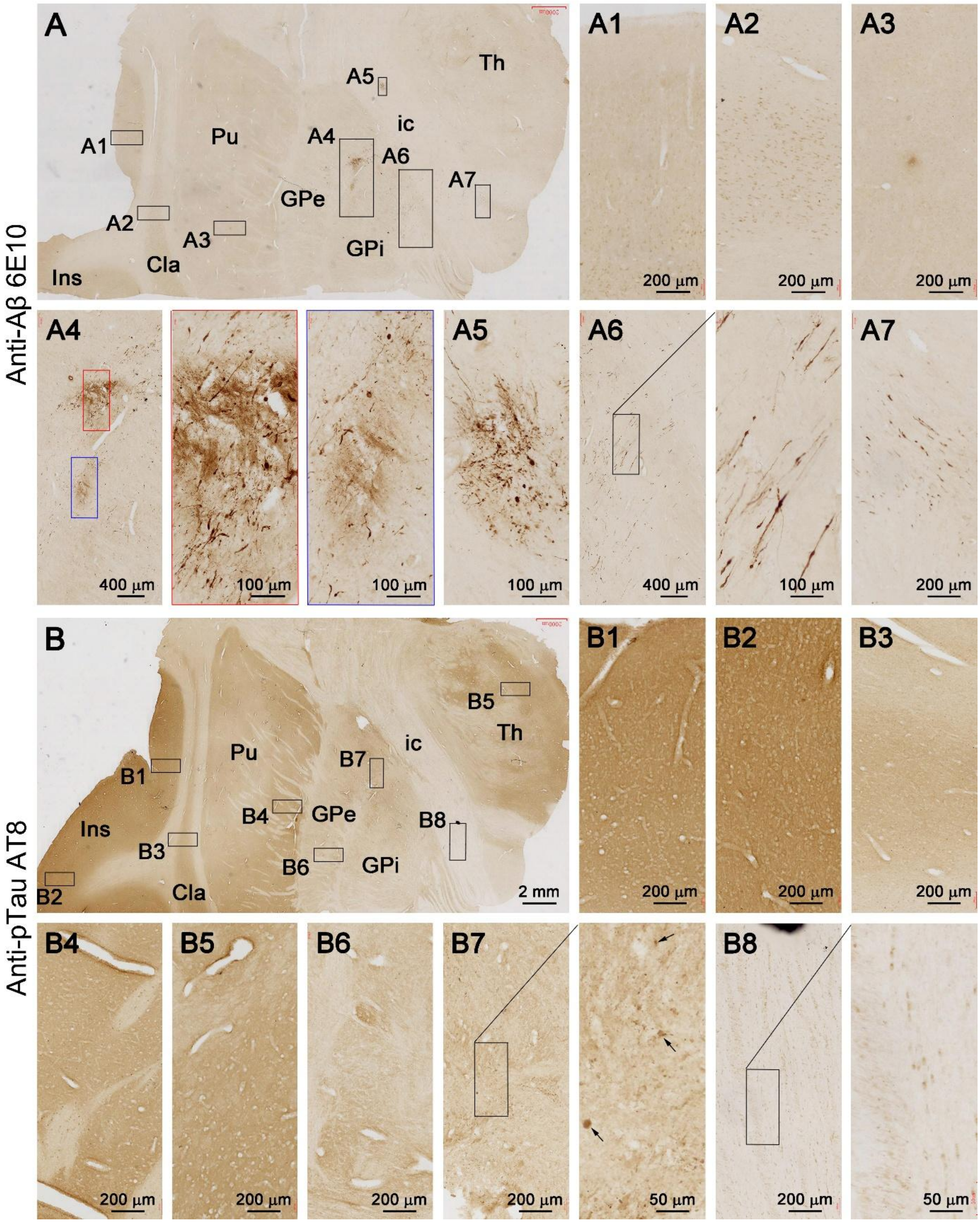


**Supplemental Figure 9. 6E10 and AT8 antibody labeling in striatum level sections in the 31 year-old case with acute myeloid leukemia.** No Aβ plaques are labeled in the insular cortex, claustrum or striatum (**A, A1-A3**). However, densely packed and isolated swollen axons are present in globulus pallidum (GP) and internal capsule (IC), with the former associated with local extracellular Aβ deposition (**A, A4-A7**). No neuronal or neuritic elements are labeled by the phosphorylated tau (pTau) AT8 antibody across the section (**B, B1-B8**) except for occasional spheroids (arrows) in the striatal areas with axonal pathology (**B7**).

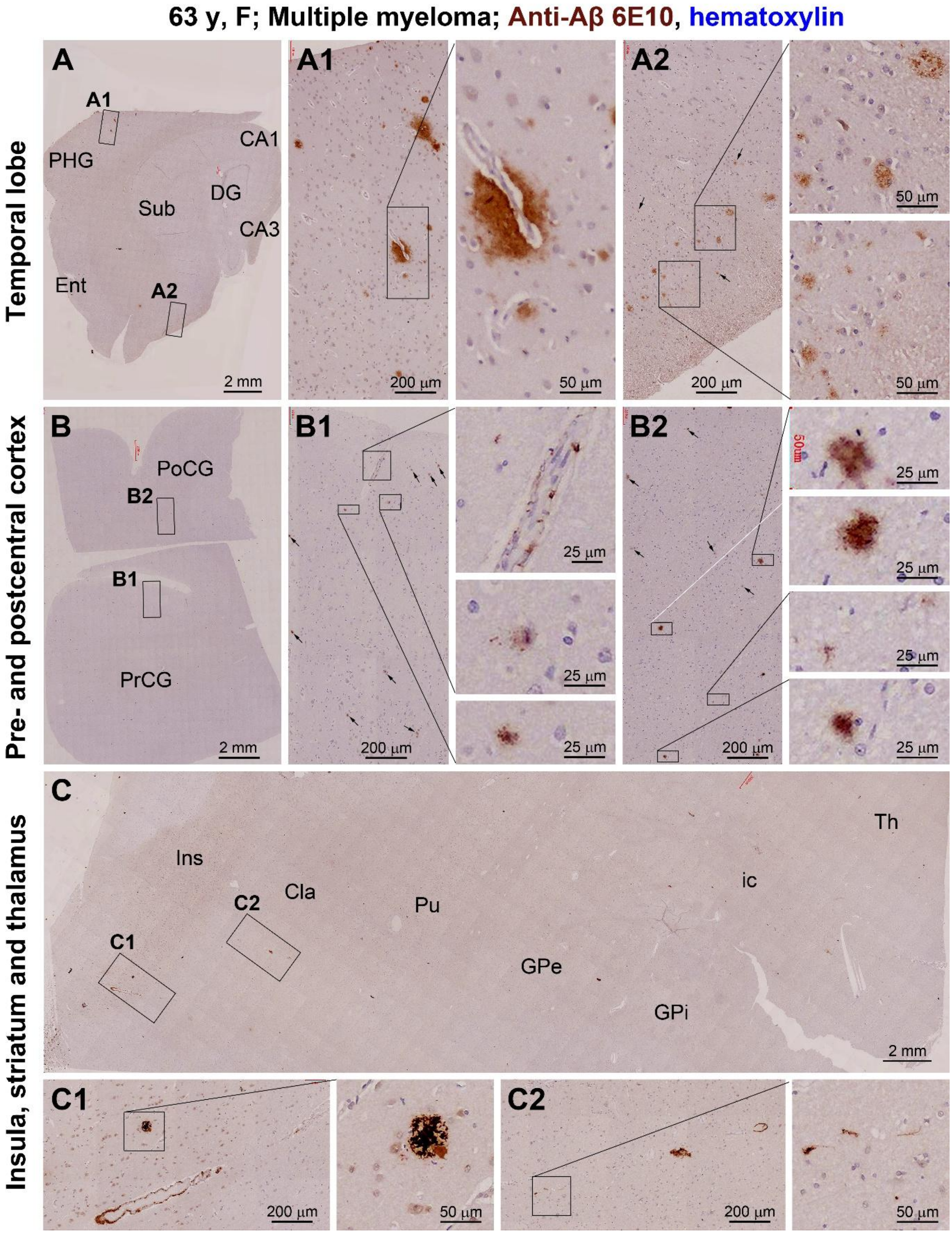


**Supplemental Figure 10. 6E10 labeled Aβ deposition in multiple brain regions in the 63 y case of multiple myeloma.** Diffuse and compact plaques as well as cerebral amyloid angiopathy (CAA) are found in temporal lobe (**A, A1, A2**), post- and precentral gyri (**B, B1, B2**), insular cortex, claustrum and striatum (**C, C1, C2**). The overall burden of the plagues is low in the regions. Very small plaques (pointed by arrows) have a fibrillary configuration (**B1, B2**). The CAA also shows a varying pattern, from isolated immunolabeling aligning with the epithelium (**B1**) to continuous Aβ deposition along the vascular wall (**C1**).

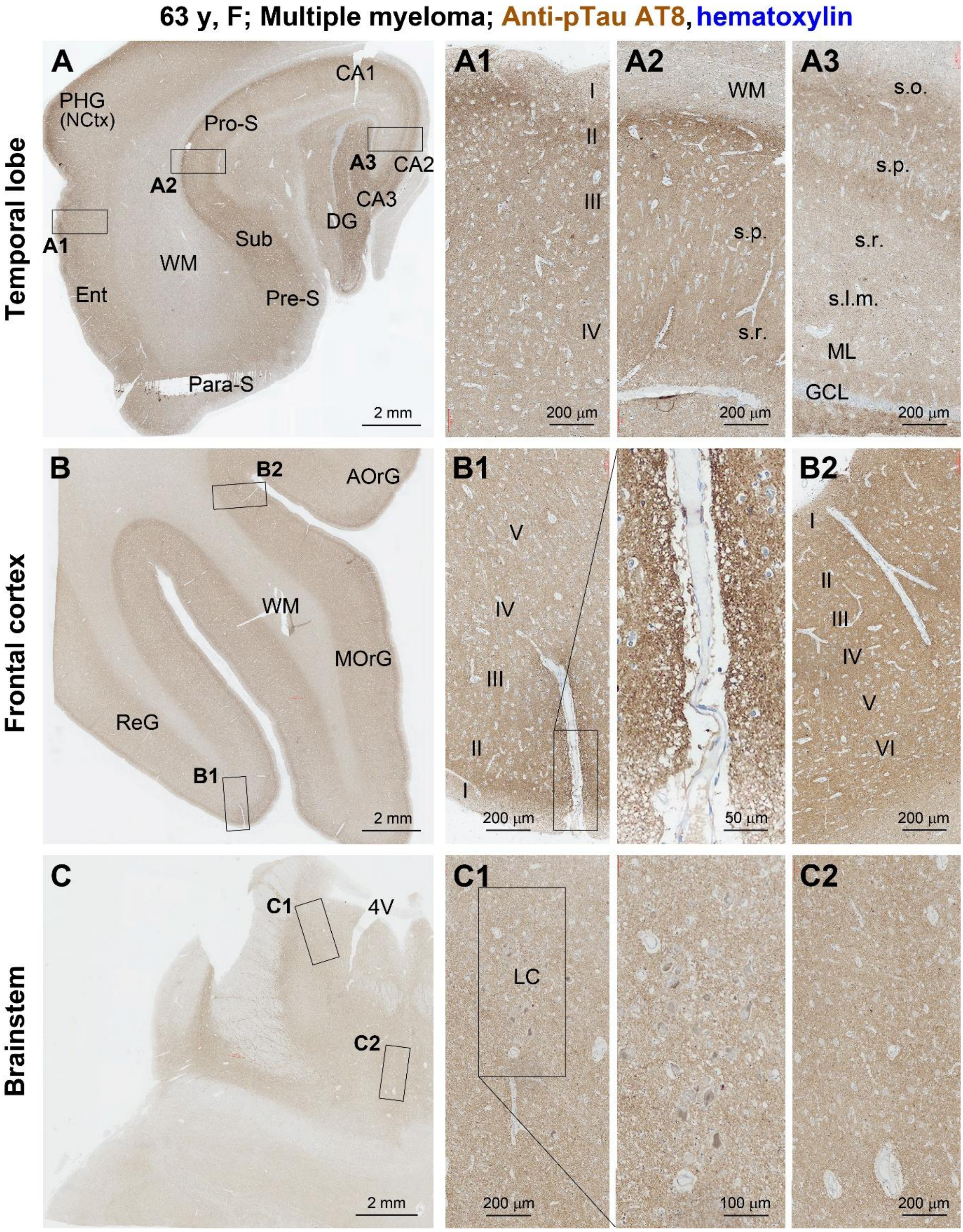


**Supplemental Figure 11. Lack of AT8 immunolabeling in representative brain regions in the 63 y case of multiple myeloma.** In the temporal lobe section, no pTau positive somata or neuritic profiles are found in the entorhinal cortex, subiculum and CA subregions (**A, A1, A2, A3**). No labeled cells are seen in the frontal cortex, although the labeling appears somewhat denser surrounding blood vessels with increased perivascular space (**B, B1, B2**). No pTau positive neurons are present in the locus coeruleus in the brainstem section (**C, C1, C2**), a brain region considered to exhibit the earliest tauopathy according to Braak staging.

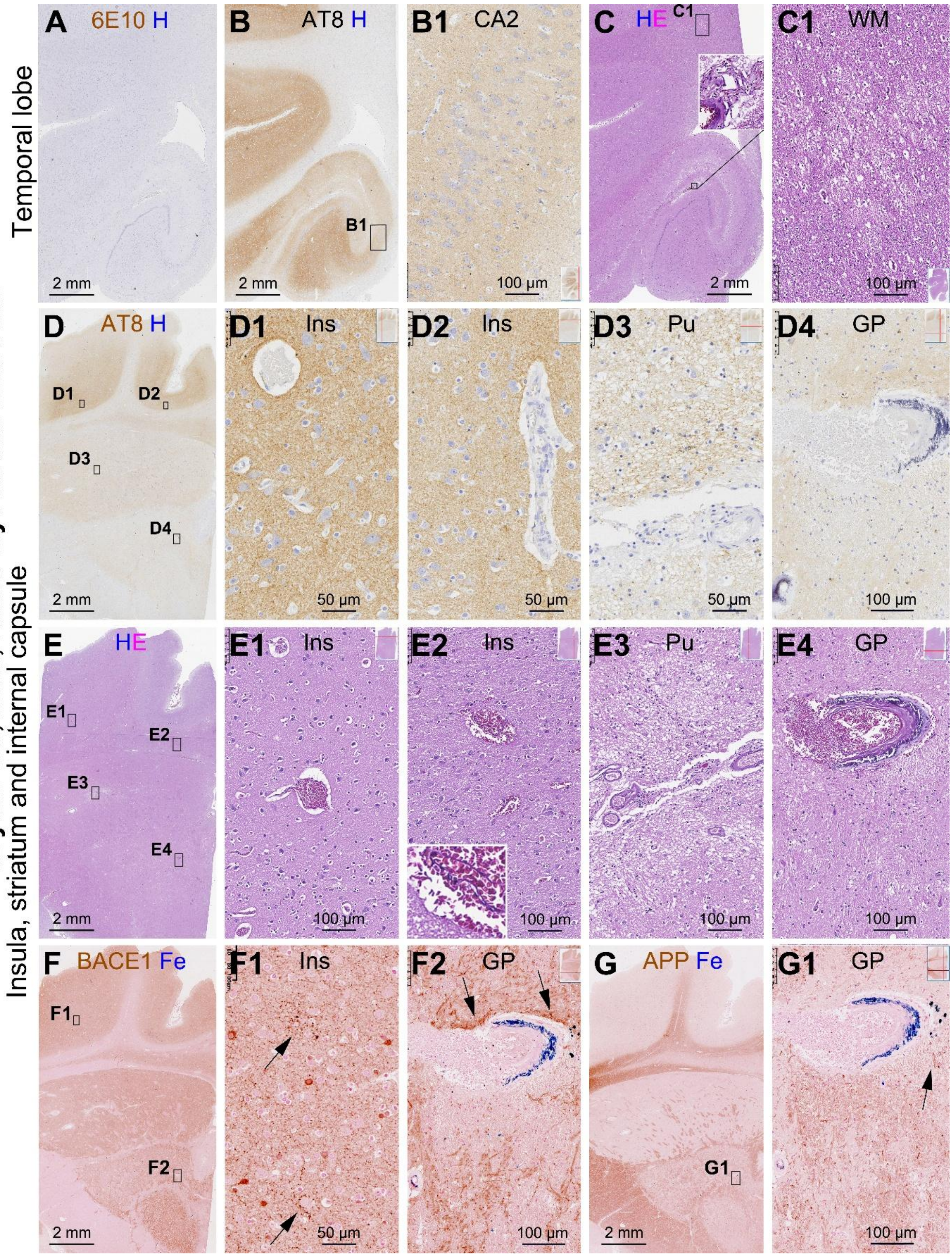


**Supplemental Figure 12. Multi-staining pathological assessment of the brain from the 52 y case died following acute myocardial infarct.** No Aβ deposition and tau pathology are found in the temporal lobe and insular level sections (**A, B, B1, D, D1-D4**). Localized white matter damage (**C, C1**), increased perivascular space (**D, D1-D4, E, E1-E3**), microbleed (**C, E2, E4**), vascular wall calcification (**D4, E4**) and iron leakage (**F2, G2**) are present. Small amounts of swollen axons (arrows) are labeled by the BACE1 and APP antibodies in the insular cortex and striatum, occurring largely around damaged blood vessels (**F, F1, F2, G, G1**).

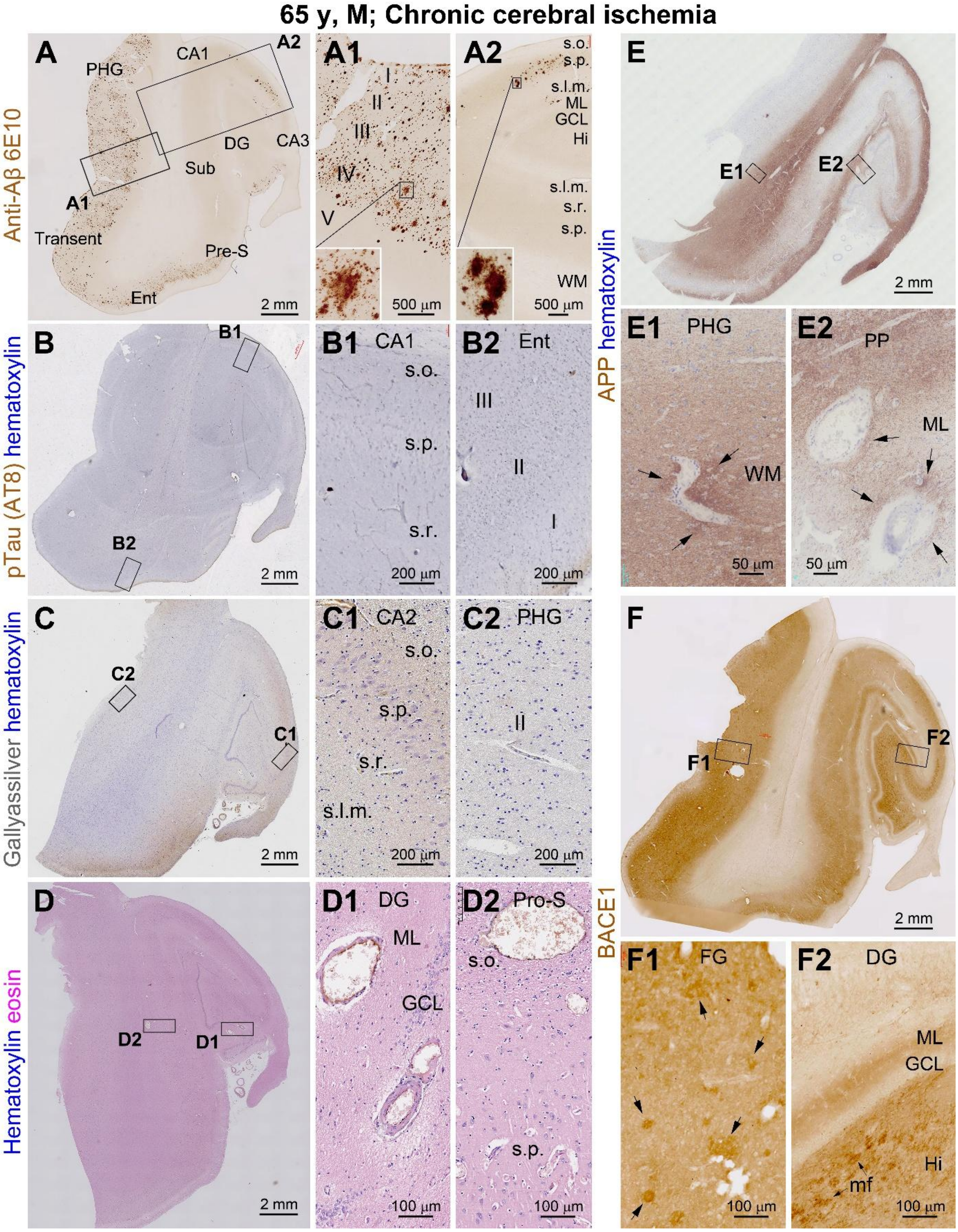


**Supplemental Figure 13. Multi-stain assessment in temporal lobe sections from the 65 y case with chronic cerebral ischemia.** Diffuse Aβ plaques are loaded in the temporal neocortex and entorhinal cortex but only occur in CA2 and the molecular layer of dentate gyrus in the hippocampus (**A, A1, A2**). No pTau and Gallyas labeling are present (**B, B1, B2, C, C1, C2**). Vascular wall thinning and perivascular edema are present (**D, D1, D2**). APP axonal labeling appears denser around blood vessels (**E, E1, E2**). BACE1 labeling exhibits patch-like increase in the cortex (**F, F1**), with normal mossy fiber labeling (**F, F2**).

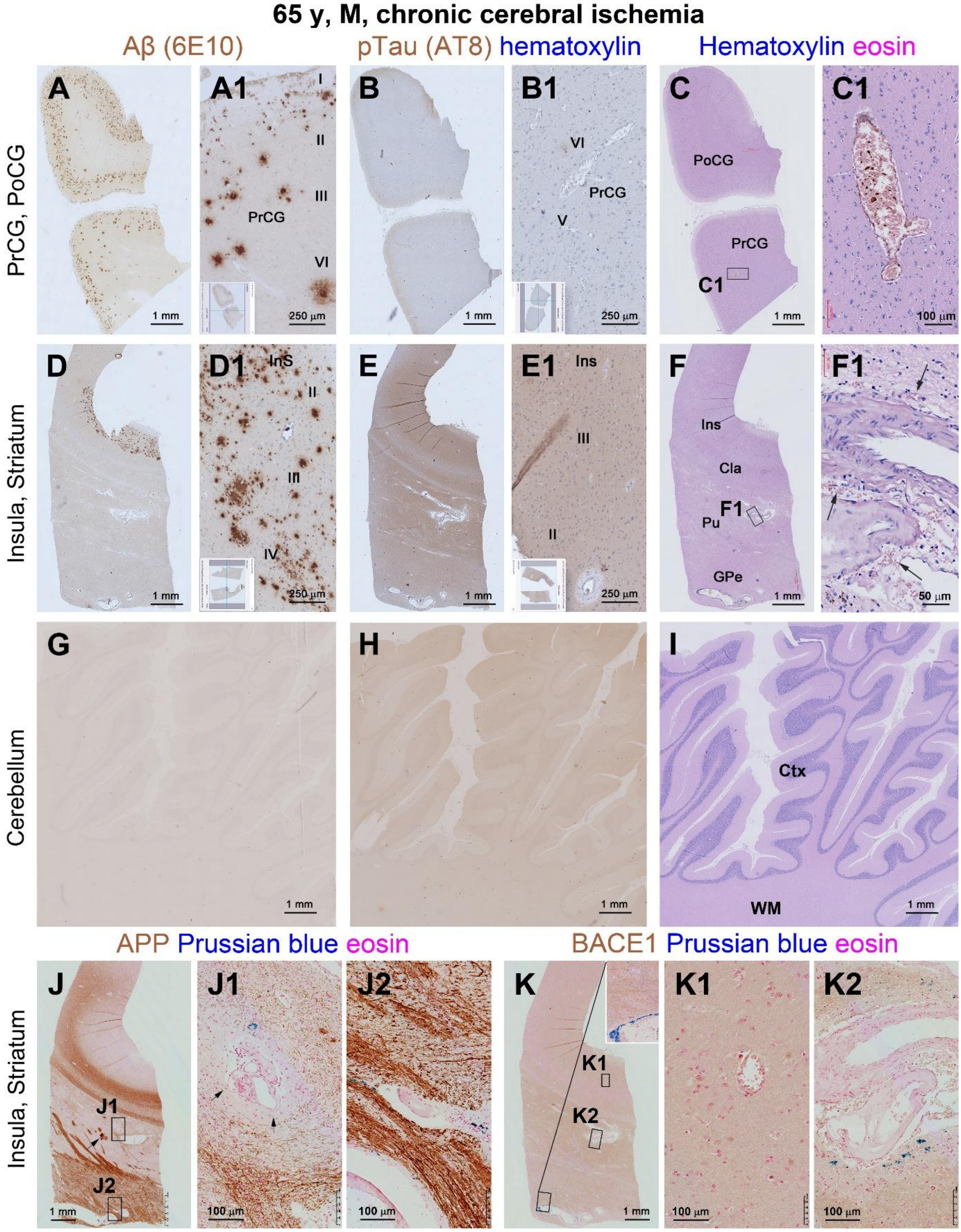


**Supplemental Figure 14. Multi-stain assessment in additional brain regions in the 65 y case with chronic cerebral ischemia.** Diffuse Aβ plaques are packed in the motor, sensory and insular cortex, but absent in striatum and cerebellum, with no tauopathy found in these areas (**A, B, D, E, G-H** and high power views). Vascular injury is seen in HE stain (**C, C1, F, F1**). APP labeling appears heavy in the striatum, with swollen axons (arrow) around the blood vessels with iron deposition (**J, J1, J2**). These axonal changes are not clearly labeled with BACE1 (**K, K1, K2**).

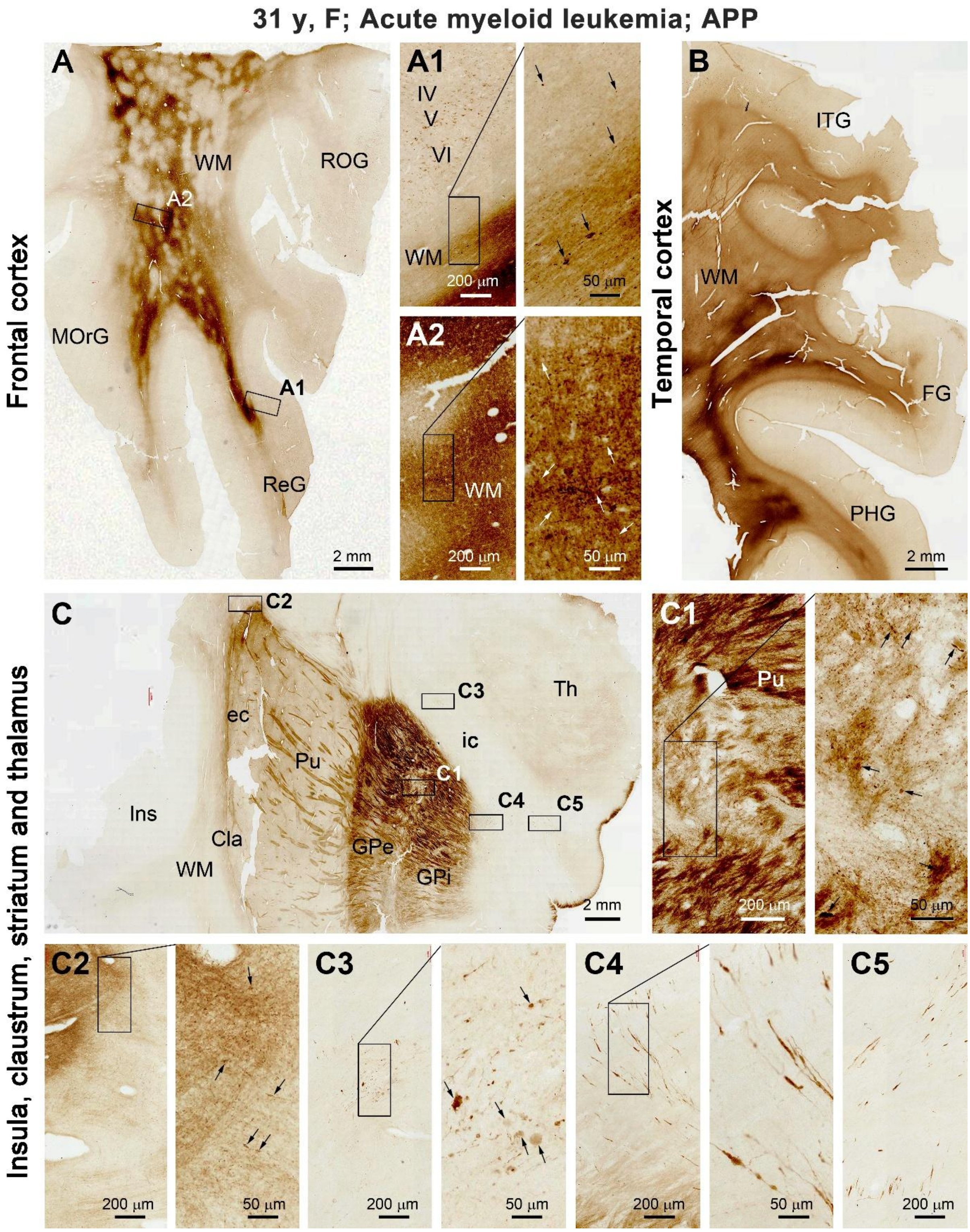


**Supplemental Figure 15. APP labeled white matter damage and axonal pathology in the 31 y case of AML.** In both the frontal (**A, A1**) and temporal (**B**) cortex, the labeling is uneven in the white matter, with swollen axonal processes seen at high magnification in these regions as well as in deep cortical layers (**A1, A2**). In the striatum level section (**C**), the axonal bundles are locally disrupted (**C1**). Swollen axonal fibers are seen in insular white matter (**C, C2**). The swollen axonal processes extend in the internal capsule. These processes and large axonal spheroids are also packed locally as clusters (**C, C3-C5**).

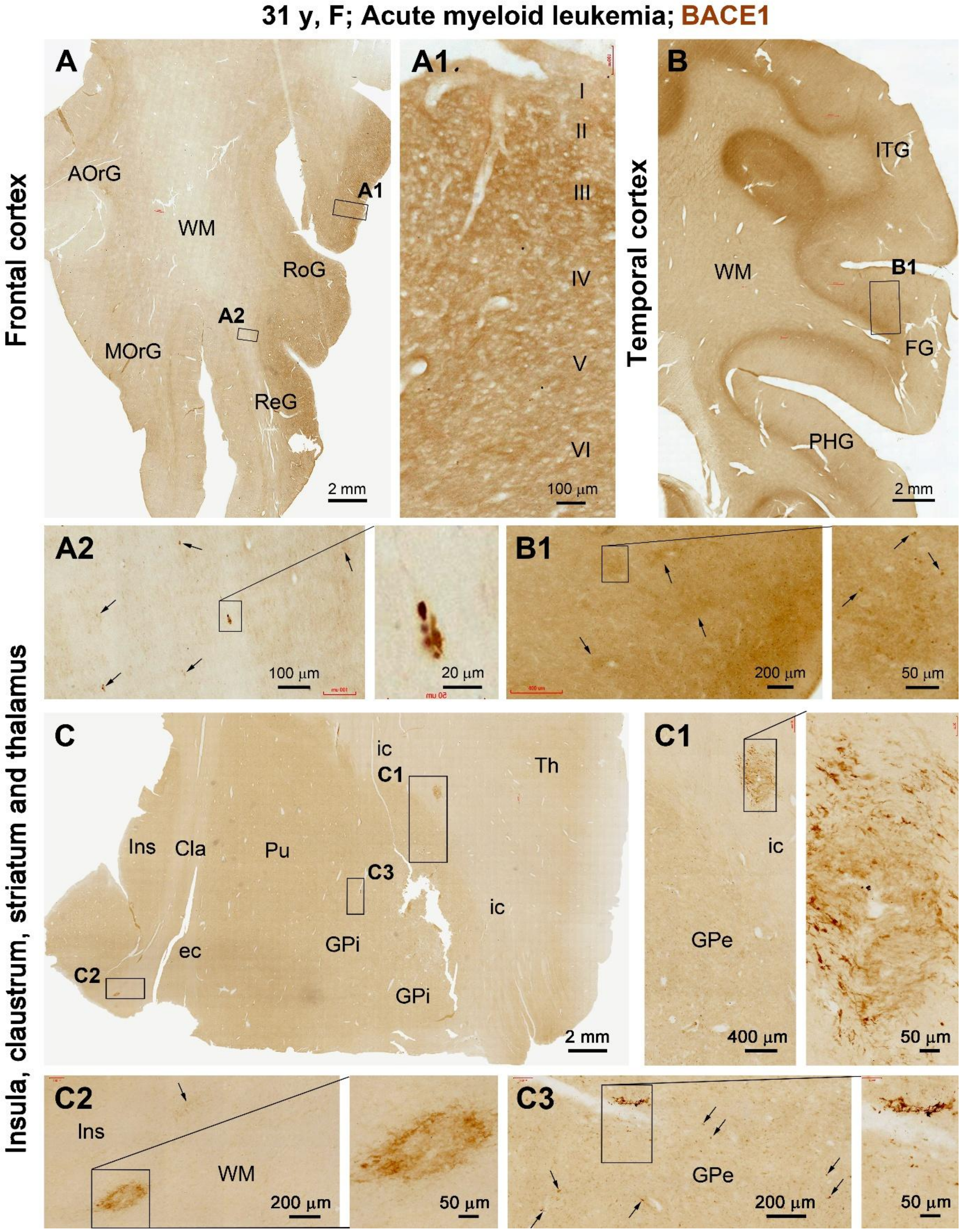


**Supplemental Figure 16. BACE1 immunolabeling in representative neocortical regions and in the striatum level section in the 31 y case of AML.** The labeling in the cortical grey matter exhibits largely a normal-looking neuropil pattern (**A, A1, B, B1**) except for slightly enhanced patch-like reactivity and small spheroids (**A1, B1**), However, strongly labeled swollen axonal processes (arrows) are seen in the white matter (**A2**). The swollen axonal neurites, clusters and axonal spheroids are clearly present in the striatum and internal capsule, occur also near blood vessels (**C, C1, C2, C3**).

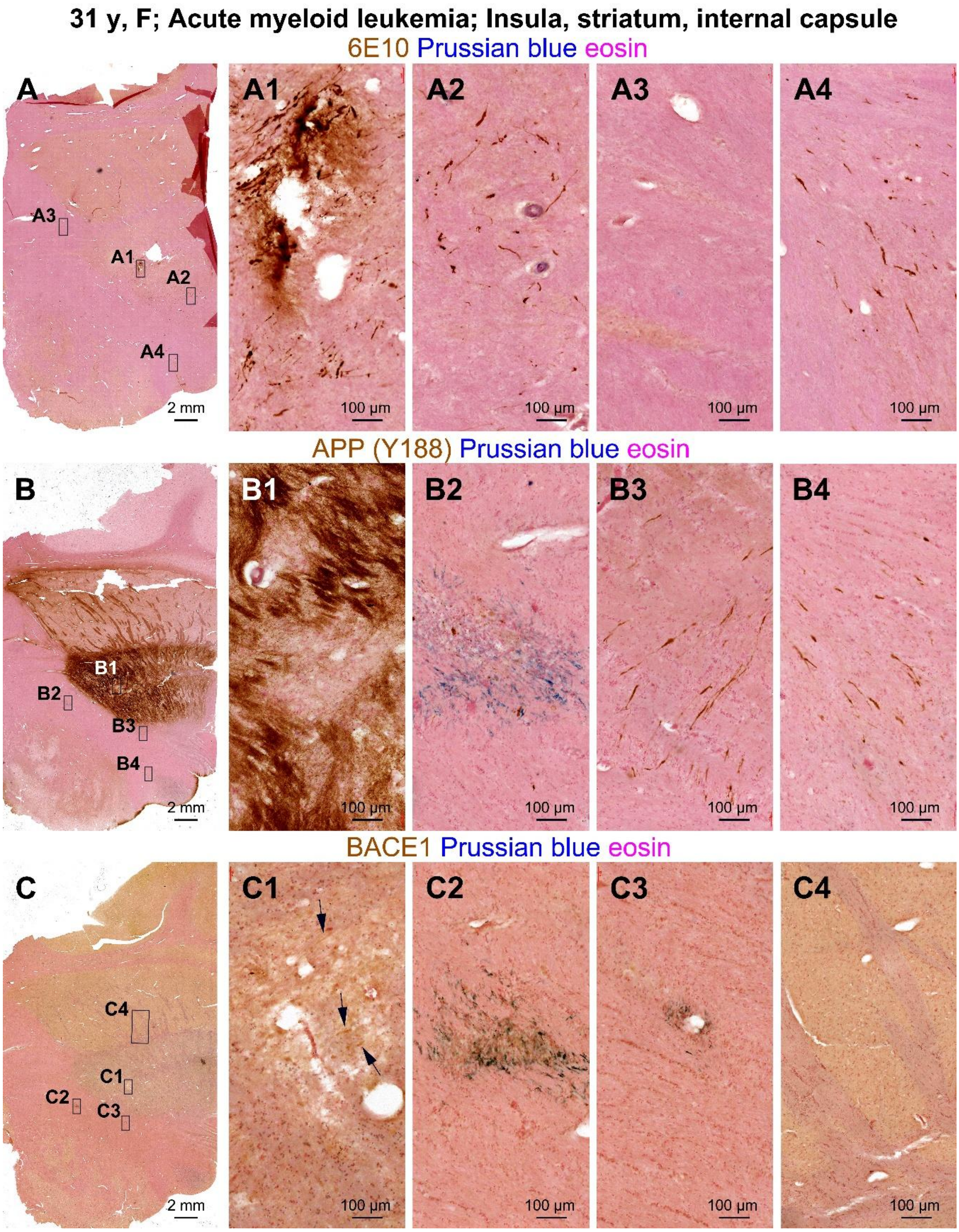


**Supplemental Figure 17. Assessing the relationship between vascular injury and axonal pathology in the striatum level sections from the 31 y AML case.** The staining and panel arrangement are as indicated, with subregions in (**A, B, C**) also illustrated in main Figure 3. Extracellular Aβ deposition is present in a striatal area where the APP and BACE1 labeling are reduced, while some swollen axons (arrows) remain (C1). In the internal capsule, APP and BACE1 labeled neuritic clusters are associated with iron deposition, while other swollen axons extend individually. There is no spread iron infiltration in the region including in the axonal fiber bundles in this blood cancer case.

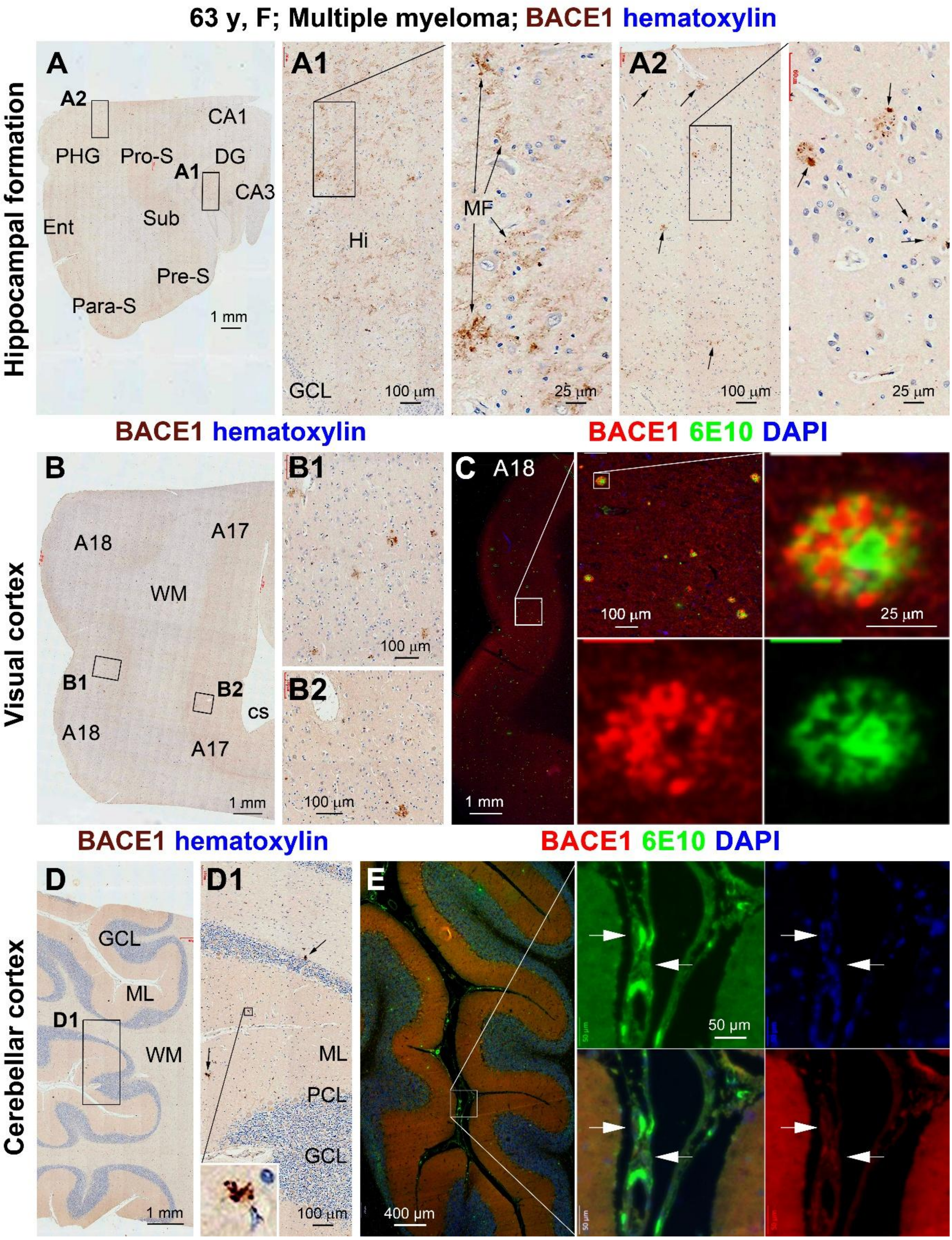


**Supplemental Figure 18. BACE1 immunolabeling in dystrophic neurites and blood vessels and its relevance to Aβ deposition in the 63 y case with multiple myeloma.** BACE1 labeling in the hippocampal mossy fiber (MF) terminals is distinct, with some exhibiting a dystrophic appearance (**A, A1**). Neuritic clusters (arrows) are present in the temporal (**A, A2**) and visual cortex (**B, B1**) and in the cerebellar cortex (**D, D1**). (**C, E**) show BACE1 and 6E10 double immunofluorescence at compact neuritic plaques in the visual cortex, and in pial vessels of the cerebellar cortex that appears to be in the endothelial cells (arrows).

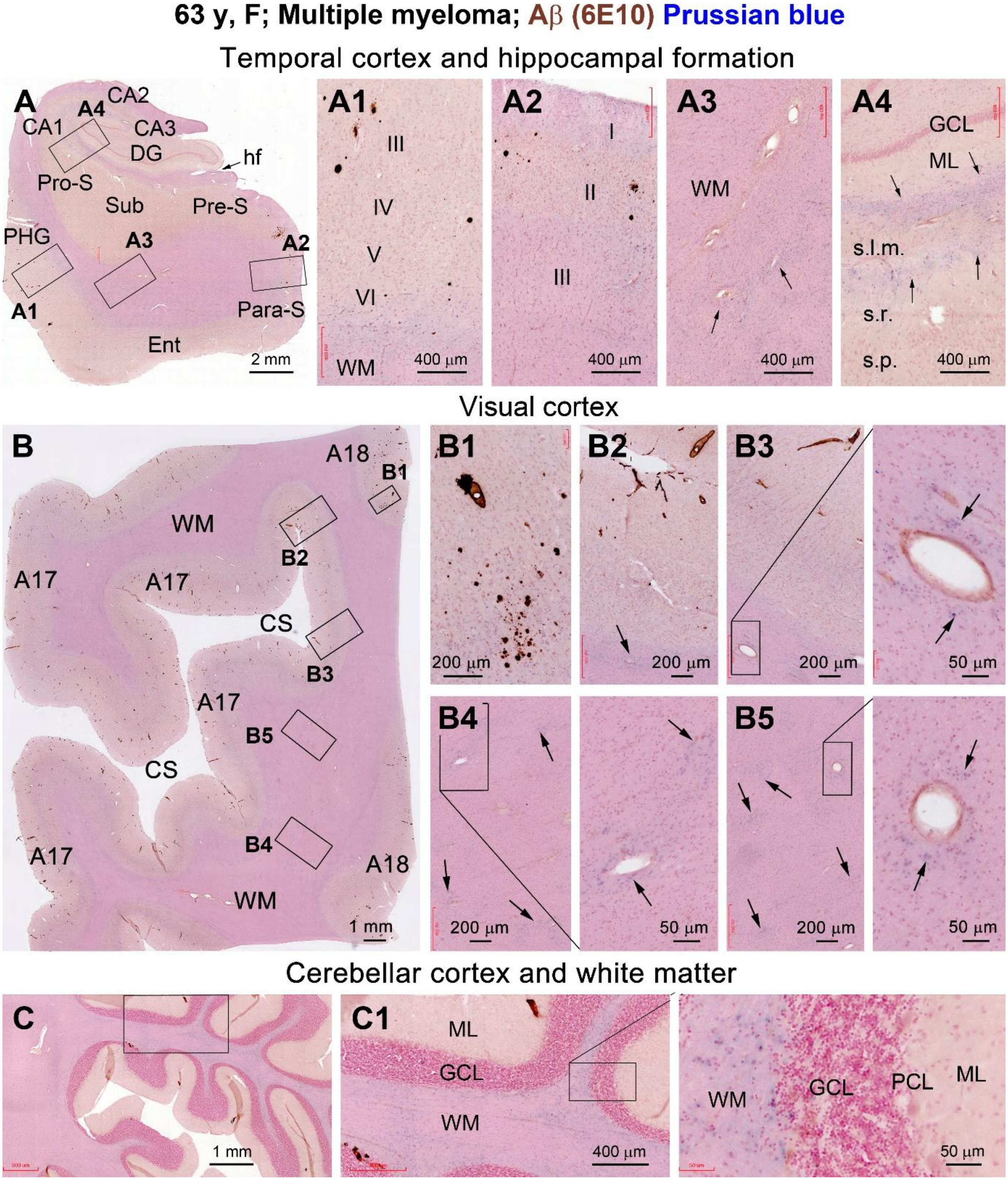


**Supplemental Figure 19. Vascular injury assessed in 6E10 immunolabeled sections with Prussian blue counterstain in representative brain regions in the 63 y multiple myeloma case.** In the temporal lobe section (**A**), Aβ plaques and CAA are present in the temporal neocortex (**A1**) and entorhinal cortex (**A2**), but not in the hippocampal formation (**A4**). These lesions are also present in the primary (A17) and secondary (A18) visual cortex and cerebellar cortex (**C, C1**). Iron deposition (pointed by arrows) is seen in the white matter of cerebral and cerebellar cortex (**A3, B, B3-5, C1**) and the stratum lacunosum-moleculare (s.l.m.) of hippocampus (**A4**), which is noticeably heavier around the blood vessels.

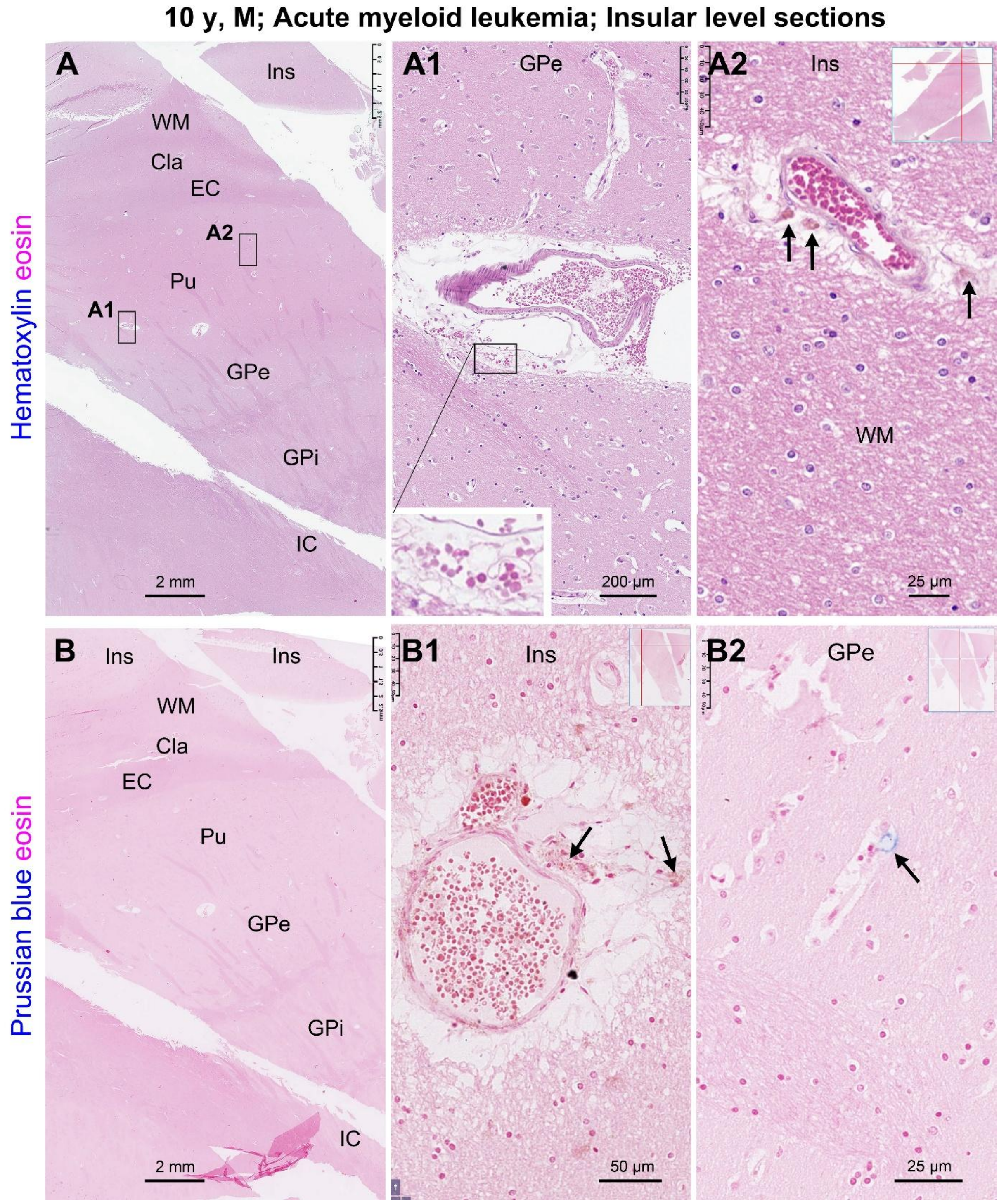


**Supplemental Figure 20. Vascular injury assessed with HE and Prussian stain in the sections passing the striatum in the 10 y case with acute myeloid leukemia.** In the HE stained section, increased perivascular space, reduced local tissue staining intensity (edema) and microbleed (red blood cells and aged reb blood cells stained in brown, pointed by arrows) are evident at the blood vessels in the striatum and internal capsule (**A, A1, A2**). However, there is no apparent increase in the proportion of nucleated cells inside the blood vessel. The above changes are also seen in Prussian stain (**B, B1**), with iron deposition identifiable at a few locations with vascular injury (**B2,** pointed by arrows).

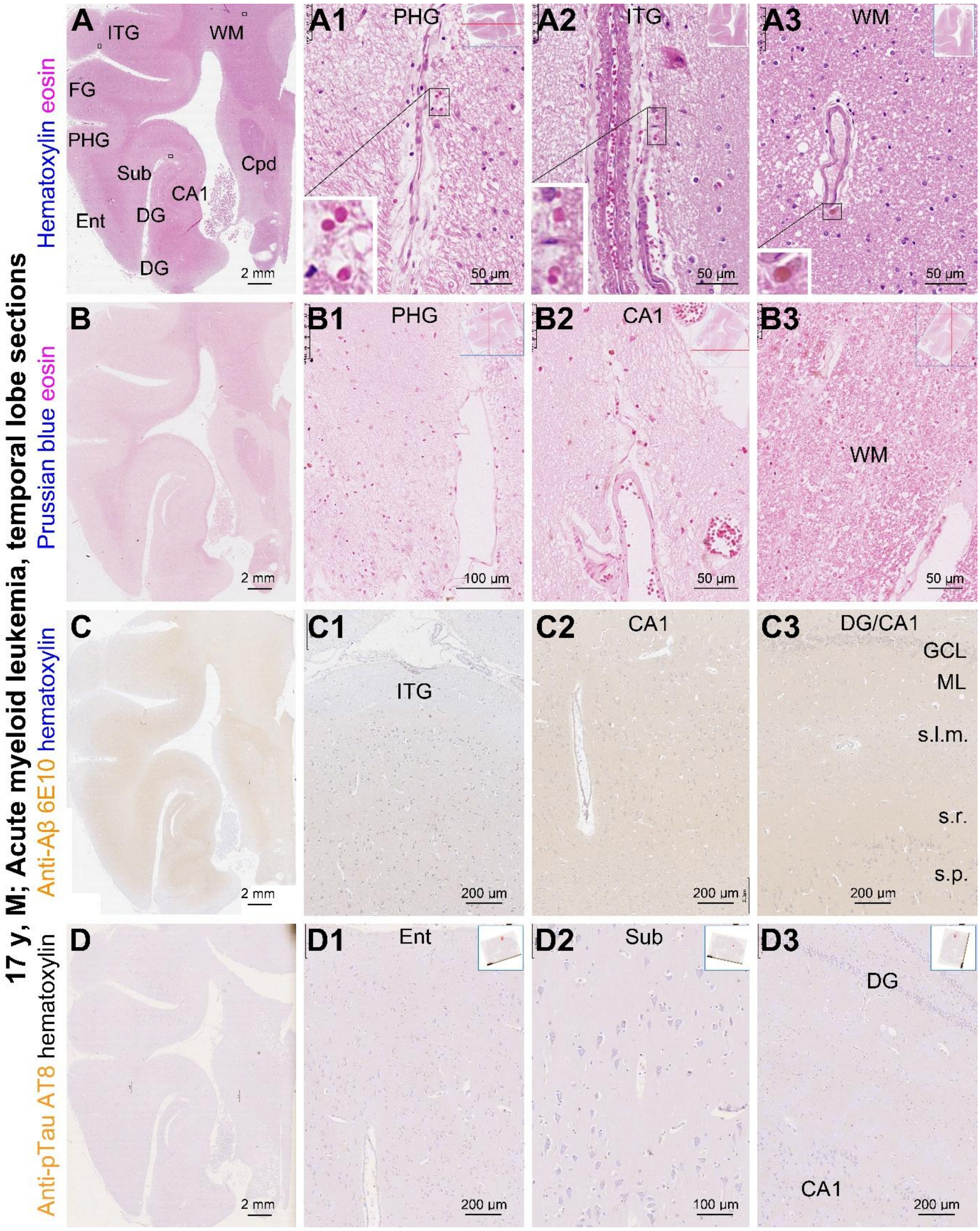


**Supplemental Figure 21. Vascular injury and assessment of AD neuropathologies in the temporal lobe sections from the 17 y with acute myeloid leukemia.** Panels (**A, A1, A2**) show increased perivascular space and microbleed around meningeal vessels in the parahippocampal gyrus (PHG) and interior temporal gyrus (ITG), and in the temporal cortex white matter (WM). The is no apparent iron deposition in the section in Prussian stain (**B1-B3**). No Aβ plaques nor tauopathy are labeled in the temporal cortical areas and the hippocampal subregions in 6E10 and AT8 immunohistochemical preparations (**C, C1-C3, D, D1-D**).

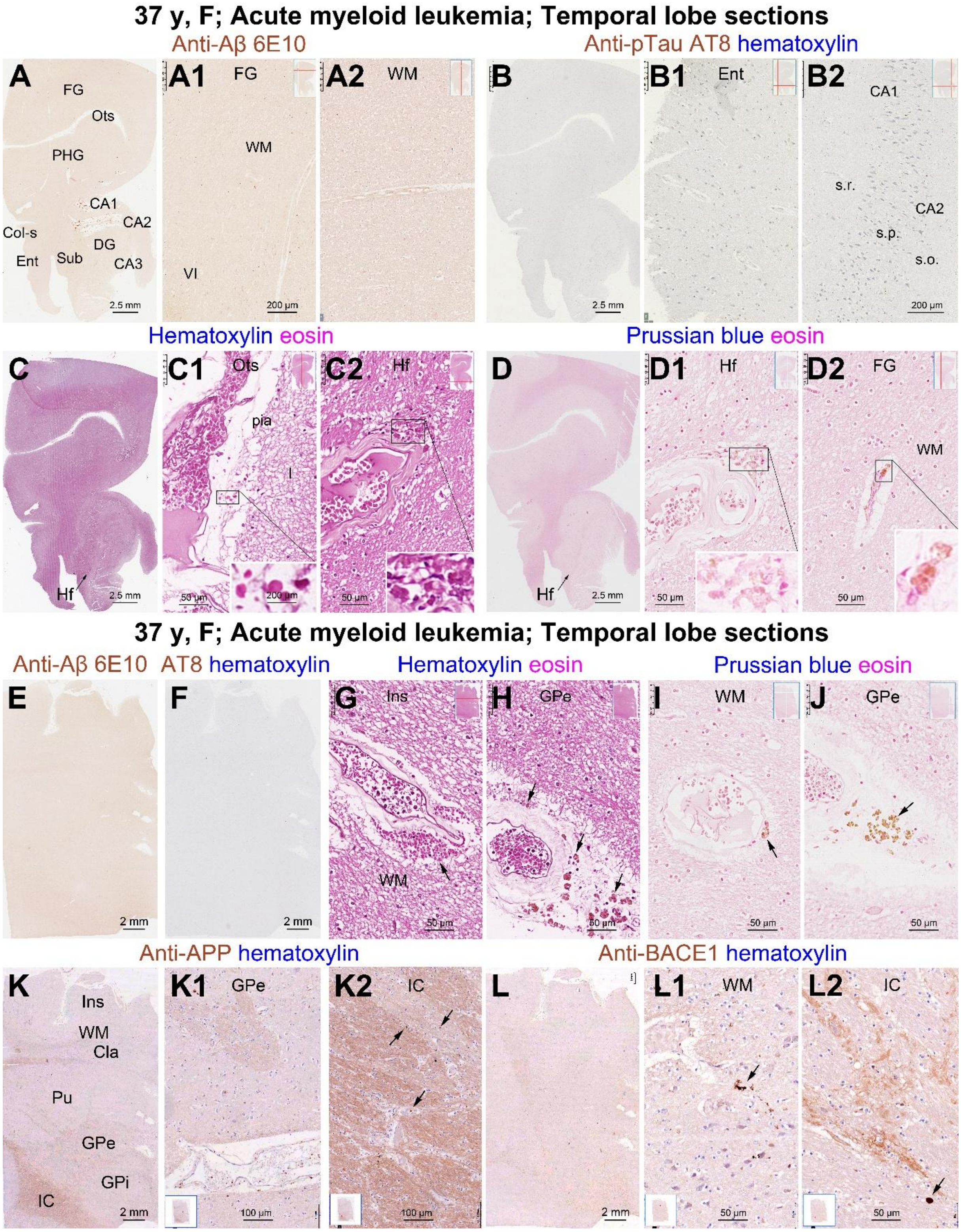


**Supplemental Figure 22. Multi-staining assessment of vascular injury and Alzheimer disease-type neuropathologies in brain regions in the 37 y with AML.** No Aβ (**A, A1, A2**) and tau (**B, B1, B2**) pathologies are present in the temporal lobe structures and across the striatum level section (**E, F**). Increased perivascular spaces and microbleed (inserts and arrows) are found in the regions without overt iron leakage (**C, C1, C2, D, D1, D2, G-J**). Axonal spheroids (arrows) are observed infrequently in the white matter, striatum and internal capsule in APP (**K, K1, K**) and BACE1 (**L, L1, L2**) immunolabeling.

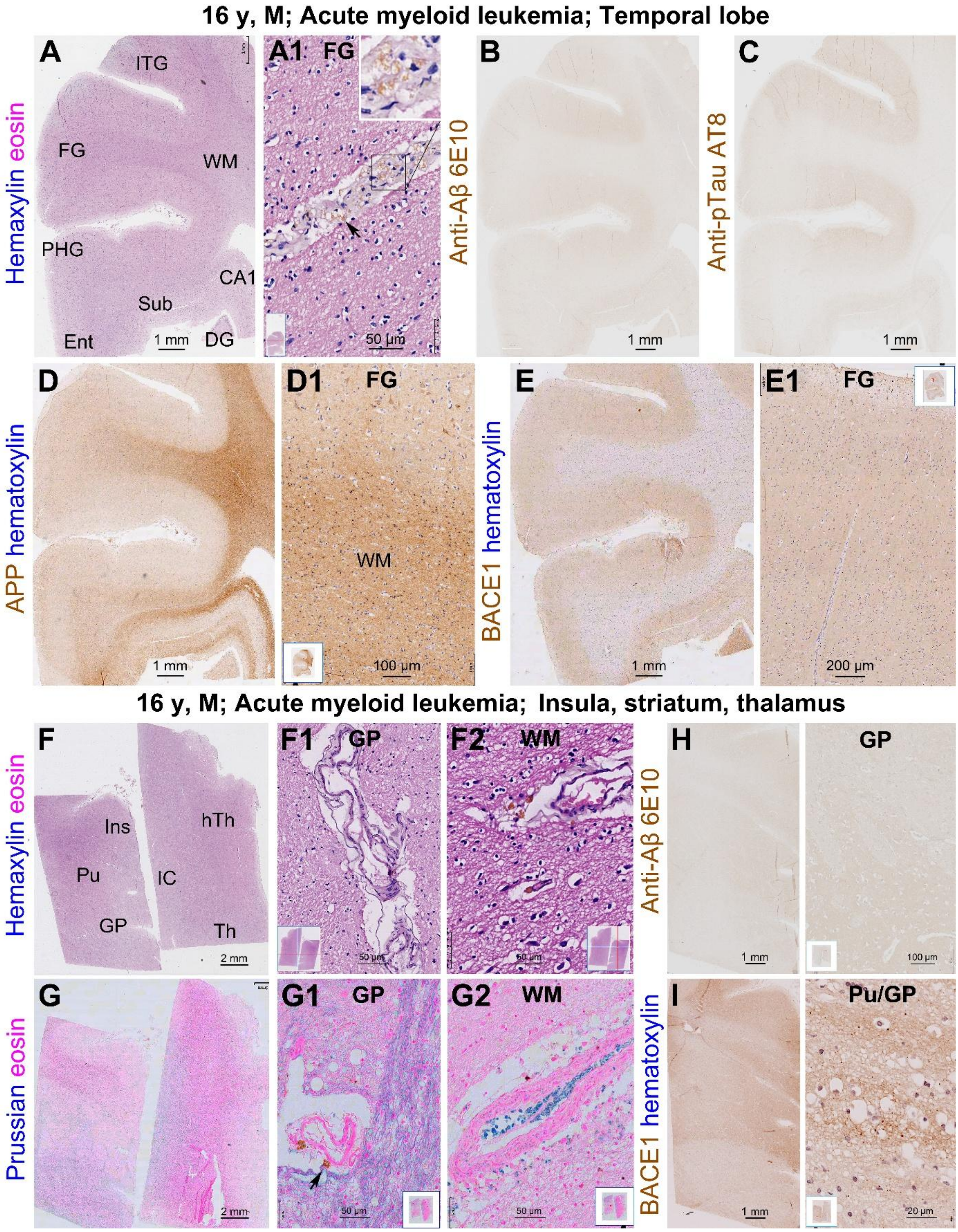


**Supplemental Figure 23. Assessment of vascular injury and AD-type lesions in the brain of the 16 y male with AML.** Vascular wall disruption and microbleed are found in the cortical white matter and striatum and internal capsule (**A, F, G** and enlarged views). Perivascular edema and occurrence of brown stained red blood cells are also found in the cortex (**A1**) and striatum (**F, F1, F2**). Iron leakage is seen in the insular section (G, G1, G2). No Aβ deposition and tauopathy exist in this brain (B, C, H). APP and BACE1 labeling patterns also appear normal (**D, D1, E, E1, I**).

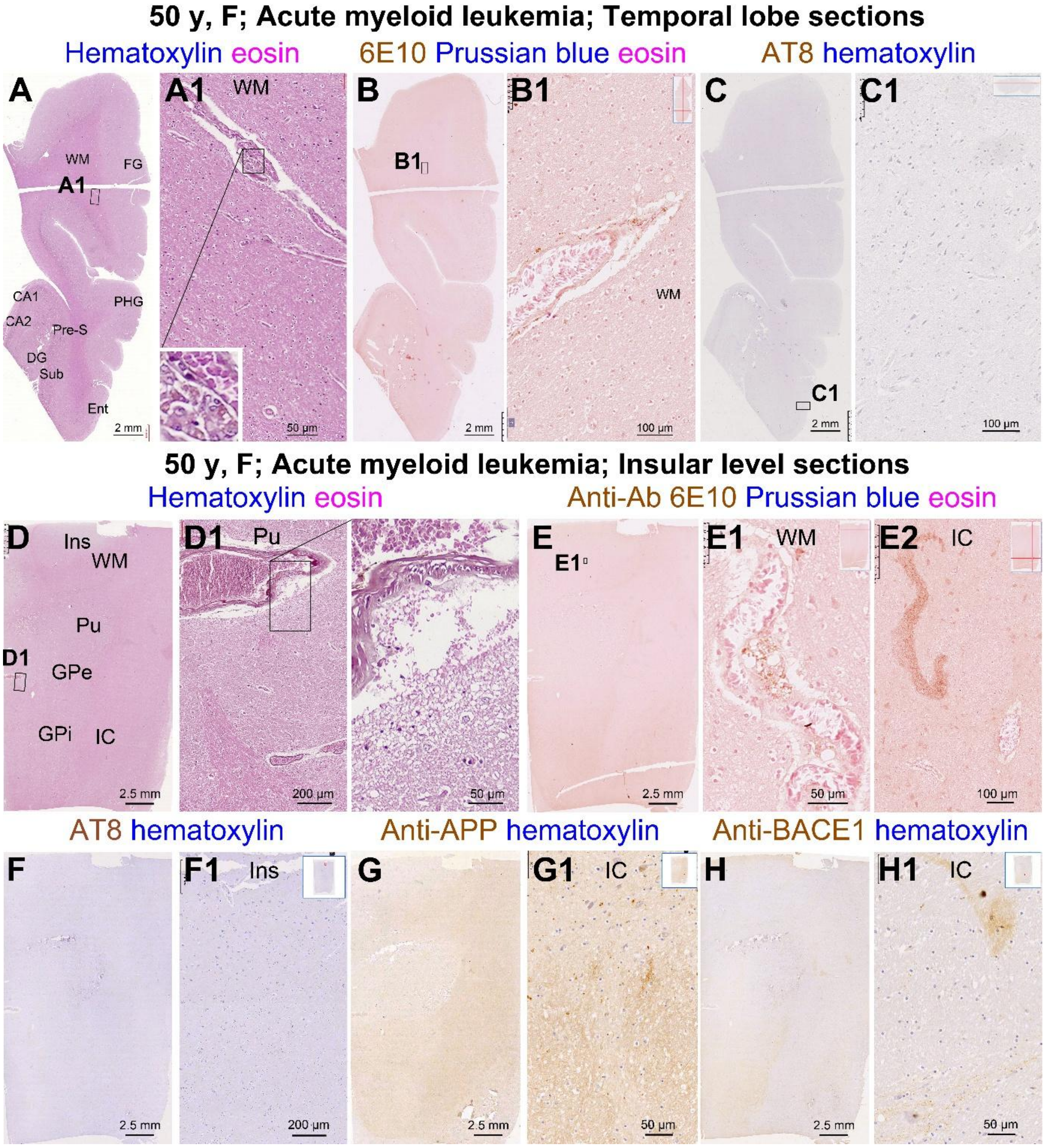


**Supplemental Figure 24. Assessment of vascular injury and AD-type neuropathologies in representative brain regions in the 50 y with acute myeloid leukemia.** Vascular injuries involving microbleed and breakdown of the vascular wall are found around leptomeningeal and intracerebral blood vessels (**A, A1, D, D1**), whereas no apparent iron leakage is detected with Prussian counterstain (**B, B1, E1, E2**). No Aβ plaques and tauopathy are found in the temporal lobe structures (**B, B1, C, C1**) and across the striatum level section (**E, E1, E2, F, F1**). However, axonal pathology is seen locally in the internal capsule in 6E10, APP and BACE1 immunolabeling preparations (**E, E2, G, G1, H, H1**). The proportion of red blood cells relative to nucleated cells inside the blood vessels appears to be in normal range in this brain as seen in HE preparation (**A1, D1**).

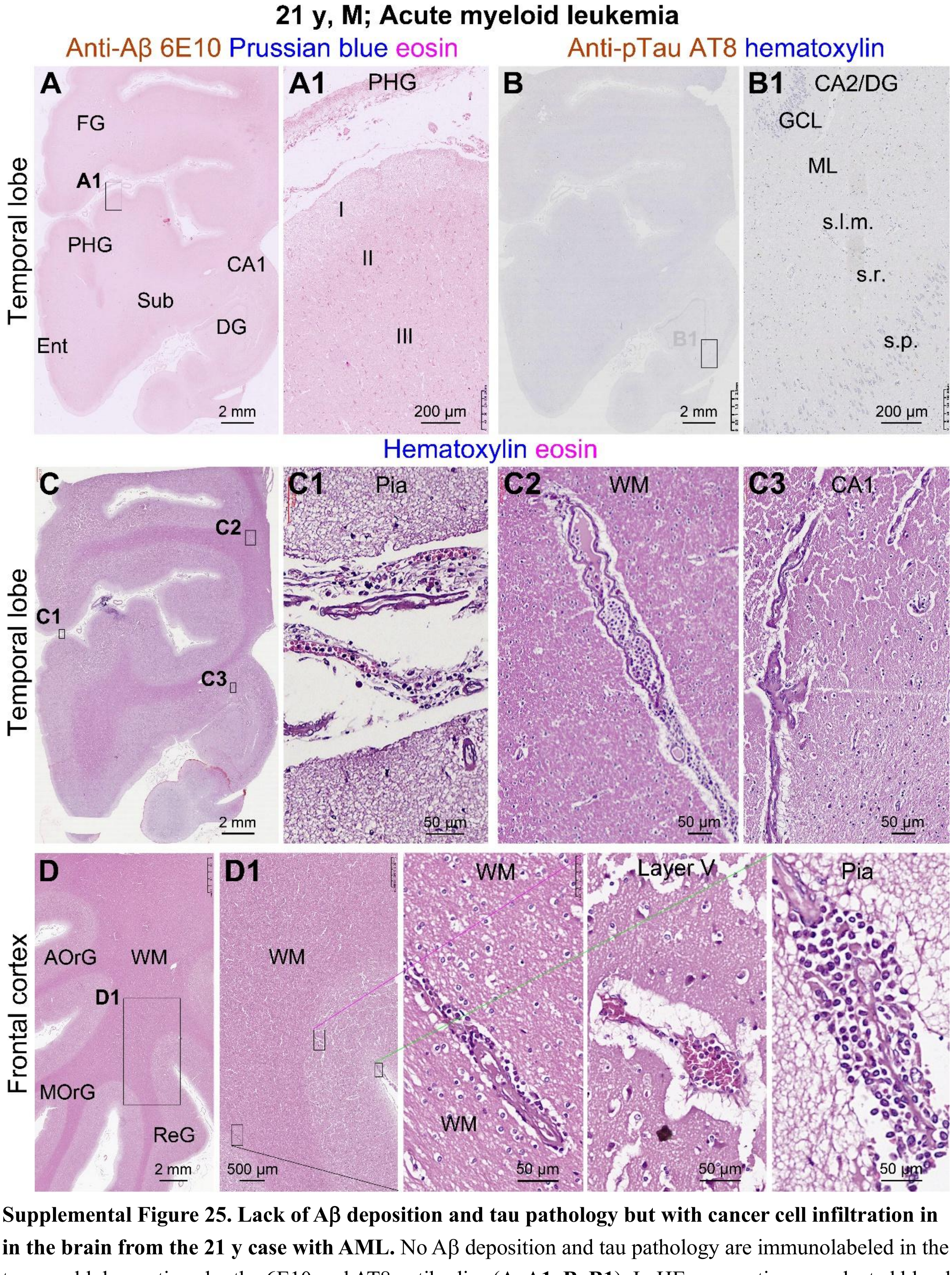


**Supplemental Figure 25. Lack of Aβ deposition and tau pathology but with cancer cell infiltration in in the brain from the 21 y case with AML.** No Aβ deposition and tau pathology are immunolabeled in the temporal lobe sections by the 6E10 and AT8 antibodies (**A, A1, B, B1**). In HE preparations, nucleated blood cells are apparently increased in proportion inside blood vessels are also leaked out of the vessels, which are seen around the leptomeninges as well as in grey and white matter areas (**C, C1-C3**; **D, D1** and enlarged panels). Increase perivascular space is seen in intracerebral blood vessels (C2, **C3, D1**).

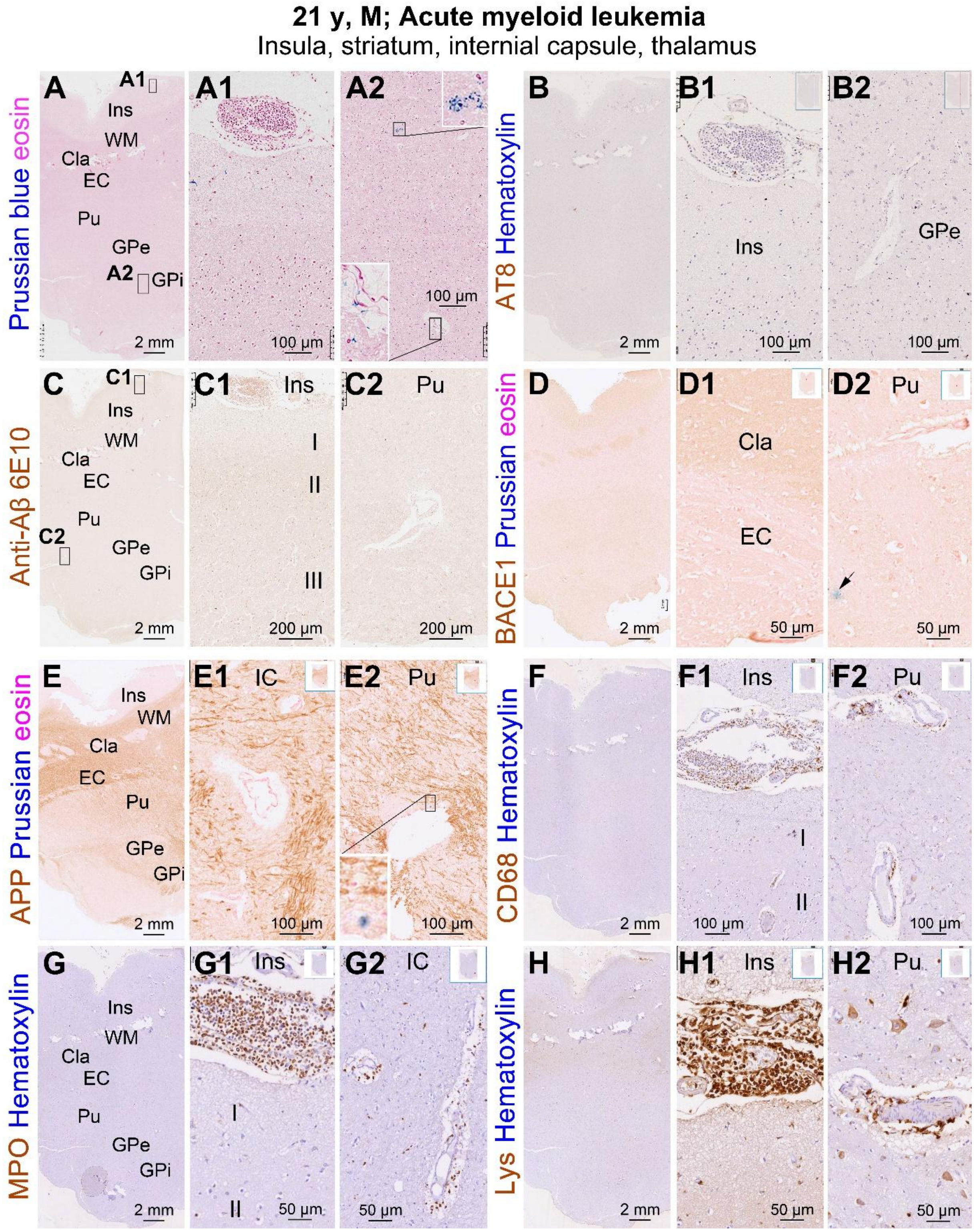


**Supplemental Figure 26. Cancer cell infiltration, vascular injury and axonal pathology in the striatum level sections in the 21 y case with AML.** Iron deposition is detectable around some blood vessels (**A, A2, D, D2, E, E2**). No Aβ deposition and tauopathy are present (**B, B1, B2, C, C1, C2**). Increased axonal APP labeling is seen around damaged blood vessels (**E, E1, E2**). The nucleated blood cells express CD68 (**F, F1, F2**) and MPO (**G, G1, G2**) partially. Lysozyme labeling is evident in these blood cells (**H, H1**) and also seen in some large-sized neuronal somata around broken vessels (**H2**).

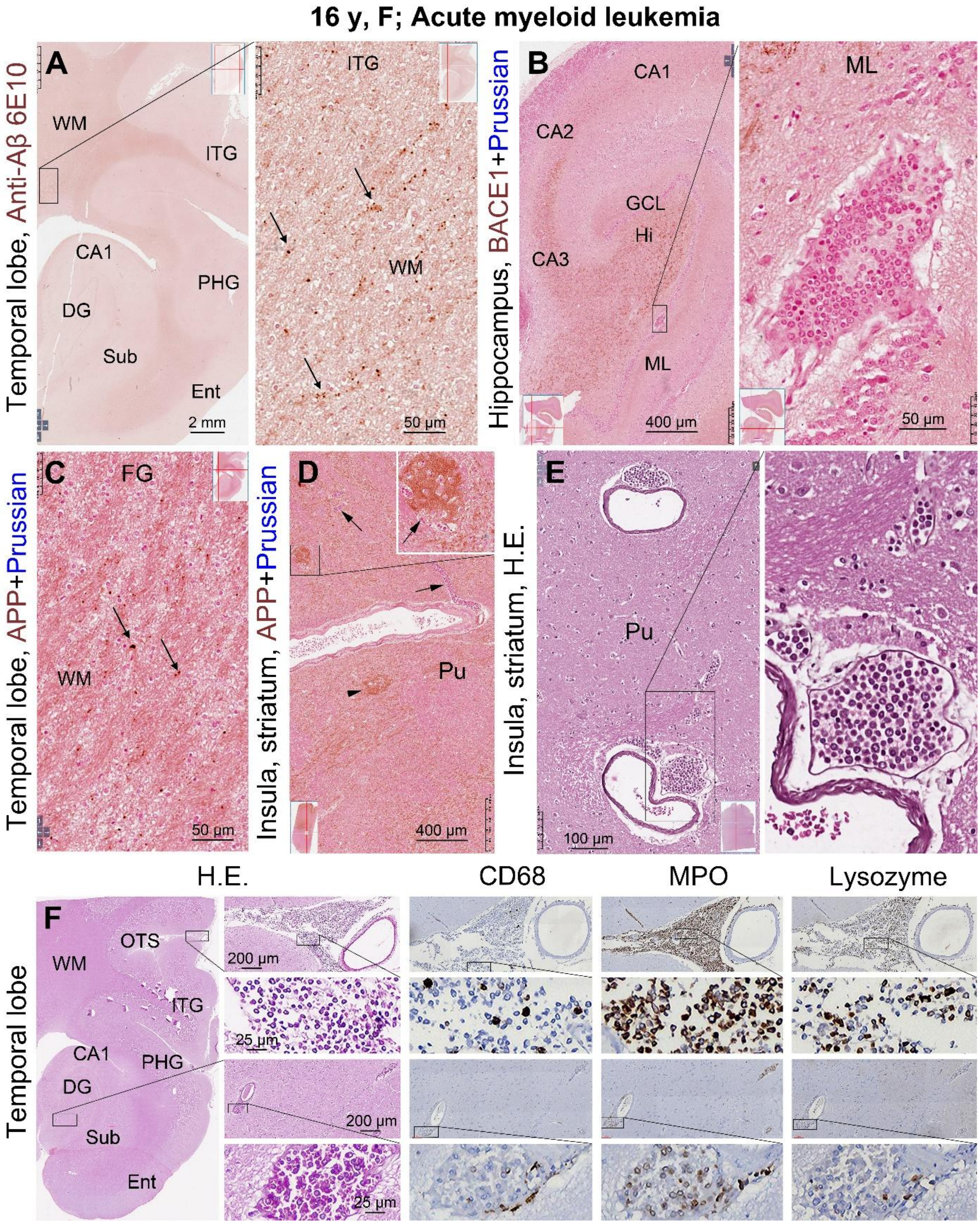


**Supplemental Figure 27. Cancer cell infiltration, vascular injury and axonal pathology in the temporal lobe sections in the 16 y female with AML.** Swollen axonal processes in cortical white matter and in the striatum (**D**) are labeled by the 6E10 (**A**) and APP (**C, D**) antibodies, and they are densely packed around vasculature in the striatum (**D**). Iron deposition is not microscopically evident in the sections (**B, D**). Nucleated cancer cells are present inside meningeal and intracerebral blood vessels and also form perivascular islands (**B, E, F**), and they partially express the myelogenous cell markers CD68, MPO and lysozyme (**F**).

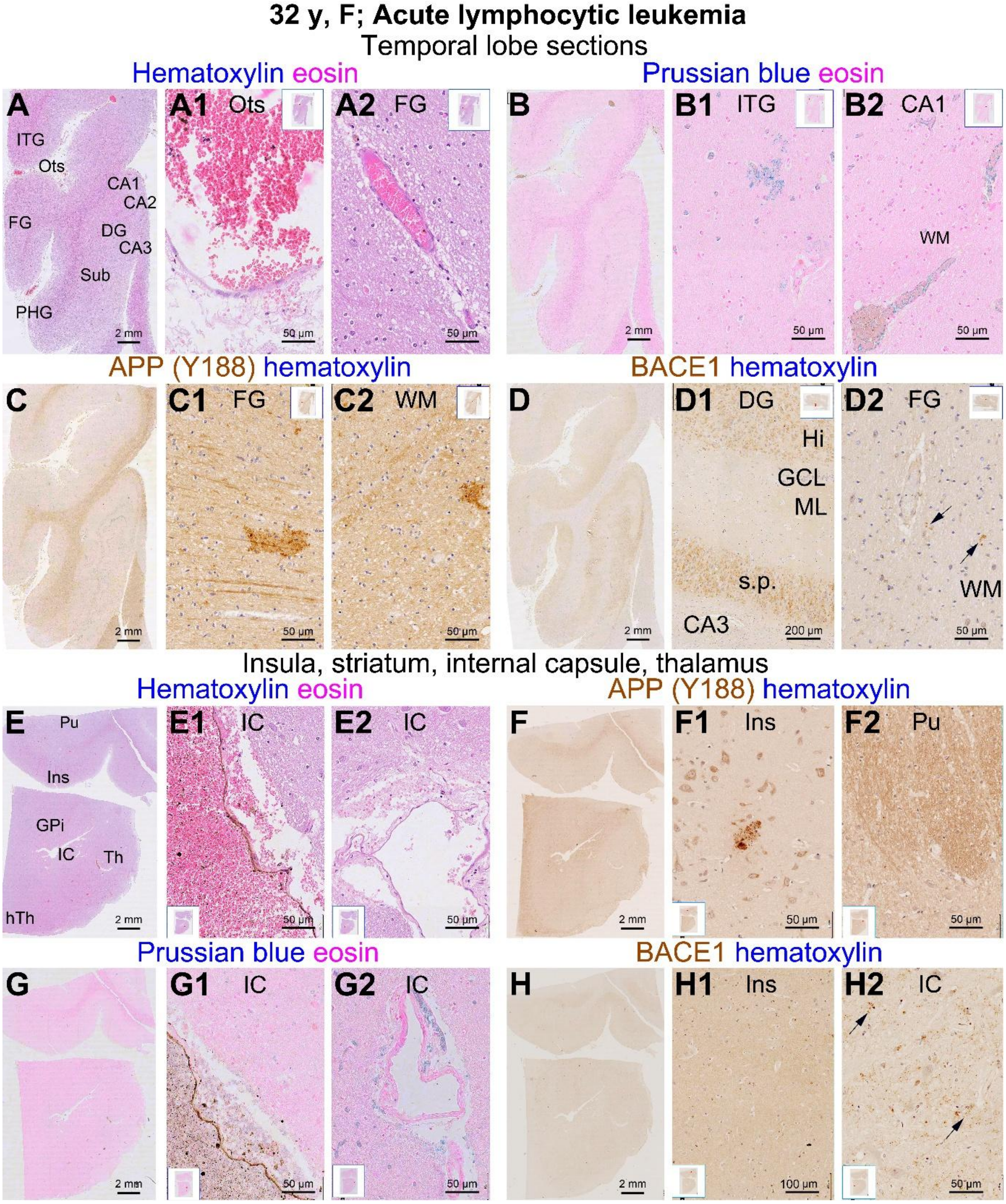


**Supplemental Figure 28. Vascular injury and axonal pathology in the brain of the 32 y donor with acute lymphocytic leukemia.** Extensive disruption of the wall of blood vessels, leakage of blood cells and deposition of iron are observed in multiple cerebral regions and subcortical areas; shown here are representative micrographs of the temporal lobe and insular level sections as indicated. In APP and BACE1 immunolabeling, swollen axonal processes or clusters (pointed by arrows) are observed infrequently in the temporal lobe cortex and white matter and in the internal capsule. The cells inside the blood vessels are predominately the red blood cells rather than nucleated cells.

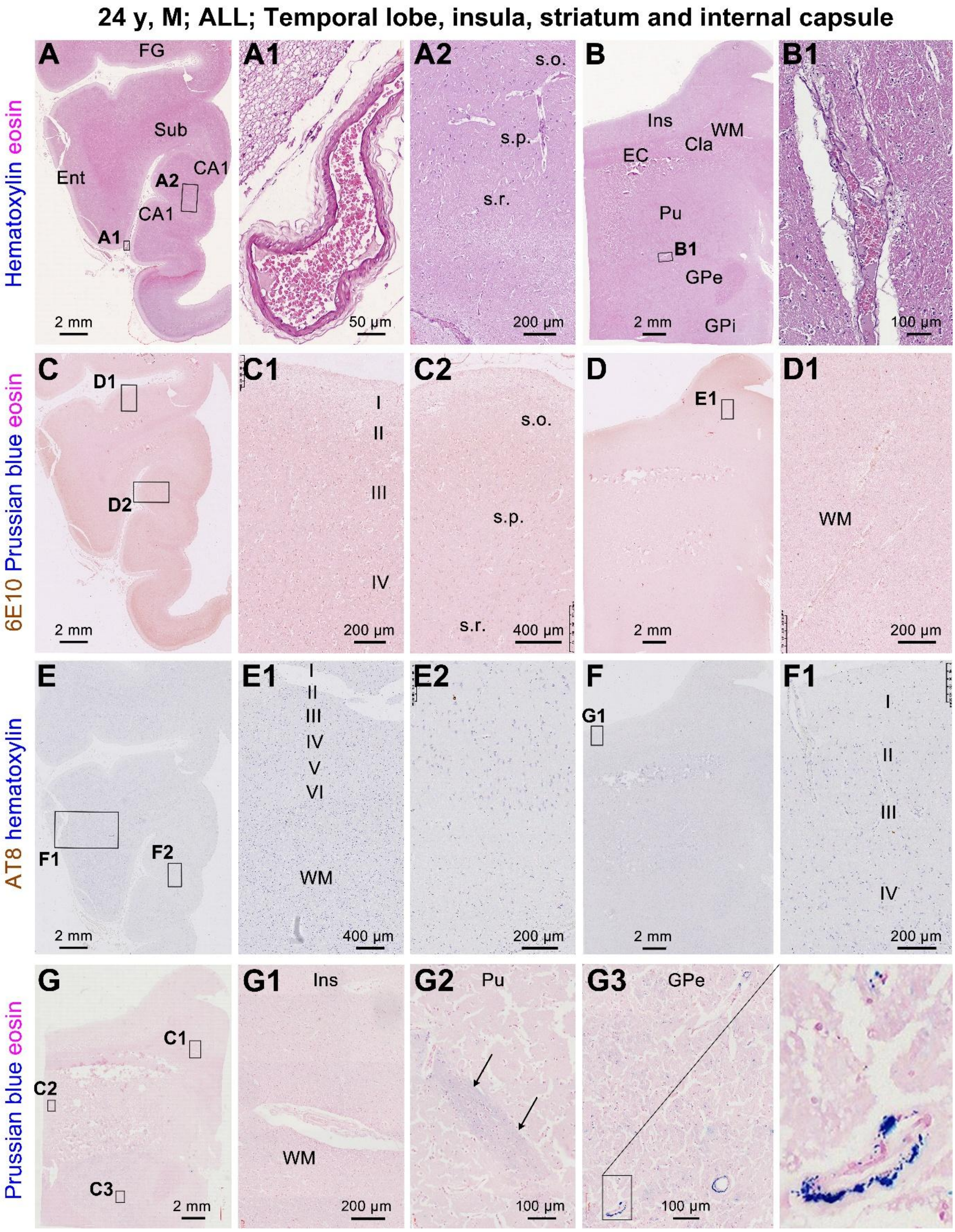


**Supplemental Figure 29. Vascular injury and the lack of neuropathological changes in the 24 y case of acute lymphocytic leukemia.** Increased perivascular space is seen in the temporal lobe and striatum level sections in HE preparation (**A, A1, B, B1**). Red blood cells are predominant inside vasculature (**A1, B1**). No Aβ deposition or tau pathology are labeled by the 6E10 and AT8 antibodies in the sections (**C, C1, C2, D, D1, E, E1, E2, F, F1**). At the striatum level, iron leakage is present around small blood vessels and in the axonal bundles (**G, G1, G2, G3**).

48 y, M; Myelodysplastic syndrome (MDS-EB-2)

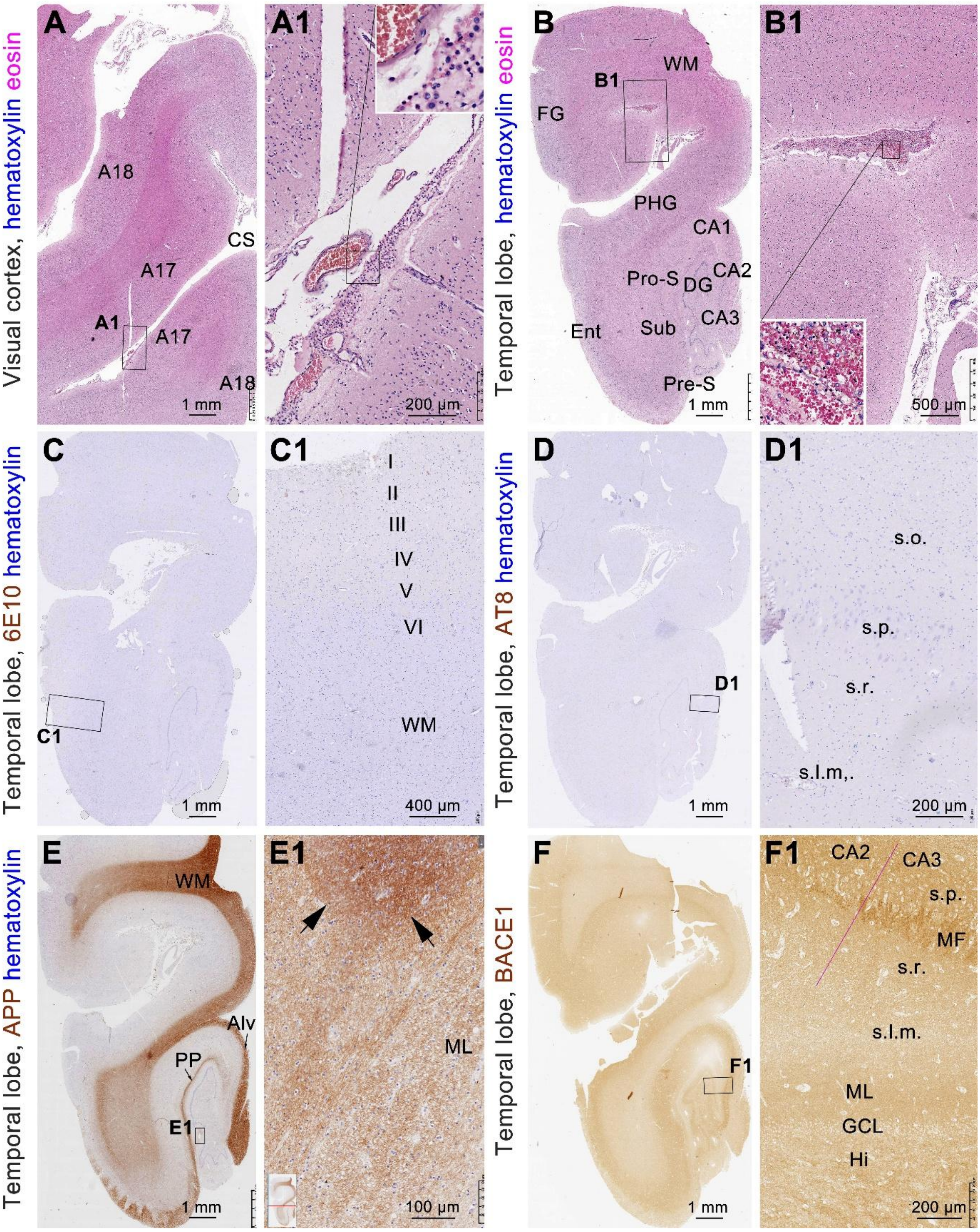


**Supplemental Figure 30. Infiltration of tumorous cells, vascular injury and white matter damage in cerebral cortex and hippocampal formation in the 48 y case of myelodysplastic syndrome.** Increased nucleated cells in blood vessels and leakage of red and white blood cells is evident around the leptomeninges in HE preparation (**A, A1, B, B1**). No Aβ deposition and tau pathology are found (**C, C1, D, D1**). The APP labeling appears enhanced locally (arrows) in the white matter, although no apparent axonal swelling (**E, E1**). BACE1 labeling exhibits a normal-looking appearance in the temporal lobe (**F, F1**).

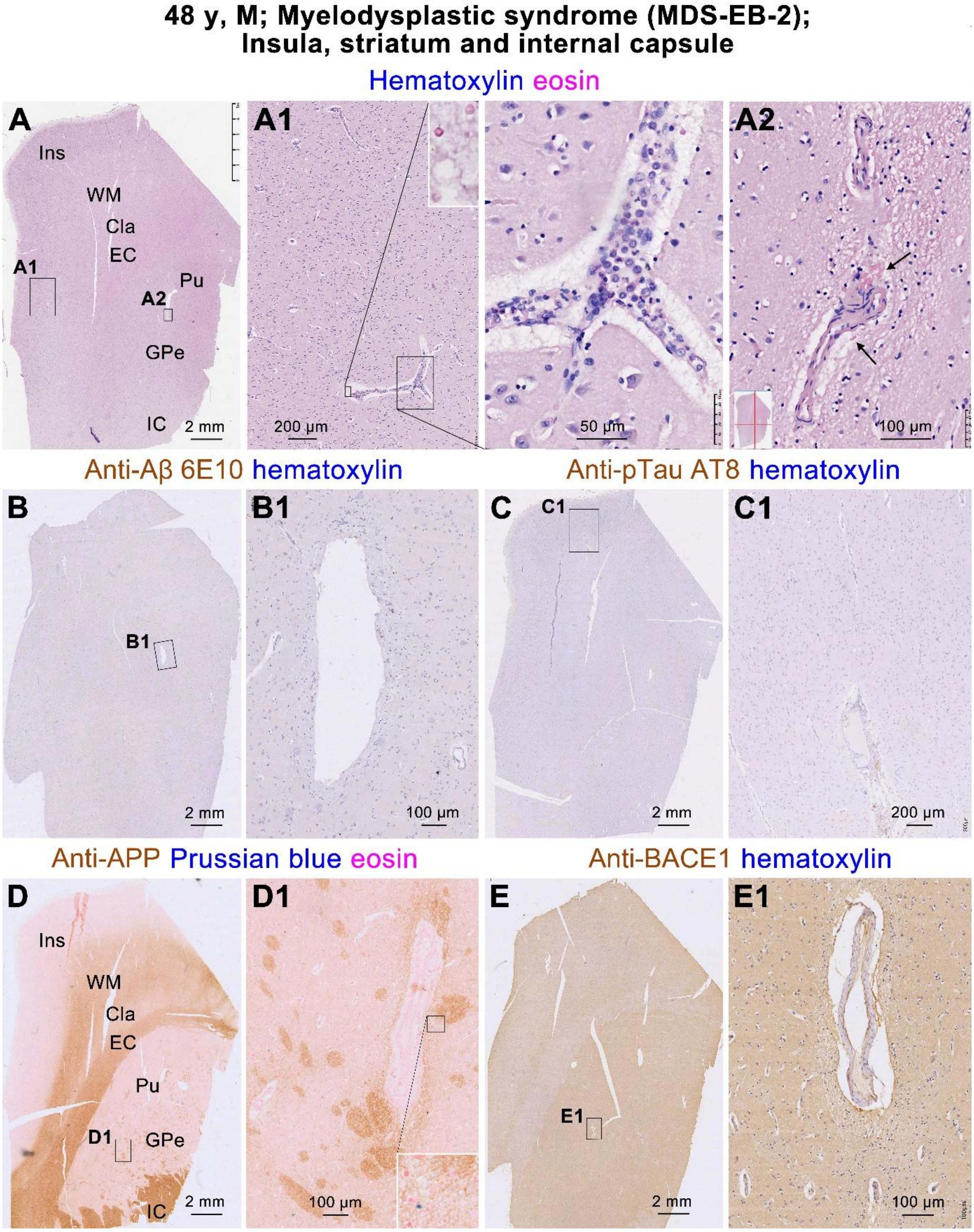


**Supplemental Figure 31. Infiltration of tumorous cells, vascular injury and white matter damage in the striatum level sections in the 48 y case of myelodysplastic syndrome.** Increased nucleated cells in blood vessels, microbleed and increase perivascular space are present in HE stain (**A, A1**). No Aβ deposition and tau pathology are present (**B, B1, C, C1**). The APP labeling appears enhanced around striatal blood vessels with minor local iron deposition (**D, D1**). BACE1 labeling exhibits a normal-looking neuropil pattern in the striatum, with a small amount of strongly labeled element located near the damaged blood vessels (**E, E1**).

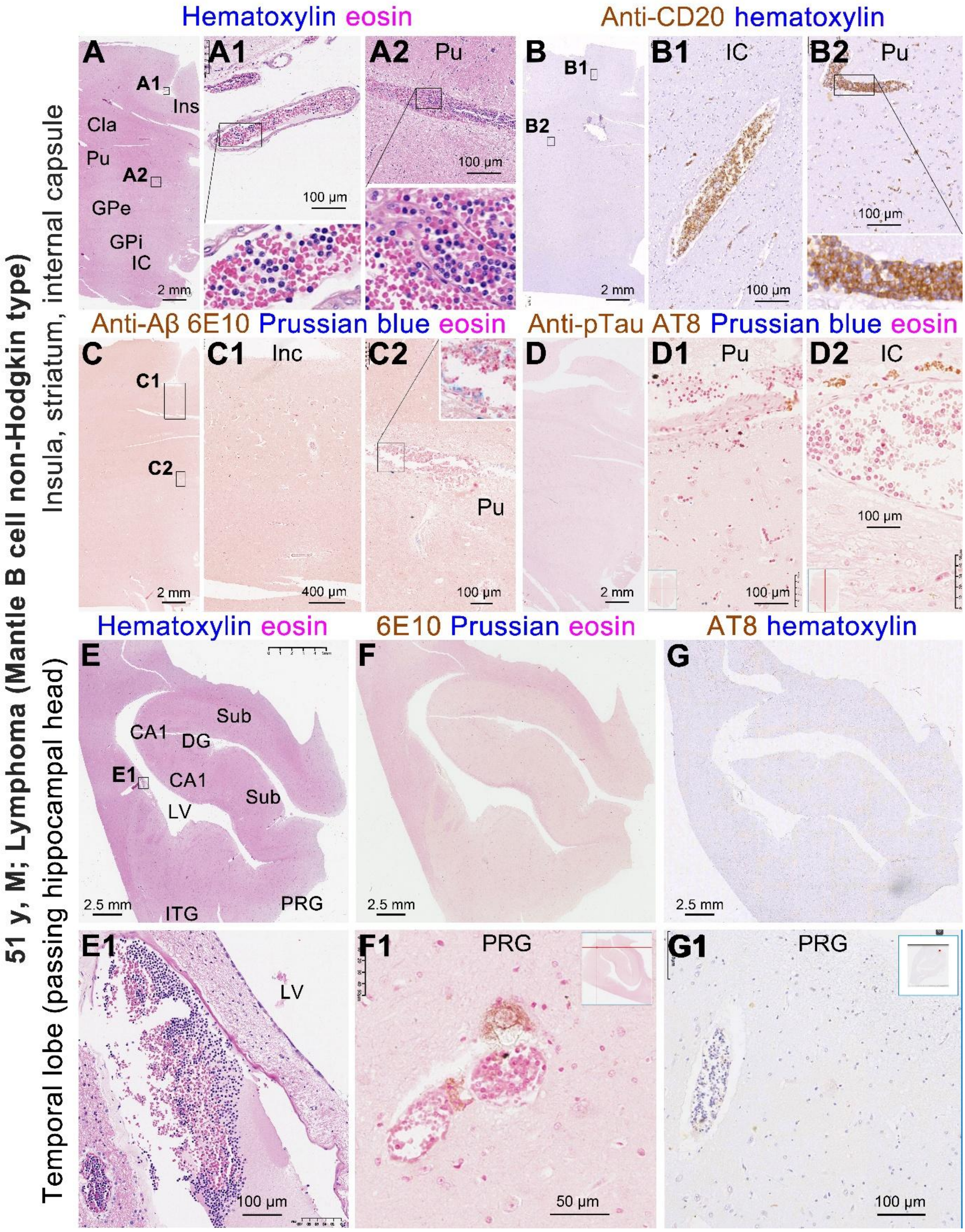


**Supplemental Figure 32. Pathological examination in representative sections in the 51 y case of mantle cell lymphoma.** Increased nucleated cells in blood vessels, microbleed including aged blood cells in brown stain, increase perivascular space, and iron deposition with mild extent, are detected in the temporal lobe and striatum level sections in HE and Prussian stains. No Aβ deposition and tau pathology are found in the sections. The tumorous blood cells are immunoreactive for CD20, which is a molecular marker of mantle cell lymphocytes.

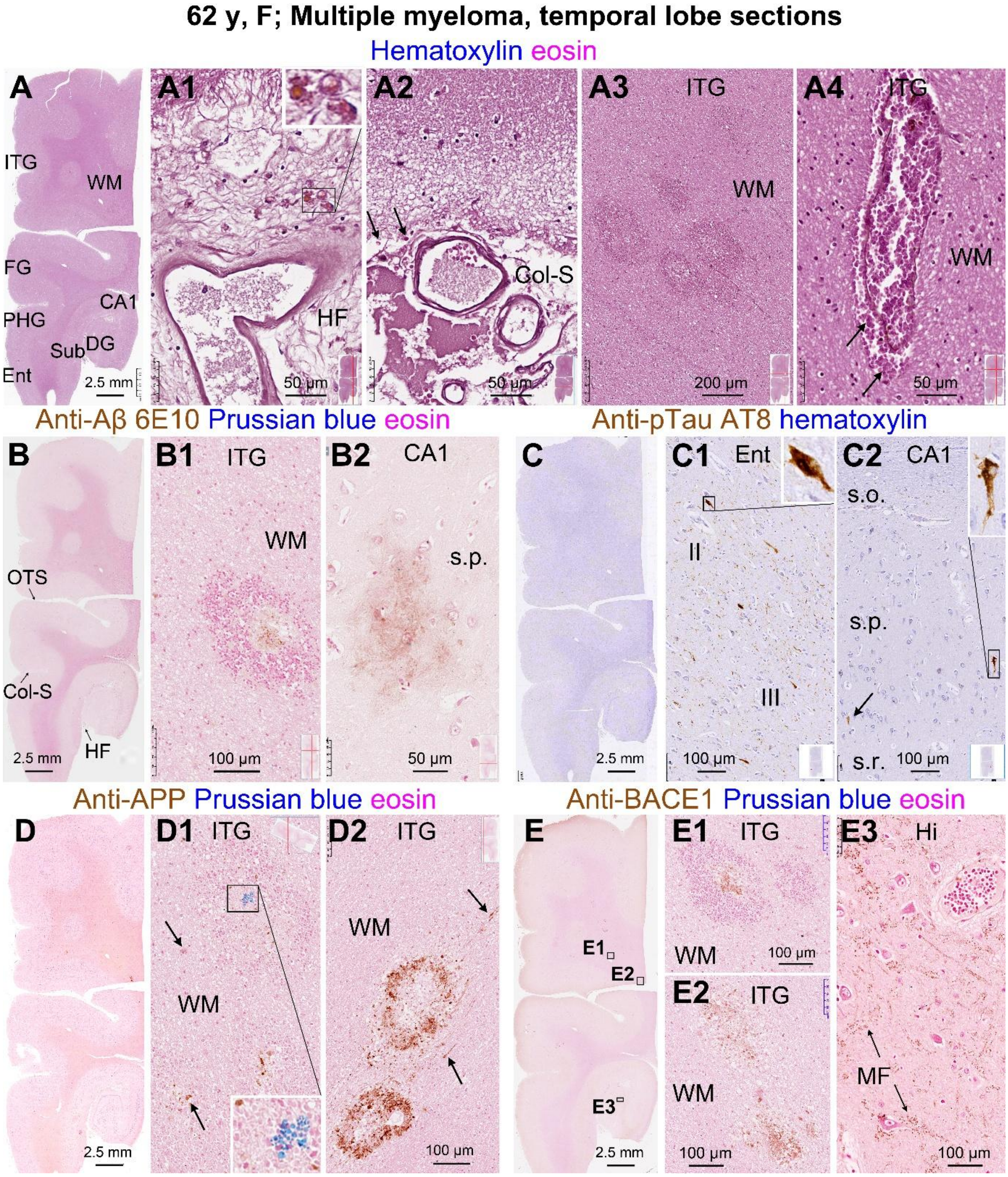


**Supplemental Figure 33. Vascular injury, white matter damage and axonal pathology in temporal lobe sections in the 62 y case of multiple myeloma.** Microbleed including leakage of aged red blood cells (stained brown) are present around blood vessels at the leptomeningeal (**A, A1, A2**) and in the white matter (**A3, A4**). No extracellular Aβ plaques are labeled by the 6E10 antibody except for isolated patch-like labeling likely related to axonal pathology (**B, B2**). However, increased APP and BACE1 immunolabeling representing amyloidogenic axonal pathology are concurrently present at the injured white matter locations (**B, B1, D, D1, D2, E, E1**). Pretangle neurons and neuropil threads are labeled by the AT8 antibody, with the tau pathology scored at Braak stage II. The mossy fiber BACE1 labeling in the dentate gyrus exhibits a normal-looking pattern (**E3**).

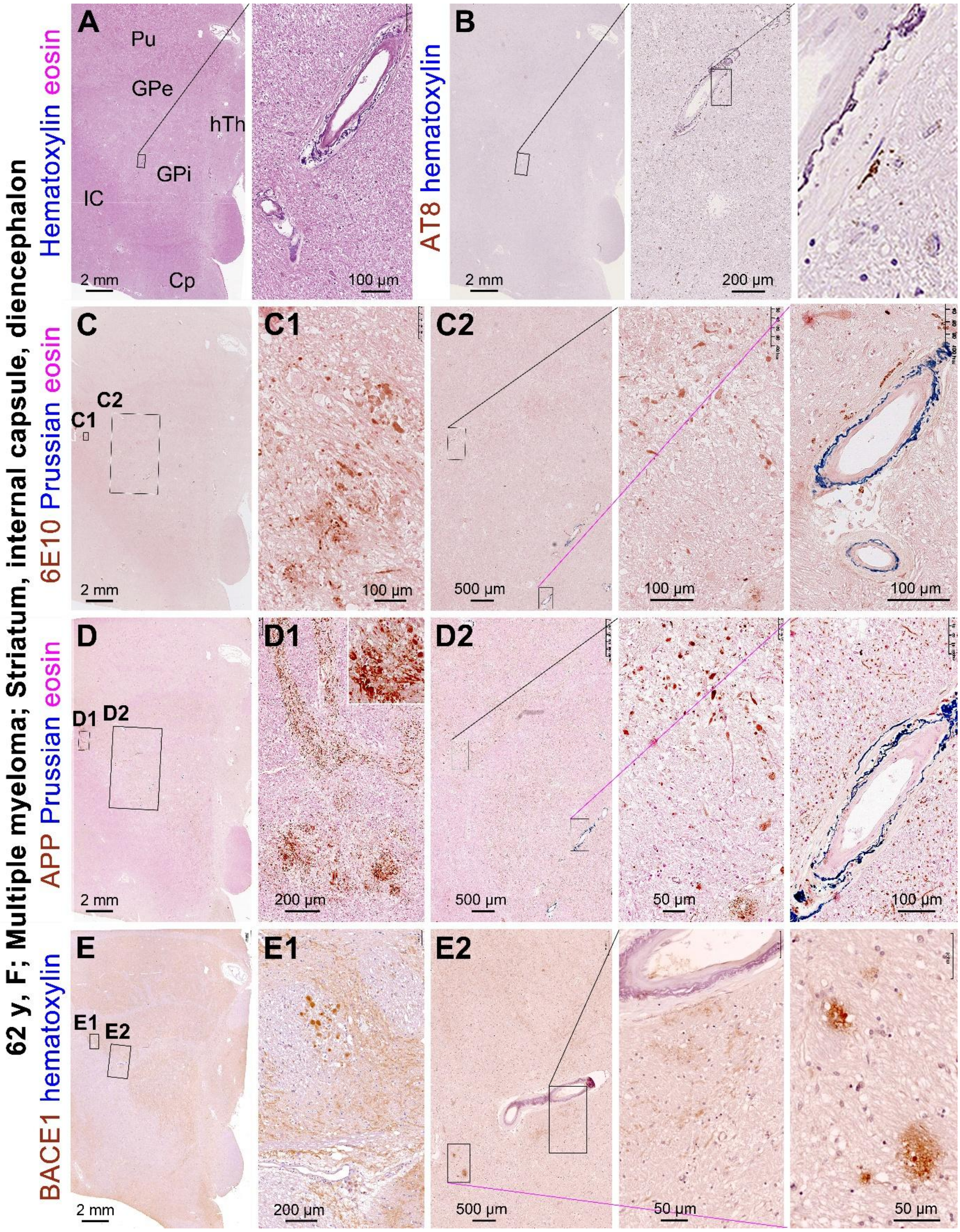


**Supplemental Figure 34. Vascular injury and axonal pathology in the striatum in the 62 y case of multiple myeloma.** Increased perivascular space, leakage of blood cells, iron deposition, hyalinization and calcification (**A, D2, E2**) of the vascular wall are shown. Swollen axons and clusters are labeled by 6E10, APP and BACE1 antibodies, some occurring around the damaged vessels (**C, D, E** and enlarged panels). In the AT8 immunolabeled section, a few labeled astrocytes are observed near the damaged vessels, whereas no neuronal somata or neuropil threads are labeled across the section (**B** and enlarged views).

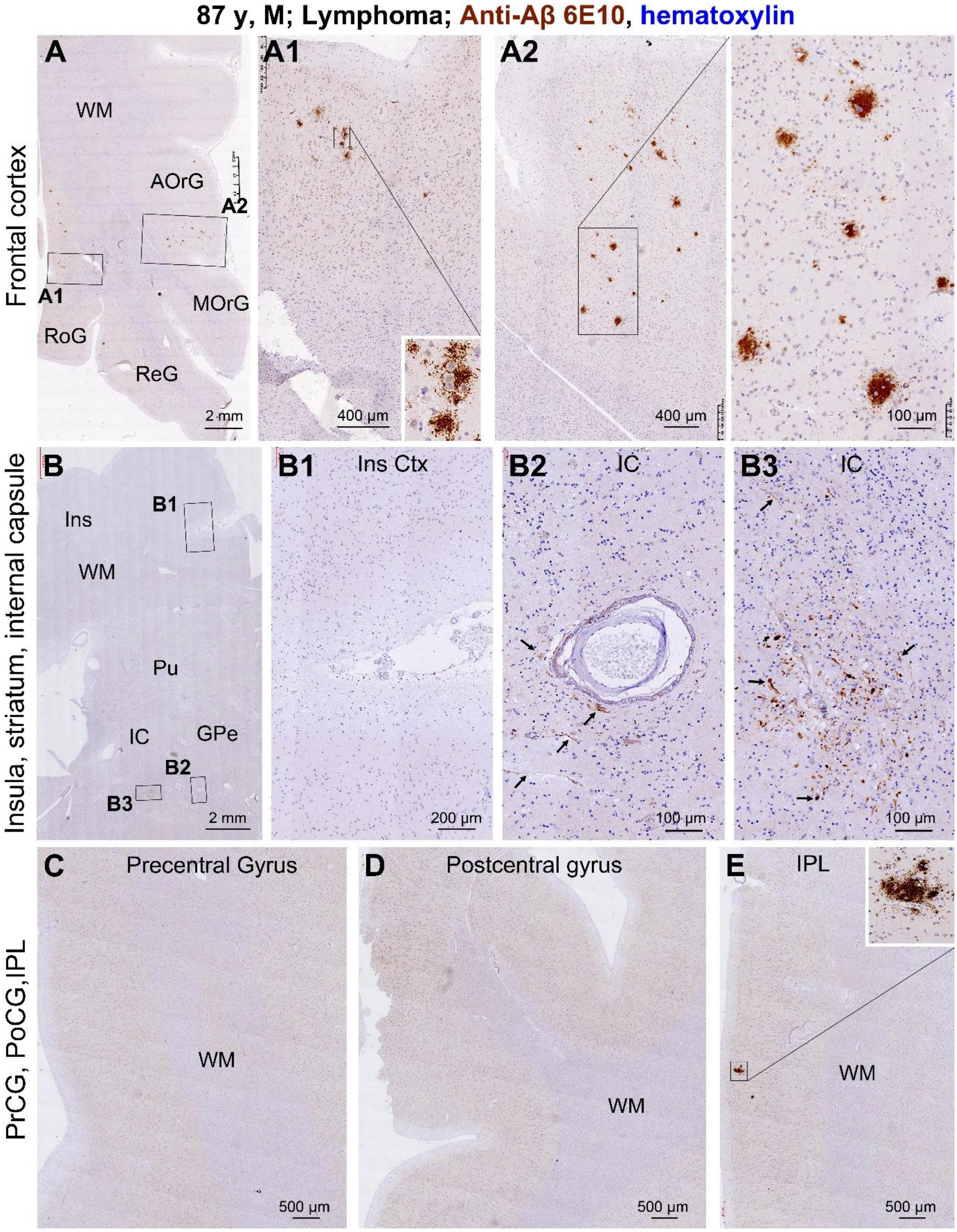


**Supplemental Figure 35. 6E10 antibody labeling in multiple brain regions in the 87 y lymphoma case.** Extracellular Aβ deposition occurs in subareas of the frontal cortex appearing as diffuse plaques (**A, A1, A2**), but is rarely seen in the insular (**B, B1**), primary motor and sensory (**C, D),** or the inferior parietal (**E)** cortices. However, swollen axonal processes and clusters are labeled by this antibody in the striatum and internal capsule around blood vessels (**B2**) and at the sites with vascular injury (**B3**).

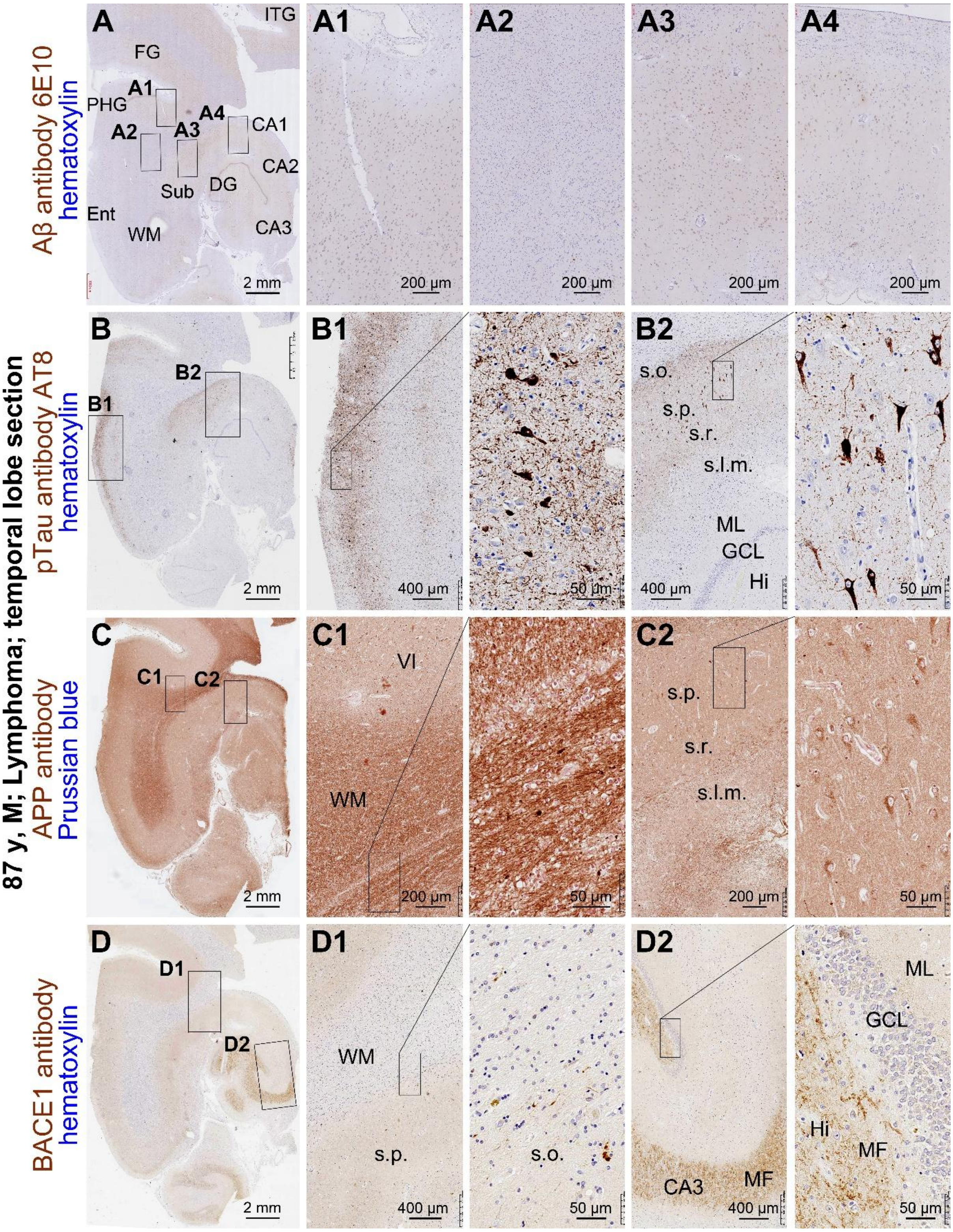


**Supplemental Figure 36. Assessment of Aβ, tau and axonal pathology in temporal lobe sections in the 87 y lymphoma case.** No Aβ plaques are labeled by the 6E10 antibody in the temporal cortex and hippocampal formation (**A, A1-4**). Tau pathology is labeled by the AT8 antibody largely in the entorhinal cortex and the hippocampus (**B, B1, B2**), with the lesions in this brain scored as Braak stage IV. Small amounts of swollen axonal processes are labeled by APP (**C, C1**) and BACE1 (**D, D1**) antibodies in the white matter, with normal looking neuronal somata in the cortex (**C2**) and mossy fiber terminals (**D2**).

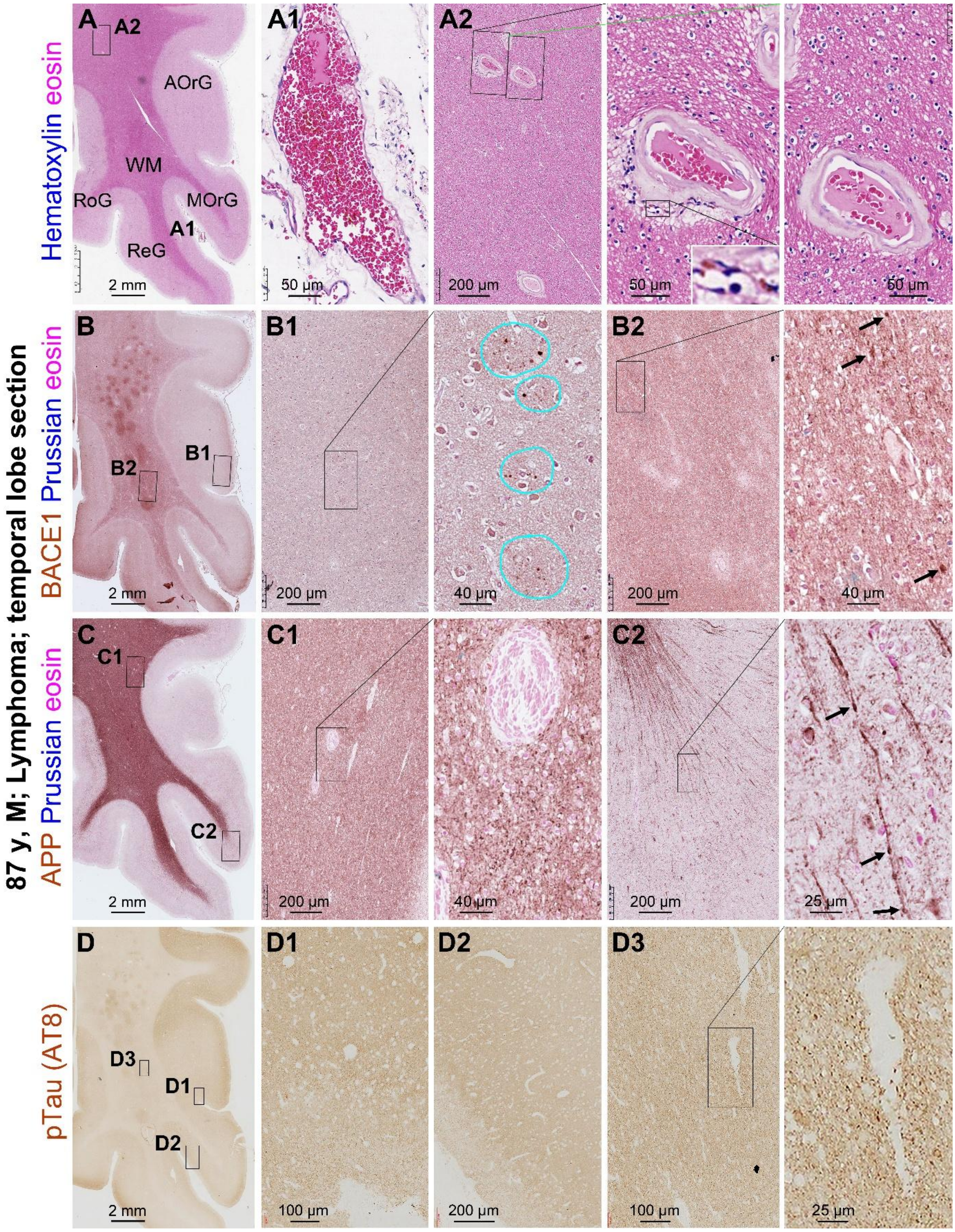


**Supplemental Figure 37. Pathological assessment in frontal cortical sections in the 87 y lymphoma case.** Red blood cells are predominant in blood vessels, increased perivascular space and microbleed at some blood vessels in the white matter (**A, A1, A2**). Uneven BACE1 and APP labeling are seen in white matter, with some swollen axons (pointed by arrows) in the cortex and white matter can be observed by closer examination (B, B1, B2; C, C1, C2). No neuronal somata or neuropil threads are labeled by the AT8 antibody in the cortex or white matter (**D, D1-D3**).

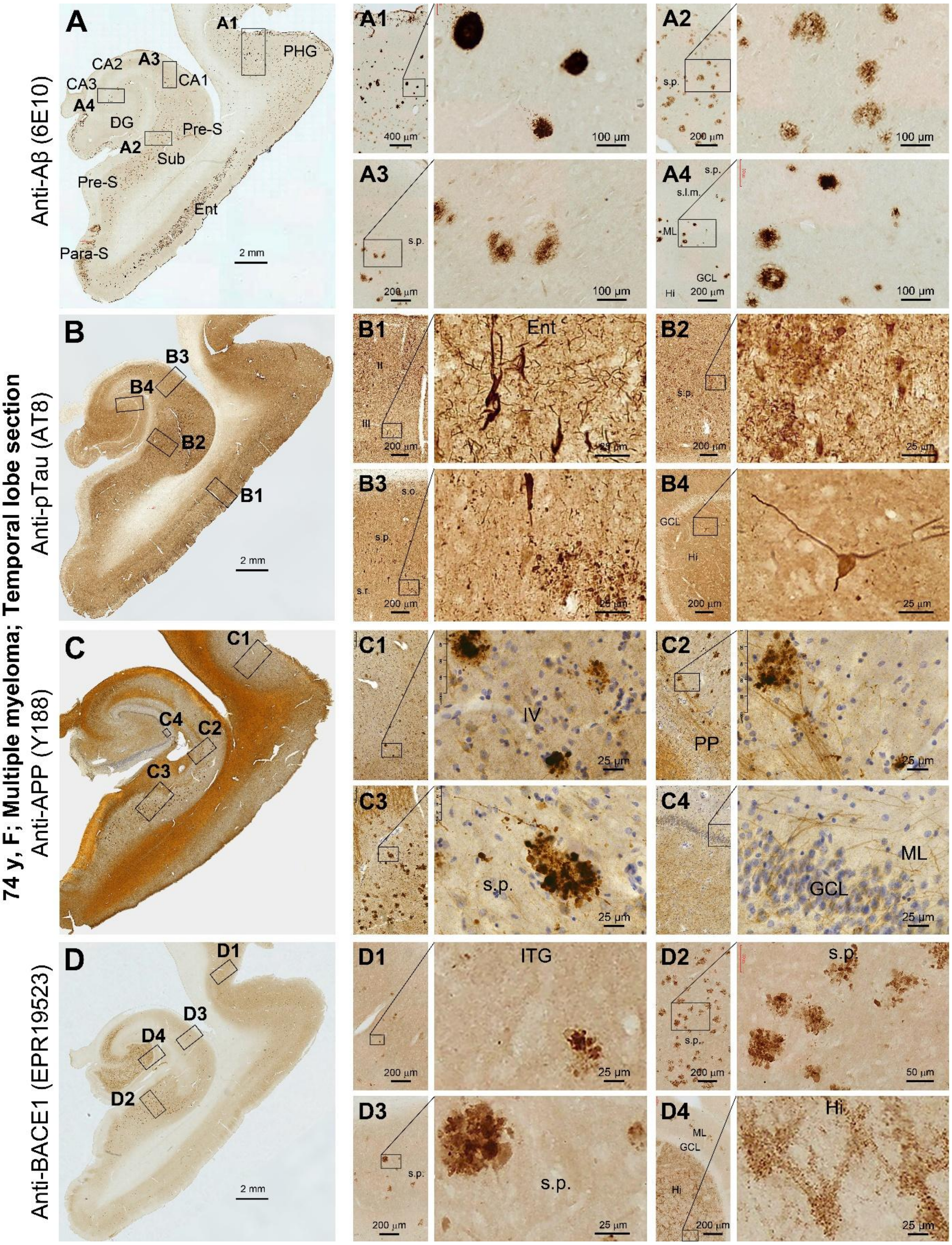


**Supplemental Figure 38. AD-type neuropathology in temporal lobe sections in the 74 y case with multiple myeloma.** Extensive Aβ pathology (Thal phase 4) is labeled by the 6E10 antibody, appearing as compact and diffuse plaques as well as CAA (**A, A1-A4**). Tangle-like neurons, neuritic clusters and neuropil threads are labeled by the AT8 antibody (**B, B1-C4**), with the whole brain tauopathy scored as Braak stage IV. Dystrophic neuritic clusters are distinctly labeled by the APP (**C, C1-C4**) and BACE1 (**D, D1-D3**) antibodies, with a normal looking BACE1 labeling of the mossy fiber terminal (**D4**).

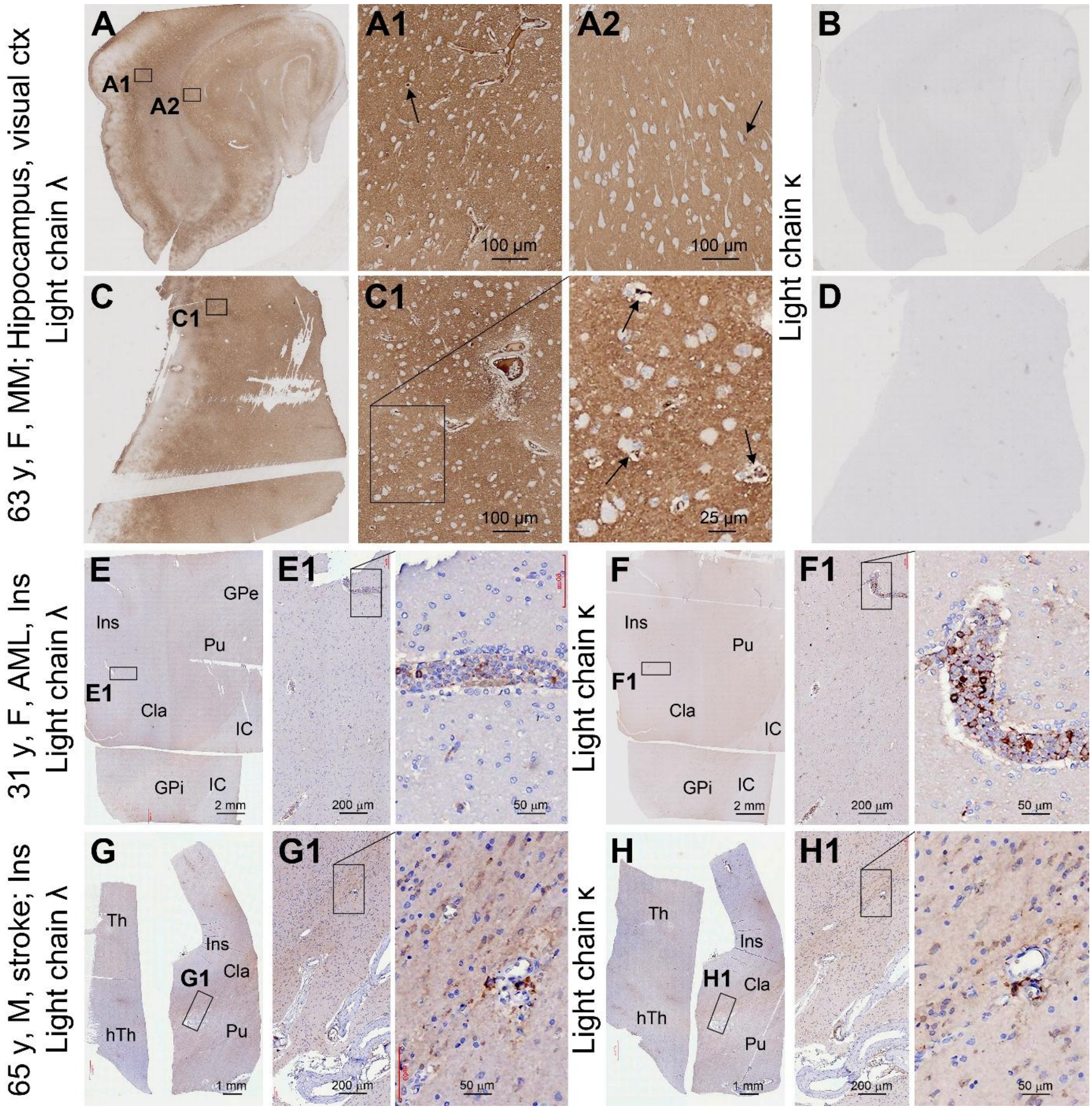


**Supplemental Figure 39. Additional characterization of immunoglobin light chain infiltration in the brains of the 63 y donor with multiple myeloma along with assay controls using other brain samples.** Panels (**A, B, C, D**) show immunolabeling of the two light chains in the temporal lobe (**A, B**) and visual cortical (**C, D**) sections. Diffuse light chain λ infusion in the extracellular space is present in the grey and white matter. Neuronal somata and apical dendrites are devoid of labeling in general; however, a few of them are labeled and appear severely shrunken (**A1, A2, C1,** pointed by arrows). On the contrary, no labeling for the light chain κ is seen in the consecutive sections of these regions (**B, D**). The immunolabeling of both the λ and κ light chains are faint in the insular (Ins) level sections from the 31 y case of acute myeloid leukemia (with cancer cell infiltration), with labeling present inside blood vessels representing the presence of these immunoglobulins in plasm (**E, E1, F, F1**). In the insular level sections from the 65 y case of chronic cerebral ischemia, the labeling for both light chains are background-like and comparable to each other in intensity. The intensity of extracellular labeling appears to be increased in the striatum relative to the insular cortex, lightly suggestive of a leakage of plasma components into the brain because of the vascular injury in the basal ganglia in this case (**G, G1, H, H, H1**).

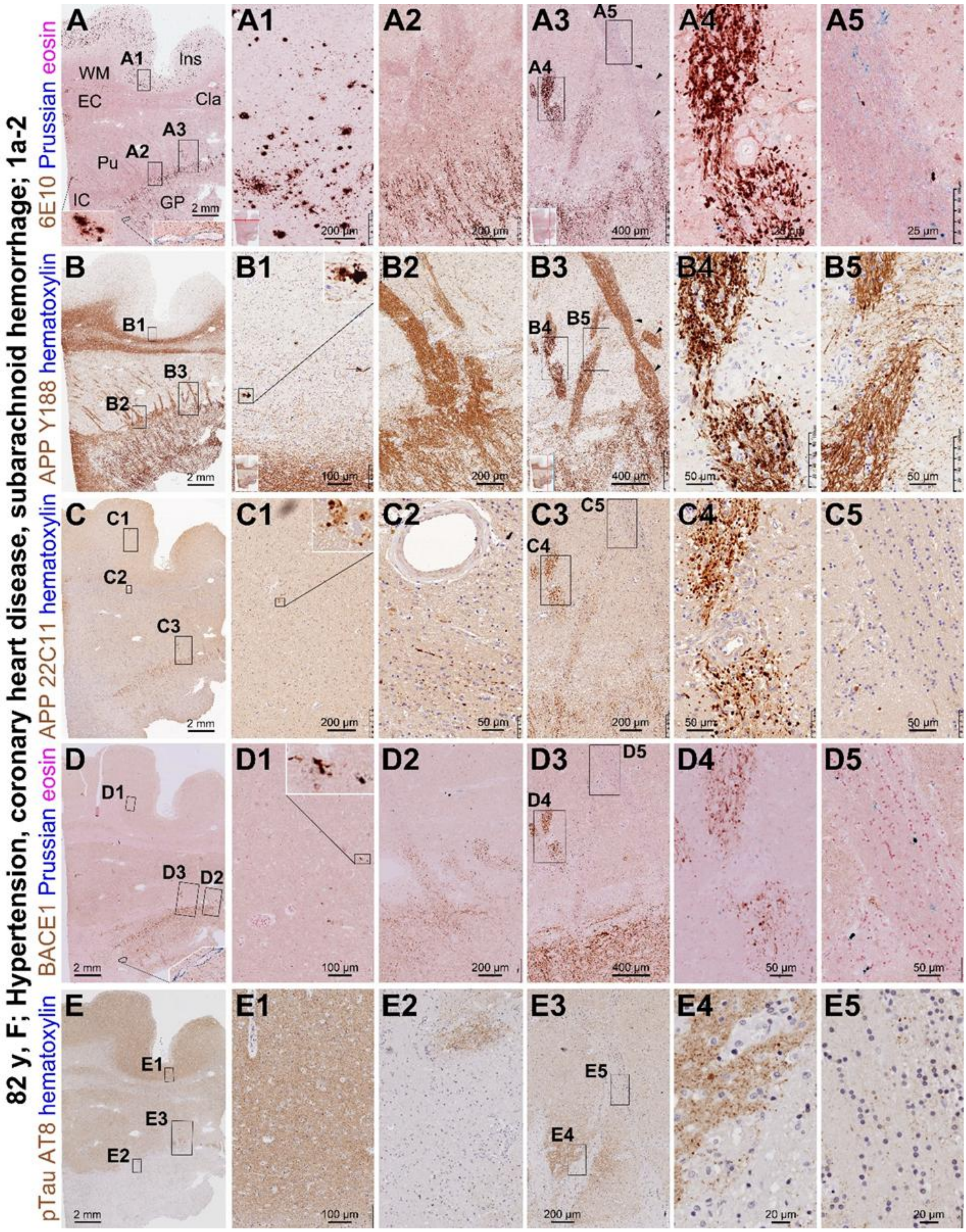


**Supplemental Figure 40**. **Aβ and axonal pathology in insular level sections in the 82 y case with cardiovascular disease**. The 6E10 reveals extensive diffuse Aβ plaques in the insular cortex with a few in the putamen (**A, A1**), and swollen axons in the striatum (**A, A2-A5**). The Y188 antibody displays a few neuritic clusters in the cortex (**B, B1**), with light labeling of the normal axonal bundles and heavy labeling of pathological axonal processes in the striatum (**B2, B5**). The dystrophic neurites and pathological axonal processes are also labeled by the APP 22C11 (**C, C1-5**) and BACE1 (**D, D1-D5**) antibodies. Iron deposits exist in the striatal axonal bundles (**A5, D5**). AT8 labeling appears normal-looking (**E, E1-5**).
